# Heterochronic myeloid cell replacement reveals the local brain environment as key driver of microglia aging

**DOI:** 10.1101/2025.09.02.673859

**Authors:** Claire Gizowski, Galina Popova, Heather Shin, Marius M Mader, Wendy Craft, Wenjun Kong, Yohei Shibuya, Bernd J Wranik, Yuheng C Fu, Constanze Depp, Tzuhua D Lin, Baby Martin-McNulty, Yongjin Yoo, Po-Han Tai, Maximilian Hingerl, Kayla Leung, Micaiah Atkins, Nicole Fong, Devyani Jogran, Agnieszka Wendorff, David Hendrickson, Astrid Gillich, Andy Chang, Beth Stevens, Marius Wernig, Oliver Hahn

## Abstract

Aging, the key risk factor for cognitive decline, impacts the brain in a region-specific manner, with microglia among the most affected cell types. However, it remains unclear whether this is intrinsically mediated or driven by age-related changes in neighboring cells. Here, we describe a scalable, genetically modifiable system for in vivo heterochronic myeloid cell replacement. We find reconstituted myeloid cells adopt region-specific transcriptional, morphological and tiling profiles characteristic of resident microglia. Young donor cells in aged brains rapidly acquired aging phenotypes, particularly in the cerebellum, while old cells in young brains adopted youthful profiles. We identified STAT1-mediated signaling as one axis controlling microglia aging, as STAT1-loss prevented aging trajectories in reconstituted cells. Spatial transcriptomics combined with cell ablation models identified rare natural killer cells as necessary drivers of interferon signaling in aged microglia. These findings establish the local environment, rather than cell-autonomous programming, as a primary driver of microglia aging phenotypes.

## Introduction

Aging is the primary risk factor for cognitive impairment^1–3^, neuroinflammation^4^ and various neurodegenerative disorders such as Alzheimer’s disease^5, 6^. How aging influences the onset and progression of these distinct neuroinflammatory and neurological conditions, however, remains unclear. Recent molecular mapping efforts using epigenetic, transcriptomic, and metabolomic profiling across diverse cell types and regions of the aging brain in mice and humans consistently reveal that glia show earlier and more pronounced age-related changes compared to neuronal cell types^7–14^. Among glial cells, microglia, the resident immune cells of the central nervous system (CNS)^15^, have invariably emerged as one of the most affected cell type in the aging brain^7, 16, 17^. Microglia from the same individual exhibit highly variable age-related changes (referred from here as ‘aging trajectories’), notably an induced interferon response and expression of ‘disease-associated microglia’ (DAM) genes (*Gpnmb*, *Spp1*, *Itgax*, *Apoe,* etc.)^18^, which depend on the brain region and local cellular neighborhood^7, 8, 12, 19^. Specifically, microglia in white matter-rich regions and the cerebellum exhibit these accelerated aging trajectories^7, 8^. However, whether these region-specific changes arise from cell-intrinsic mechanisms such as impaired DNA integrity, mitochondrial damage, or microglia exhaustion ^20, 21^, or are a consequence of local, cell-extrinsic signals like myelin debris, infiltrating T cells, or age-related changes in other cell types ^12, 16, 22, 23^, is unclear.

Identifying the underlying molecular mechanisms that drive microglia aging has been particularly challenging. It requires years to breed and age transgenic lines for genetic perturbations, which imposes a significant limitation on efficient and iterative experimentation. Further, *in vivo* viral transduction of microglia continues to pose significant challenges^24^, as current viral vectors lack both efficiency and cell-type specificity^25^, and can elicit a substantial interferon response^26^, thereby making it difficult to distinguish age-related increases in neuroinflammation from that induced by the transduction method itself. In contrast, myeloid cell replacement strategies, in which microglia are replaced via a combination of bone marrow transplantation and administration of colony-stimulating factor 1 receptor (CSF1R) inhibitors (Plexxikon; PLX) ^27, 28^, or direct intracranial injection of primary neonatal microglia in PLX-treated brains^29^ provide valuable alternatives. However, these approaches are limited by the low number of hematopoietic stem cells (HSCs) in bone marrow, and the difficulty of genetically editing primary microglia in vitro, respectively^17^. To our knowledge, there are no comprehensive, scalable experimental strategies for single or pooled genetic manipulations of myeloid cells in aged brains that can definitively distinguish intrinsic from extrinsic drivers of age-related changes. Here, we have adapted and developed a modifiable system using HSCs for brain myeloid cell replacement to study the effects of extrinsic aging on microglial function.

## Results

### Scalable CNS Myeloid Cell Replacement Workflow in Young and Aged Mice

To probe for the spatiotemporal molecular changes of microglia across the mammalian brain at young and old age, we performed paired single-cell transcriptomics and mapping of 105 surface proteins (scRNA-seq + CITE-seq) of CD45+ immune cells derived from cortex and cerebellum of young and aged wildtype mice (WT; C57BL/6J strain; Figures 1A,B and S1A). Given that age-related molecular changes accumulate more rapidly in the cerebellum than in the cortex^7, 19^, we focused our analyses on these regions as they provide an informative contrast. We utilized both transcriptome and surface-proteome data to identify microglia and their states (labeling adopted from Fumagalli et al.^30^), as well as clusters of border-associated macrophages, peripheral macrophages, granulocytes, T cells and a small population of natural killer cells (NK cells; Figures S1B-L; Table S1). Consistent with previous microglia bulk transcriptomic studies (Figures S1M-P)^19, 31^, our single-cell data revealed extensive differential gene expression patterns between cerebellar and cortical microglia at young baseline (Figure 1C,D). Additionally, we observed corresponding shifts in several surface proteins (*Ptprc*/CD45, CD34, PECAM1, etc.) confirming that region-specific mRNA expression changes are reflected at the protein level (Figures 1C and S1H; Table S1). In contrast to cortical microglia, cerebellar microglia showed elevated expression of antigen-presentation and pro-inflammatory genes, indicative of a low-grade interferon response (Figure 1C,D; Table S1), as previously described^19^. The data can be interactively explored at https://tinyurl.com/Microglial-Aging-Explorer.

**Figure 1.**
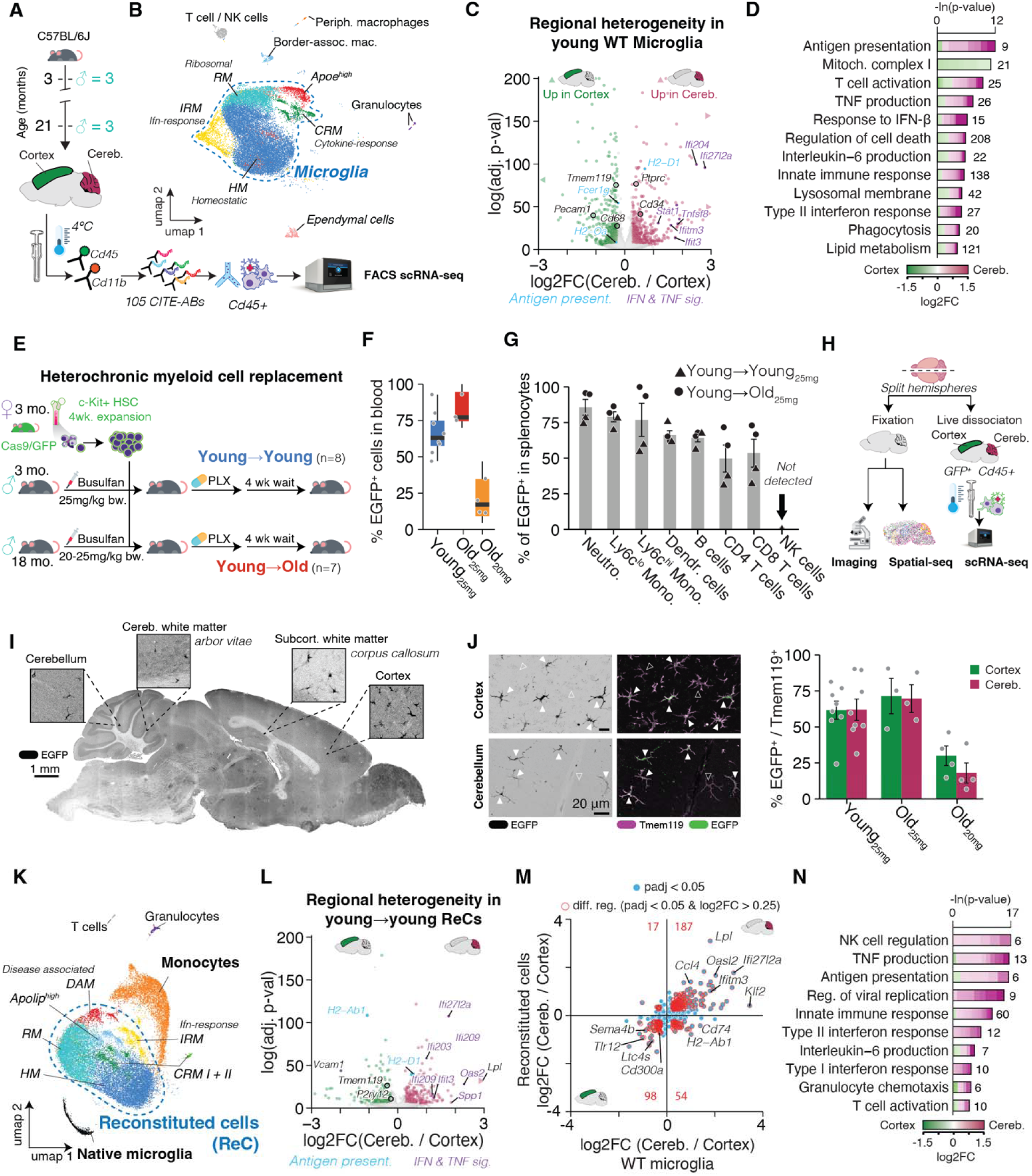
Ex vivo expansion of hematopoietic progenitors enables scalable regional and heterochronic myeloid replacement. **(A)** Experimental design: FACS isolation of CD45+ cells from cortex and cerebellum of 3- and 21-month-old male C57BL/6J mice (tissues from n = 3 animals per age group were pooled) profiled by CITE-seq and scRNA-seq. **(B)** UMAP representation of CD45+ cells: homeostatic microglia (HM), interferon response microglia (IRM), ribosomal processing microglia (RM), cytokine response microglia (CRM), Apoe^high^ microglia, border-associated macrophages, T/NK cells, granulocytes and ependymal cells. Cell composition data is provided in Figure S3I. **(C)** Volcano plot of differential gene expression in young WT microglia (cerebellum vs cortex). Key genes in antigen-presentation, interferon and tumor necrosis factor-signaling (IFN; TNF) are highlighted. **(D)** Representative gene ontology (GO) analysis for region-specific DEGs: lengths of bars represent negative ln-transformed padj using *Fisher’s exact test*. Colors indicate gene-wise log2FC(cerebellum/cortex). Numbers indicate DEG counts per term. **(E)** Schematic of heterochronic myeloid replacement: Cas9/GFP+ c-Kit+ HSCs were expanded ex vivo and transplanted into busulfan-conditioned young (3 mo) or aged (18 mo) recipients. Young→young (blue) recipients received busulfan at dosages of 25 mg/kg bodyweight (bw.), while young→old (red) recipients received busulfan at dosages of 20 mg or 25 mg/kg bw. See Figure S2 for details. **(F)** % EGFP+ cells in peripheral blood of recipients; resolved by age and busulfan dose. **(G)** % EGFP+ in splenocytes of young (▴) and aged recipients (●). Mean ± SEM. **(H)** Experimental workflow post-reconstitution: One hemisphere was fixed for imaging and CosMx spatial transcriptomics, the other hemisphere was freshly isolated and mechanically dissociated on ice for scRNA-seq of GFP+CD45+ cells. **(I)** Representative sagittal section showing EGFP+ cells. **(J)** Representative images and quantification of EGFP and Tmem119; filled arrows indicate EGFP+Tmem119+ cells; empty arrows EGFP^-^, native myeloid cells. Mean ± SEM. n = 4-6/group. **(K)** UMAP of reconstituted GFP+ cells (ReC) : Homeostatic myeloid cells (HM), interferon response myeloid cells (IRM), ribosomal processing myeloid cells (RM), cytokine response myeloid cells (CRM), Apoe^high^ cells, Disease-associated myeloid cells (DAM). A subset of native microglia was captured. Cell composition data is provided in Figure S3J. **(L)** Regional DEGs in young ReC (cerebellum vs cortex). **(M)** Scatter plot comparing log2FC in ReC (y-axis) versus WT microglia (x-axis); red, genes with padj < 0.05 and |log2FC| > 0.25; numbers indicate counts of region-biased genes per quadrant. **(N)** GO-term enrichment for 285 regional DEGs found in ReC and WT microglia (format as in D). Complete DGE and GO tables are listed in Table S1.

Though there is a growing body of evidence demonstrating that environmental factors contribute to microglial regional heterogeneity, the degree to which this specialization is hard-wired in resident microglia remains unclear^31–34^. We sought to determine whether these signalling factors exclusively direct regional specification on resident microglia or if they can also induce differentiation of myeloid cells of different origins. To probe our hypothesis and gain a deeper understanding of how aging trajectories arise in microglia, we adapted and developed a robust and scalable workflow to repopulate the CNS of young and aged WT recipient animals with myeloid cells from intrinsically young donor cells (Figures 1E and S2A). We based our protocol on previously published methods^27, 28^ using busulfan-based chemotherapeutic peripheral bone marrow conditioning followed by transient administration of a CSF1R inhibitor (PLX5662; referred to as PLX) to achieve a near-complete and homogeneous replacement of brain-resident microglia with circulation-derived myeloid cells. To increase replicability and circumvent the use of fresh bone marrow that contains low amounts of HSCs, we adopted an in vitro expansion protocol to cultivate a pool of 100M hematopoietic progenitor cells^35^ that have CRISPR gene editing capabilities and are fluorescent by isolating c-KIT+ cells from female Rosa26-CAG::Cas9-EGFP knock-in mice^36^. Gene expression profiles were characterized with scRNA-seq which revealed the resulting cell pool to consist of various progenitor populations and hematopoietic stem cells (Figures S2B-D). We refer to this cell pool for brevity as ‘expanded HSCs’ (eHSCs).

Young and aged WT mice (3 and 18 months-old, respectively) exhibited robust reconstitution of the peripheral myeloid (75-100%) and lymphoid lineage (50-70%) with donor-derived cells (Figure 1F,G). Consistent with previous reports, we observed no reconstitution of NK cells, likely due to their resilience to chemical or radiation-based ablation^37^. Immunofluorescence and confocal microscopy analyses confirmed brain repopulation by EGFP+ cells co-expressing the microglia marker TMEM119 (Figures 1H-J). Further validation by fluorescence activated cell sorting (FACS) confirmed the presence of EGFP^+^ immune cells, predominantly of the myeloid (CD11b+) lineage, throughout the brain (Figure S2E). Peripheral and brain-specific chimerism were significantly correlated, and busulfan treatment produced a dose-dependent increase in chimerism (Figures 1H-J, S2E and S3A). Together, our workflow utilizes the scalable nature of eHSC expansion to reconstitute the peripheral and brain-specific immune system of young and aged mice.

### Reconstituted Myeloid Cells Adopt Region-Specific Microglial Identities

To investigate the acute impact of the brain environment on myeloid cells, we conducted scRNA-seq of EGFP+/CD45+ cells isolated from the cortex or cerebellum of young mice after CNS myeloid cell replacement. After filtering low-count and low-quality cell clusters, we confirmed expression of both the Cas9-EGFP and the female-specific *Xist* transcripts in most scRNA-seq clusters with the exception of a small population of recipient-derived (native) microglia and ependymal cells (Figures 1K and S3A-E). Consistent with previous CNS myeloid cell replacement strategies, chimeric myeloid cells in the brain exhibited expression of markers like *Ms4a7*^38, 39^, but lacked expression of microglia transcription factors *Sall1* and *Sall3* (Figure S3E)^27, 38^. This indicates that chimeric myeloid cells retain markers of their peripheral ontogeny. We observed two clusters that we labelled as ‘monocytes’, harboring high expression levels of peripheral myeloid cell marker genes *Cd74*, *H2-Aa* and *H2-Ab1*. Pseudotime analysis indicated that these monocytes were on one end of a bifurcated trajectory that was marked by gradual down-regulation of these marker genes while adopting a set of transcriptional clusters akin to those observed in WT microglia (Figure S3F-K; Table S1), including interferon-response (IRM), cytokine-response (CRM) and disease associated-like (DAM) myeloid clusters. We classified DAMs in ReCs based on the expression of *Spp1*, and *Gpnmb*^30^. Thus, while not derived from the same developmental trajectory as microglia, chimeric myeloid cells adopt a defined set of microglia-like molecular states, indicating that these cells have adapted to the CNS environment. We refer to these chimeric, donor-derived myeloid cells with microglia-like clusters as ‘Reconstituted Cells’ (ReCs).

We next asked whether the distinct transcriptional profiles of region-specific microglia could be replicated in ReCs. To this end, we focused on differentially expressed genes (DEGs) found in cerebellar versus cortical WT microglia, including *Lpl*, *Ccl4* and *Sema4b* (Figures 1C and 1L). Differential expression analysis between cerebellar and cortical ReCs identified a set of 285 region-specific DEGs that significantly overlapped with those in WT microglia (Figure 1L-N; Table S1). We then utilized this set of 285 genes to generate a transcriptional “signature”^7, 40^ of cerebellar microglia, which included several antigen presentation and interferon signaling genes (Figure 1N). The resulting signature score was significantly elevated in ReCs, monocytes and native microglia - which were not used to train this signature - isolated from the cerebellum of mice at any age, which was well-replicated across animals (n=6-7; Figure S3L). We subsequently use this set as a regional identifier to distinguish cerebellar from cortical WT microglia and ReCs.

To corroborate these region-specific expression patterns with independent methods, we performed spatial transcriptomics (CosMx platform; referred to as Spatial-seq) using a molecular imaging platform on fixed-frozen sagittal brain sections from young and aged reconstituted mice (n=2 per age; >80% chimerism). We integrated, clustered and labelled the resulting 569,481 single-cell transcriptomes, before co-integrating a cluster of likely immune cells with the high-coverage scRNA-seq data from cortex and cerebellum to enable accurate identification of ReCs and their states (Figures 2A-C and S4A-G). Notably, due to the absence of interferon response genes in the default CosMx panel, we could not identify IRMs. ReCs located in the cerebellum exhibited elevated cerebellar microglia signature scores relative to those in other brain regions (Figure 2D), indicating that ReCs also exhibit cerebellar signatures *in situ*. We observed widespread distribution of ReCs across the brain, along with region-specific differences in their density and tiling patterns (Figure 2E). For instance, the cerebellum was less densely repopulated with ReCs compared to the cortex, and within the cerebellum, ReCs were predominantly localized in the arbor vitae (white matter of the cerebellum) and the granule cell layer, while largely absent from the molecular layer. This unique tiling behavior has been previously described in WT microglia (Figure 2E-G)^34, 41, 42^, suggesting that ReCs adopt region-specific microglial phenotypes beyond transcriptional shifts.

**Figure 2.**
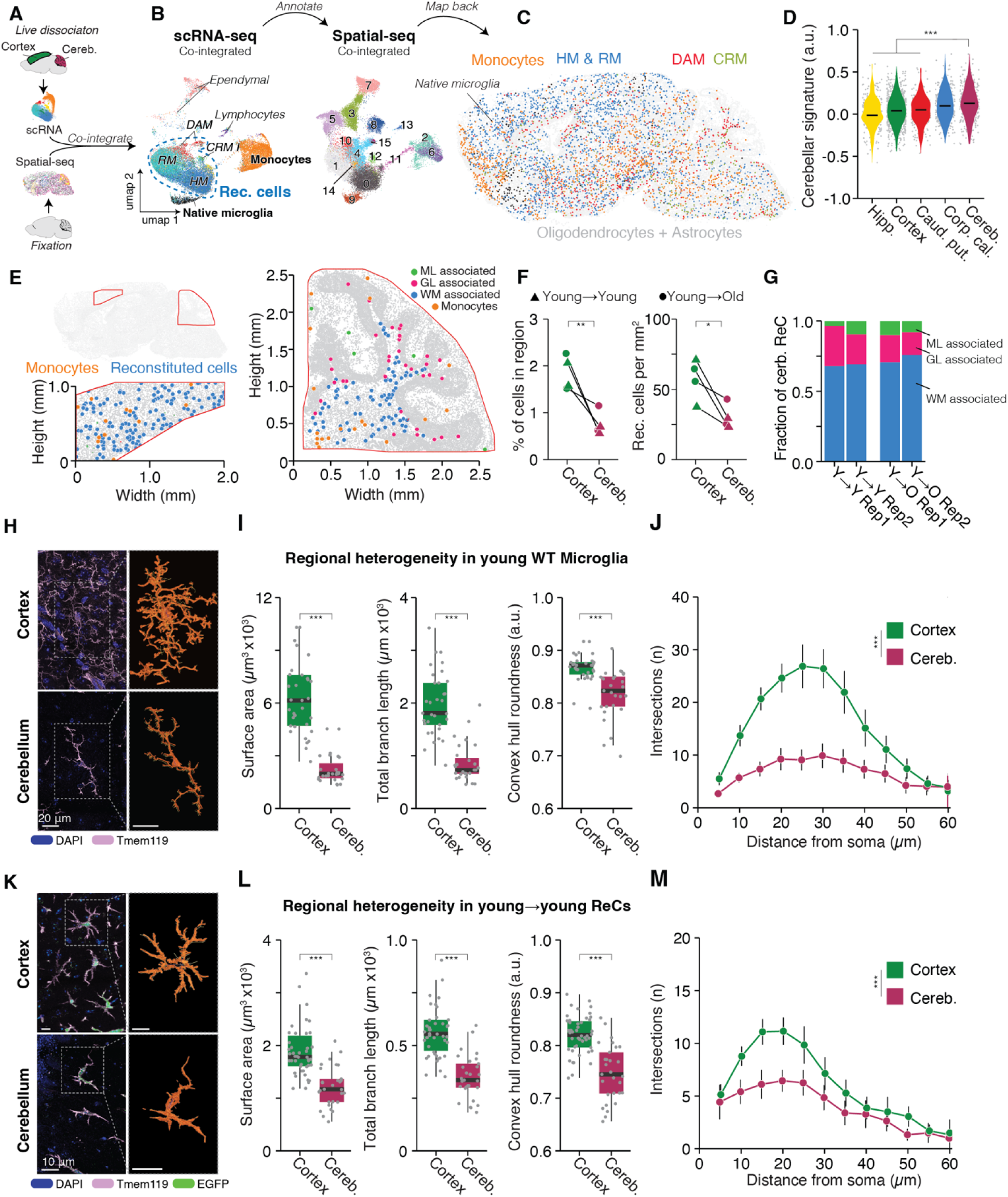
Reconstituted cells adopt region-specific tiling and morphology profiles. **(A)** Analysis workflow: annotated, whole transcriptome scRNA-seq of live GFP+CD45+ cells (see Figure 1K) was co-integrated with immune cells identified in CosMx data. **(B)** UMAP embedding of co-integrated scRNA-seq (left) and spatial-seq (right) colored by annotated clusters (cluster IDs as shown). **(C)** Spatial distribution of GFP+ ReC populations across representative CosMx data, overlaid on oligodendrocytes and astrocytes (grey). **(D)** Violin plots of cerebellar signature (a.u.) across hippocampus (Hipp.), cortex, caudate-putamen (Caud.-Put.), corpus callosum (Corp. Cal.) and cerebellum (Cereb.); ***p < 0.001 by two-sided *Wilcoxon test*. **(E)** Spatial positions of ReCs in representative sections of cortex and cerebellum. ReCs annotated to cerebellar molecular layer (ML, green), granular layer (GL, cyan) and white matter (WM, magenta) are indicated. **(F)** Left, fraction of ReCs in cortex and cerebellum of young→young (▴) and young→old (●) chimeras. Right, density of reconstituted cells per mm². Paired two-tailed *t test*.; **p < 0.01. **(G)** Relative fraction of cerebellar ReCs in ML, GL and WM. **(H)** Representative confocal images of Tmem119 and DAPI in cortex and cerebellum with skeletonized and traced morphology (orange). **(I)** Microglial surface area (×10³ µm²), total branch length (×10³ µm) and convex hull roundness in cortex and cerebellum of young mice; ***p < 0.001 by two-tailed *t test*. **(J)** Sholl profile of microglial intersections versus distance from soma in the cerebellar molecular and granule cell layer and cortex (n = 3 mice per group, 12 to 20 cells per animal and region were quantified). Two-sided *Wilcoxon rank-sum tests* on area under the curve AUC. Mean ± SEM; ***p < 0.001. **(K)** Same as (H) for ReCs. **(L)** Same as (I) for ReCs **(M)** Same as (J) for ReCs.

To further validate these findings, we performed confocal imaging and quantitative morphological analysis of WT microglia and ReCs in the cerebellum and cortex. Consistent with previous reports^31, 34^, cerebellar microglia in the molecular and granule cell layers exhibited a distinctive ‘rod-like’ morphology, characterized by significantly reduced surface area, branch length, roundness, and complexity, as measured by Sholl analysis, compared to cortical microglia (Figure 2H-J). In comparison to WT microglia, ReCs exhibited a compact morphological profile, with an overall smaller size and significantly reduced branching structure. Nonetheless, ReCs preserved region-specific morphological features. ReCs in the cerebellum showed significantly decreased surface area, branch length, roundness, and branching relative to ReCs in the cortex, mirroring the relative morphological distinctions observed in endogenous microglia from these brain regions (Figure 2K–M). In summary, our eHSC-based workflow results in repopulation of the brain with peripherally-derived myeloid cells that adopt microglia-like transcriptional states, as well as region-specific molecular, morphological and distribution features of microglia.

### Exposure to the Aged Brain Environment Triggers Age-Related Molecular and Morphological Changes in Intrinsically Young ReCs

The robust and rapid adoption of region-specific features by ReCs supports the notion that distinct CNS regions act as key modulators of myeloid cell biology^31, 34^. We therefore wondered whether ReCs derived from young donors would retain the molecular features of young cells in aged recipient brains or, instead, adopt age-related characteristics. We first confirmed that cerebellar WT microglia exhibit more DEGs with aging (padj<0.05, |log2FC| > 0.25 between old and young cells; referred to as ‘age-related DEG’) than in the cortex (Figure 3A,B). Of the 730 DEGs that were significantly differentially expressed in both the cortex and cerebellum, many showed stronger regulation in cerebellar microglia, indicating more pronounced expression changes in the cerebellum relative to the cortex. To validate these findings at the protein level, we examined our CITE-seq dataset and confirmed that several of these genes also showed higher protein expression in cerebellar microglia (Figure 3B). This included, among others, the phagocytosis inhibitor *Cd22*/CD22^43^, DAM marker *Itgax*/CD11c^44^, cell adhesion regulator *Vcam1*/CD223^45^, and interferon response genes *Bst2*/CD317 and *Ly6a*/Sca-1^46^ (Figures 3A,B, S5A; Table S2). While microglia in both the cortex and cerebellum of aged brains showed activation of interferon signaling, shifts in lipid metabolism and induction of antigen presentation genes, interferon response genes were more prominently regulated in cerebellar microglia. Additionally, a mild upregulation of most ribosomal proteins was observed in the cerebellum but not in the cortex (Figures 3C and S5B; Table S2).

**Figure 3.**
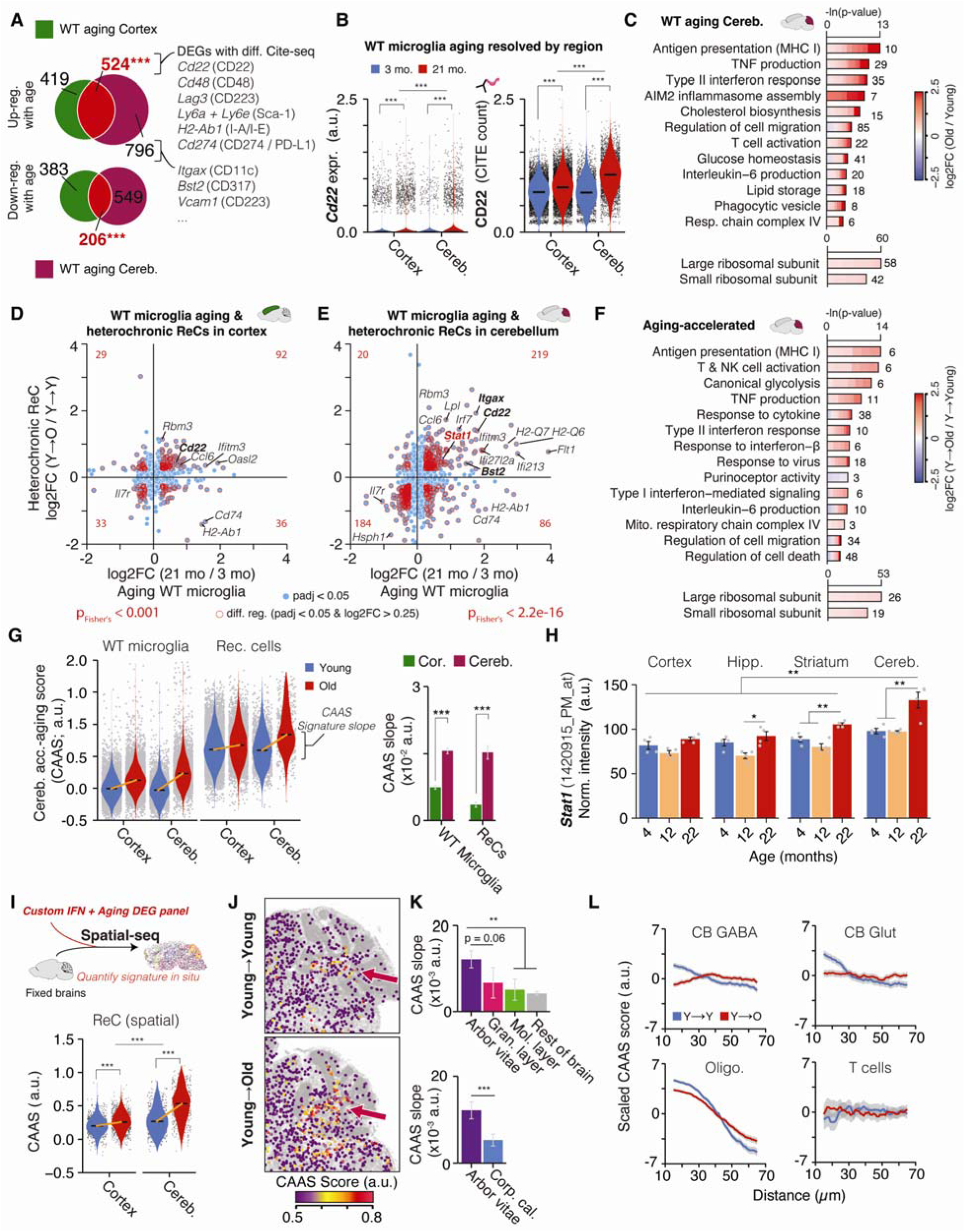
The aged brain environment induces age-related expression patterns in intrinsically-young reconstituted cells. **(A)** Overlap of DEGs up-regulated (top) or down-regulated (bottom) with age in WT microglia from cortex (green) versus cerebellum (magenta); numbers indicate DEG counts. Significance of overlap was evaluated by Fisher’s exact test. ***p < 0.001. Representative genes with matching differential surface protein abundance as measured by CITE-seq are listed. **(B)** Violin plots of *Cd22* RNA expression (left, a.u.) and corresponding protein CITE count (right) in aged microglia. Median is indicated. *MAST*, Benjamini-Hochberg correction. ***padj < 0.001 **(C)** Representative GO-term enrichment for age-related DEGs in cerebellar microglia. **(D, E)** Scatter plots comparing per-gene log2FC(21 mo/3 mo) in WT microglia (x-axis) versus heterochronic ReCs (y-axis) in cortex (D) and cerebellum (E). Only genes with padj<0.05 are shown. DEGs (padj < 0.05 & |log2FC| > 0.25) found both cell types are indicated in red. The number of overlapping DEGs in each quadrant is indicated in red. **(F)** GO-term enrichment for 403 aging-accelerated DEGs in ReCs. **(G)** Left, distributions of cerebellar-acceleration aging score (CAAS) in WT microglia and ReC from cortex and cerebellum. Medians are indicated. Right, CAAS slope of linear regressions for each group. Mean ± 95% confidence intervals. Two-sided *Tukey’s HSD test*. The highest (least significant) p value is indicated. ***p < 0.001. **(H)** Quantification of *Stat1* (1420915_PM_at) in microglia bulk RNA microarray data across regions and aging from^19^. (n = 4/group). Statistical significance was assessed using limma’s *moderated t test* with FDR correction. Mean ± SD. *p < 0.05, **p < 0.01. **(I)** Top, schematic for Spatial-seq of young and aged chimeric brains using CosMx neuropanel and a custom 40 gene panel covering interferon response and CAAS genes. Custom panel listed in Table S2. Bottom, Violin plots of in-situ quantified CAAS in ReCs for cortex and cerebellum. ***p < 0.001 by two-sided Wilcoxon test. Quantification and statistical analysis of CAAS slope can be found in Figure S7. **(J)** Spatial maps of CAAS (color scale) over cerebellar sections from young→young and young→old chimeras; red arrows indicate area of increased CAAS with age. **(K)** Quantification of CAAS slope in indicated cerebellar subregions (top) and across white matter tracts (bottom). Mean ± 95% confidence intervals. Two-sided *Tukey’s HSD test*. The highest (least significant) p value is indicated. *p < 0.05, **p < 0.01. **(L)** Distance-dependent CAAS profiles for ReCs relative to cerebellar (CB) GABAergic neurons, CB glutamatergic neurons (CB Glut), oligodendrocytes (Oligo.) and T cells. A constant that equals to the mean score across all distances is subtracted from each curve. Data are presented as mean (solid line) ± SEM (shade). Complete DGE and GO tables are listed in Table S2.

We next asked whether young ReCs, when introduced into aged brains, would adopt similar age-related expression shifts or retain molecular features of young cells (’young→old’ vs ‘young→young’; referred to as heterochronic reconstitution). Focusing on shared DEGs found both in aged WT microglia and in ReCs in old brains (passing significance cutoff and log2FC threshold of 0.25 in both groups), we observed a highly significant overlap of DEGs in both cortex and cerebellum. In total, we observed 125 overlapping DEGs in the cortex and 403 DEGs in the cerebellum, with 49 genes regulated in myeloid cells across regions (Figure 3D,E; Table S2). While the 403 overlapping genes represent only ∼20% of all age-related DEGs found in WT microglia, this gene set still exhibited strong functional enrichment for heightened interferon signaling, ribosomal protein induction and up-regulation of the antigen presentation machinery (Figures 3F; Table S2). We observed a similar, albeit weaker, functional enrichment for overlapping DEGs in the cortex (Figure S5C; Table S2). Thus, a functionally relevant subset of the major gene expression changes observed in microglia of aged WT mice is also induced in young ReCs following reconstitution into aged brains.

Beyond gene expression changes, microglia in aged brains also undergo morphological shifts, becoming less ramified and more dystrophic^17, 47, 48^. To determine if ReCs exhibit similar age-related morphological changes, we quantified the morphological features of WT microglia and ReCs in the cortex and cerebellum (molecular and granule cell layer) of young and aged mice. Aged cortical WT microglia displayed significantly reduced surface area, shorter branch length, and decreased branching (Figure S5D,E). Similarly, cortical ReCs in aged brains showed significant reductions across these same morphological metrics. Interestingly, neither WT microglia nor ReCs in the grey matter of the cerebellum (i.e. molecular and granule layer) exhibited age-related morphological changes, which may be due to their inherent rod-like shape, which is present even at a young age (Figure S5F,G). These findings demonstrate that local cues within the CNS acutely drive region-specific, aging-like changes in both gene expression and morphology in ReCs.

### Magnitude of Age-Related Expression Shifts in ReCs Depends on Brain Region

We combined the 403 DEGs showing matching age-related expression in cerebellar WT microglia and ReCs (Figure 3E,F; Table S2) into a signature, herein referred to as the *cerebellar accelerated aging signature* (CAAS). The CAAS was more strongly induced with aging in cerebellar than cortical WT microglia, and this region-specific pattern was recapitulated in ReCs, despite their overall higher baseline signature score (Figure 3G). Native microglia in the cerebellum of aged reconstituted mice also displayed CAAS induction near-identical to what we observed in ReCs (Figure S6A). These robust effects were reproducible across independent biological samples, regardless of brain chimerism percentage (Figure S6B), and were equally encoded by all ReCs and microglia clusters (Figure S6C-D). While we observed an upregulation of the DAM-like marker genes *Gpnmb* and *Spp1*^44^ in cerebellar ReCs, expression of these genes in WT cerebellar microglia was too low for detection as DEGs, and thus were not included in the CAAS. However, meta-analysis of bulk-isolated microglia^19^ confirmed strong up-regulation of DAM-like genes *Itgax*, *Gpnmb* and *Spp1* in the aged cerebellum (Figure S6E-G), indicating that aged cerebellar microglia adopt a DAM-like expression profile, one that is likewise rapidly induced in ReCs in aged brains. Thus, the well-documented, region-specific variation in expression of key age-related microglia genes, including *Stat1*, *Itgax*, *Gpnmb*, *Spp1* and *Cd22*^7, 8, 12, 19^, is likely driven by the local brain environment (Figures 3H and S6E-G).

To gain deeper spatial insights into the accelerated aging effects in ReCs across the whole brain, we performed Spatial-seq on sagittal sections with a custom-built 40 gene panel that included several CAAS genes as well as interferon response genes (Figures 3I-J and S7A-E; Table S2). ReCs in the aged cerebellum exhibited the strongest CAAS increase compared to other brain regions *in situ*, including the cortex, thalamus and striatum (S7G-H). Within the cerebellum, ReCs in the arbor vitae exhibited the strongest CAAS increase compared to those in the granule cell or molecular layer, and this increase was more pronounced than in other white matter regions such as the corpus callosum (Figures 3K,L and S7F-G). Brain-wide analyses of cell proximity effects identified cerebellar granule neurons, Purkinje neurons, and oligodendrocytes as candidate drivers of age-related CAAS increases in ReCs, whereas T cells showed no significant effect on CAAS (Figure 3L). Interestingly, while DAM-like genes were induced specifically in the arbor vitae, interferon response genes were induced uniformly across the cerebellum (Figure S7H-K), suggesting these transcriptional programs in the aged cerebellum may be driven by distinct factors.

We next evaluated the CAAS across several published datasets to test the robustness of the signature. Among several brain regions analyzed in a Smart-seq2 dataset^49^, cerebellar microglia exhibited the strongest age-related CAAS increase (Figure S8A-C). Moreover, single-cell and single-nucleus datasets of the aging brain consistently identified microglia as the cell type with the strongest age-related CAAS increase^7, 46^(Figure S8D-I). Although microglia comprise only 5-10% of the total cell population within a given brain region, an increased CAAS score was still detectable in bulk RNA-seq of whole cerebellar tissue^7^ when compared with other brain regions (Figure S8J-L). We also observed a significant overlap of CAAS genes with well-documented transcriptome signatures from aged human microglia^50^ including interferon response genes *Irf7, Bst2*, *Tap1,* and *Psmb8* (Figure S8M). These findings identify the local brain environment as a key driver shaping microglial aging phenotypes and highlight the CAAS as a robust, conserved indicator of these changes.

### Young CNS environment reverses molecular and morphological features of intrinsically old myeloid cells

Given the apparent plasticity of ReCs, we sought to evaluate the adaptability of myeloid cells to different brain environments and the reproducibility of this myeloid cell engraftment approach. To this end, we employed an orthogonal replacement model that enabled reciprocal heterochronic (young→old; old→young) and isochronic (young→young; old→old) myeloid cell replacement without requiring an ex vivo HSC culture step. Using fresh bone marrow harvested from young (2 mo) and in-house aged (20 mo) female UBC::EGFP mice^51^, we performed whole-brain myeloid replacement in young (3 mo) and aged (18 mo) female WT mice (Figure S9A)^27, 52, 53^, which yielded brain myeloid cell chimerisms > 90% in all mice (Figure S9B-D). CD45/CD11b/EGFP+ cells were isolated via FACS and sequenced in bulk from the whole brain (n=30,000 cells/sample). Remarkably, principle component analysis showed clustering by recipient age, independent of the intrinsic age of the donor cells (Figure S9E). Likewise, myeloid cells adopted the DEG profile characteristic of the recipient age such that young→old myeloid cells acquired old→old DEG patterns, whereas old→young cells adopted young→young patterns (Figure S9E–H; Table S2). In total, 759 DEGs exhibited shifts in response to the local brain environment. This gene set contained several CAAS genes (e.g. *Ccl3*, *Cd9*, *Lilbr4a*) as well as DAM-like marker genes (*Gpnbm* and *Spp1*), and was significantly enriched for interferon response genes, cytokine production and monocyte chemotaxis (Figure S9G-I; Table S2). Consistent with the expression shifts, young and aged donor cells engrafted into the aged brain showed greater morphological aging features than those engrafted into the young brain (Figure S9J-M). Together, these findings demonstrate that myeloid cells adopt molecular and structural features of the young or aged CNS environment, independent of transplantation method and donor age, genotype, or sex.

### STAT1 Mediates Age-Related Expression Changes in the Cerebellum

The adoption of aging-like expression patterns in the aged cerebellum indicates that ReCs actively sense their environment. Given that WT microglia and ReCs in the aged cerebellum exhibited a strongly induced interferon response, including *Stat1* upregulation (Figure 3H), we asked whether cell-autonomous suppression of this pathway could prevent the adoption of age-related expression changes. To this end, we reconstituted young and aged mice with eHSCs that were electroporated with either non-targeting control or STAT1-targeting guide RNAs *in vitro* (Control and Stat1^-/-^, respectively; Figure 4A). EGFP+ blood cells post-busulfan of animals receiving Stat1^-/-^ cells exhibited expectedly high deletion of the Stat1 gene (98% KO score, Figure S10A,B), confirming successful engraftment and differentiation in vivo. Stat1^-/-^ and Control cells reconstituted young and aged brains with comparable efficiencies, and displayed region-specific morphology and expression patterns, although we observed fewer Stat1^-/-^ ReCs in the interferon response cluster (Figures 4B-E and S10A-K). In young mice, cortical and cerebellar Stat1^-/-^ ReCs retained significant differences in regional identifier DEGs, suggesting that STAT1 knockout does not disrupt their capacity to sense and adapt to the local brain environment present at young age (Figure S10 J,K). In brains of aged mice, however, Stat1^-/-^ ReCs were strongly protected against the age-related induction of interferon response and antigen presentation genes, while also showing marked dampening of *Cd22* and DAM-like genes such as *Gpnmb*, *Spp1*, *Itgax*, alongside upregulation of the homeostatic marker *P2yr12* (Figures 4F-I and S10L-N; Table S2). Notably, the effects of STAT1 deficiency were present across all myeloid cell states and not restricted to the IRM (Figure S10O,P). Moreover, STAT1 knockout prevented the increased CAAS in cerebellar ReCs and significantly dampened the 49 DEGs affected in aged WT microglia and ReCs in both cortex and cerebellum (Figure 4J,K). Finally, Spatial-seq analysis (Figure S11A-E) confirmed that STAT1 deficiency in ReCs lowered the CAAS *in situ* without affecting their density or spatial distribution patterns (Figures 11F-I). Collectively, these findings identify STAT1 signaling as a key mediator of age-related interferon responses in ReCs and potentially microglia, with an additional role in regulating antigen presentation and DAM-like gene expression during aging.

**Figure 4.**
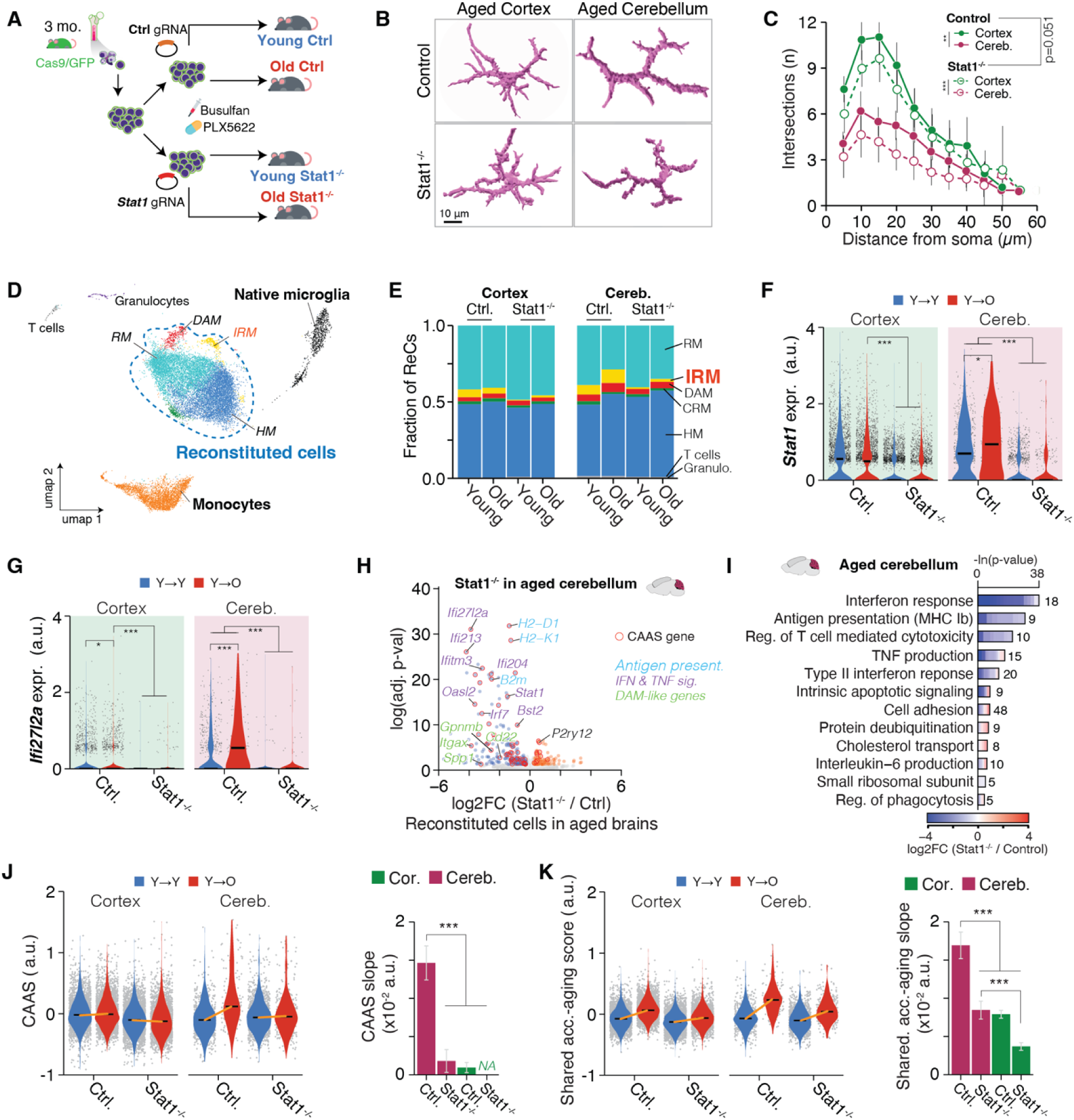
Stat1 deletion abrogates cerebellar-specific aging phenotypes upon heterochronic reconstitution. **(A)** Schematic of heterochronic reconstitution with control (Ctrl) or *Stat1* gRNA–expressing Cas9/GFP+ eHSCs transplanted into young (3 mo) or aged (18 mo) recipients. **(B)** Representative morphological reconstructions of Ctrl and Stat1^-/-^ ReCs in aged recipients. **(C)** Sholl analysis of (B) (n = 3 mice per group, 10 to 15 cells per animal and region were quantified). Two-sided *Wilcoxon rank-sum tests* on area under the curve AUC. Mean ± SEM; *p < 0.05, **p < 0.01, ***p < 0.001. **(D)** UMAP embedding of GFP+CD45+, Ctrl and Stat1^-/-^ cells derived from cortex and cerebellum. Per sample, tissues from two animals per group with similar chimerism and same genotype were pooled. **(E)** ReC cluster composition across regions and genotypes. **(F,G)** *Stat1* (F) and *Ifi27l2a* (G) expression in ReCs from young→young (Y→Y) and young→old (Y→O) chimeras. Median is indicated. MAST, Benjamini-Hochberg correction. ***padj < 0.001. **(H)** Volcano plot of differential expression in Stat1^-/-^ versus Ctrl ReCs from aged cerebellum. CAAS genes highlighted; key IFN-signaling and DAM-like genes labeled. **(I)** Representative GO-term enrichment for DEGs in (H) in aged cerebellum. **(J)** Left, violin plots of CAAS in ReCs from cortex and cerebellum of aged Ctrl and Stat1^-/-^ chimeras. Median is indicated. Right, CAAS slope from linear regression for each group; mean ± 95% CI; two-sided *Tukey’s HSD test*. The highest (least significant) p value is indicated. **(K)** Same as (J) for signature based on 49 accelerated-aging genes found in both cortex and cerebellum.

### Lymphocytes contribute to the interferon response in aged cerebellar microglia

To nominate candidate local signaling networks and factors that may drive the age-related induction of interferon response genes in ReCs and WT microglia, we used CellChat^54^ to analyze cell-cell communication pathways. Previous studies have predicted T cells as key drivers of interferon signaling in microglia during aging^12, 46^, thus we initially focused on CD45+ cells from our scRNA-seq of WT aged mice (Figure 1A,B). Surprisingly, analysis of interaction strength predicted an age-associated increase in type II interferon (IFN-II) signaling between interferon gamma (IFN-γ) secreting NK cells and microglia (Figure 5A-C). Omitting NK cells from the cell pool *in silico* led to the loss of the IFN-II signaling axis in the resulting CellChat network (Figure 5D). To extend our network analysis and identify candidate signaling networks beyond those captured in our CD45+ dataset, we analyzed the Allen Brain scRNA-seq atlas^55^, where we identified NK cells, tanycytes, ependymal cells and T cells as the only significant sources of IFN-γ in the brain (Figures 5E,F, S12A,B). In contrast, potential IFN-β sources were restricted to a small subset of microglia, with no other brain cell types contributing. To better understand how IFNs drive transcriptional changes in microglia, we performed bulk RNA-seq on primary neonatal microglia *in vitro* after stimulation with IFN-γ or IFN-β. Both treatments induced overlapping expression patterns, including type I interferon (IFN-I; *Bst2*, *Isg15*, *Irf7*, *Oasl1)* and IFN-II response genes (*Psme1*, *Psme2*, *Psmb8)*^56^, albeit with differing magnitudes (Figure S12C-I; Table S3), suggesting expression signatures of IFN-I and IFN-II may be largely indistinguishable in microglia in vivo. Notably, only IFN-γ induced DAM-like genes (*Itgax*/Cd11c, *Cst7, Tspo)*. These findings support a model in which lymphocyte-derived IFN-γ amplifies interferon signaling in aged microglia or ReCs.

**Figure 5.**
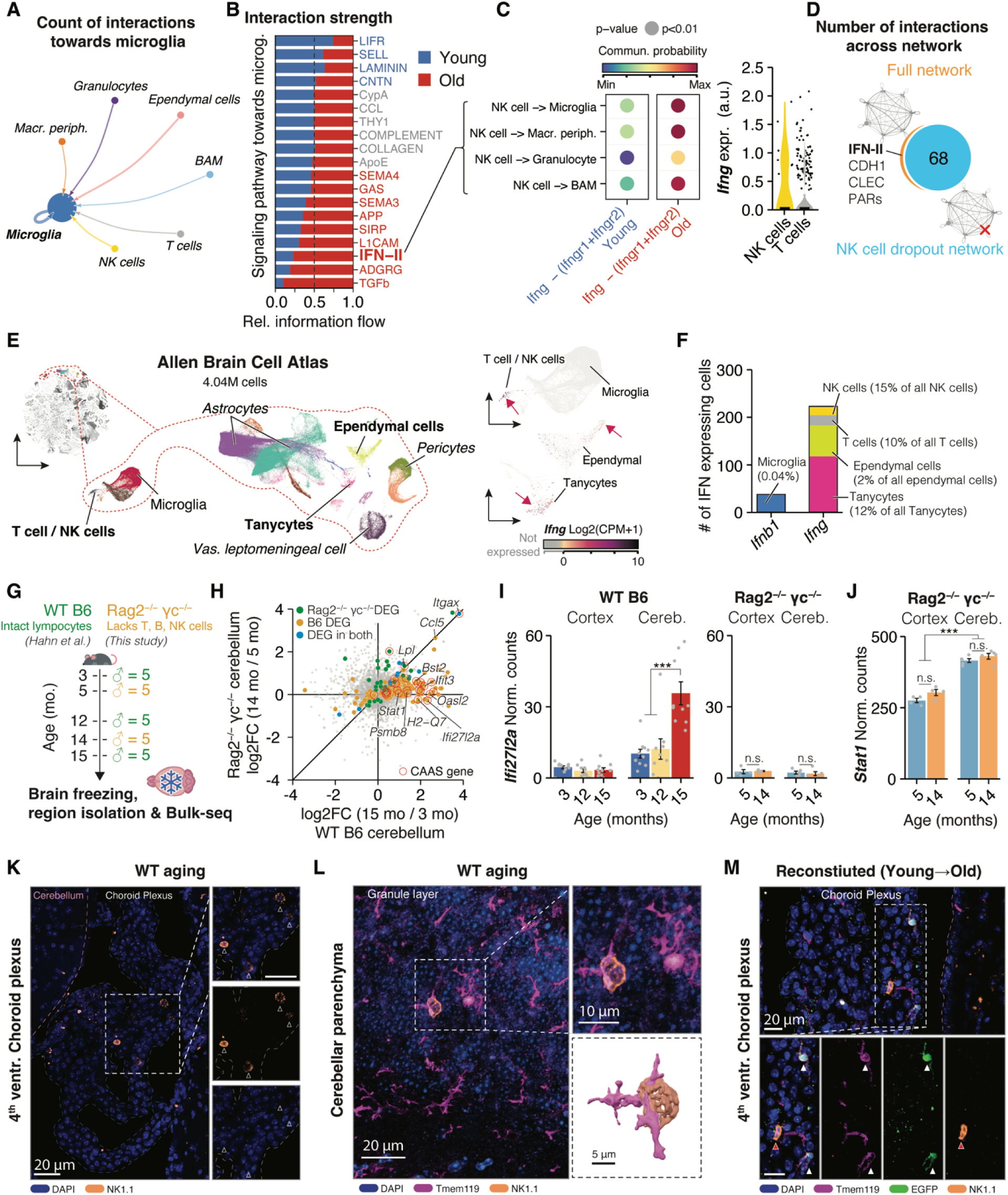
Lymphocyte–mediated interferon signaling in aged microglia. **(A)** CellChat^54^ analysis of cell-to-microglia interactions in young and aged brains based on data in Figure 1B. **(B)** Relative information flow of top signaling pathways into microglia in young (blue) versus aged (red) brains; IFN-II and related ligands (red) dominate in aging. **(C)** Left, circle plots of communication probability and significance for cell-cell interactions identified in ‘IFN-II signaling’. Right, *Ifng* expression in NK and T cells. **(D)** Venn diagram of inferred cell-cell interactions in full versus NK cell–dropout networks; number indicates remaining edges. Four predicted interactions were lost when NK cells were removed from analysis. **(E)** UMAP of cells from the Allen Brain Cell Atlas^87^ with glia populations enlarged, colored by major cell types; inset highlights IFN-expressing populations. **(F)** Counts of cells expressing *Ifnb* or *Ifng* and their percentages of all cells of the same cell type in Allen Brain Cell Atlas. Only cell types with at least 10 cells and 0.01% of all cells of the same type exhibiting IFN expression were considered. **(G)** Cohort overview. Whole brains were collected from 5 and 14 months-old male (n = 5/group) Rag2^-/-^γc^-/-^ animals. Cortex and cerebellum samples were punched from frozen slices and analyzed via bulk RNA-seq. Data was compared to published datasets of C57BL/6JN mice^7^. **(H)** Scatter plot comparing cerebellar log2FC(15 mo/3 mo) in WT C57BL/6JN (x-axis) versus Rag2^-/-^γc^-/-^ (y-axis) bulk RNA-seq; CAAS genes in red. DEGs (padj < 0.1) found in either or both mouse backgrounds are indicated. **(I,J)** Expression of *Ifi27l2a* (I) and *Stat1* (J) across regions, age groups and genotypes. Mean ± SEM. Two-sided *Wald test*. ***p < 0.001, n.s. = not significant. **(K,L)** Representative immunofluorescence of NK1.1+ cells and DAPI in 4^th^ ventricle choroid plexus (K) and cerebellar parenchyma (L) of aged WT mice**. (M)** Same as (K) in Young→Old ReC brain. Red arrowheads mark native NK1.1+ EGFP^-^ cell juxtapose to EGFP+ myeloid cell, marked by white arrowheads. We observed in brains of seven aged reconstituted mice only a single EGFP+NK1.1+ cell (< 3% of all NK1.1+ detected). Complete DGE tables are listed in Table S3.

To examine the role of lymphocytes in driving age-related interferon signaling in microglia *in vivo,* we aged Rag2^-/-^γc^-/-^ mice, which lack B, T and NK cells^57^. We characterized this double-knockout line by performing bulk RNA-seq of punched cerebellar and cortical tissue from young (5 mo) and middle-aged (14 mo) male animals (n=5 per age group; Figure 5G), since mice begin to display substantial age-related expression shifts between 12 to 15 months^7^. As a reference dataset, we meta-analyzed Bulk-seq data from brain tissue of WT mice aged 3, 12 and 15 months (n=5 per aged group), that were processed via the same Bulk-seq method^7^. While DAM-like genes *Itgax* and *Lpl* were evenly upregulated in cerebellar tissue of aged WT and Rag2^-/-^γc^-/-^, only WT mice displayed an increased expression of IFN-I and IFN-II interferon response genes (*Stat1, Oasl2*, *Ifi27l2a*, *Ifit3, Isg15*, *Psmb8;* Figure 5H-J; Table S3). In line with our CellChat analysis, genetic depletion of lymphocytes thus prevented age-related interferon signaling in cerebellar microglia.

Given that our *in silico* predictions differed from previous analyses^12^, we investigated NK cells as potential drivers of IFN-II responsive genes in WT microglia. We confirmed the presence of NK cells in the cerebellum-adjacent fourth ventricle choroid plexus (ChP) of WT mice^58, 59^, and observed rare instances of NK cells in close proximity to microglia within the cerebellum of aged mice (Figure 5K,L). In the ChP of reconstituted animals, we observed predominantly host-derived EGFP^−^ NK cells, despite successful repopulation of the same niche by EGFP+ myeloid cells (Figure 5M). Indeed, NK cells were not replaced in our reconstitution protocol (Figure 1G), which aligns with previous reports of NK cell resilience to chemotherapy and radiation-based bone marrow ablation^37^. As such, these findings support the hypothesis that NK cells may act as persistent drivers of transcriptional changes in both WT microglia and ReCs.

### Absence of T and B cells has limited impact on age-related interferon signaling in microglia

To determine which lymphocyte pool is primarily responsible for the increase in microglia interferon response, we aged a cohort of Rag2^-/-^ female mice^60^, which lack T and B cells but have functional NK cells, and performed scRNA-seq of cerebellar and cortical immune cells at 17 months of age (Figures 6A,B and S13A-D). Surprisingly, cerebellar Rag2^-/-^ and WT microglia exhibited nearly identical expression levels of interferon response genes (*Ifi27l2a*, *Stat1*, *Ifi204*, *Bst2*), and the correlation of log2FoldChanges between cerebellar and cortical microglia persisted both in aged Rag2^-/-^ and aged WT mice (Figures 6C,D and S13E-H). Similarly, cerebellar microglia exhibited a significantly increased CAAS score compared to cortical microglia in aged Rag2^-/-^ mice, mirroring aged WT mice (Figure S13I; Figure 3H).

**Figure 6.**
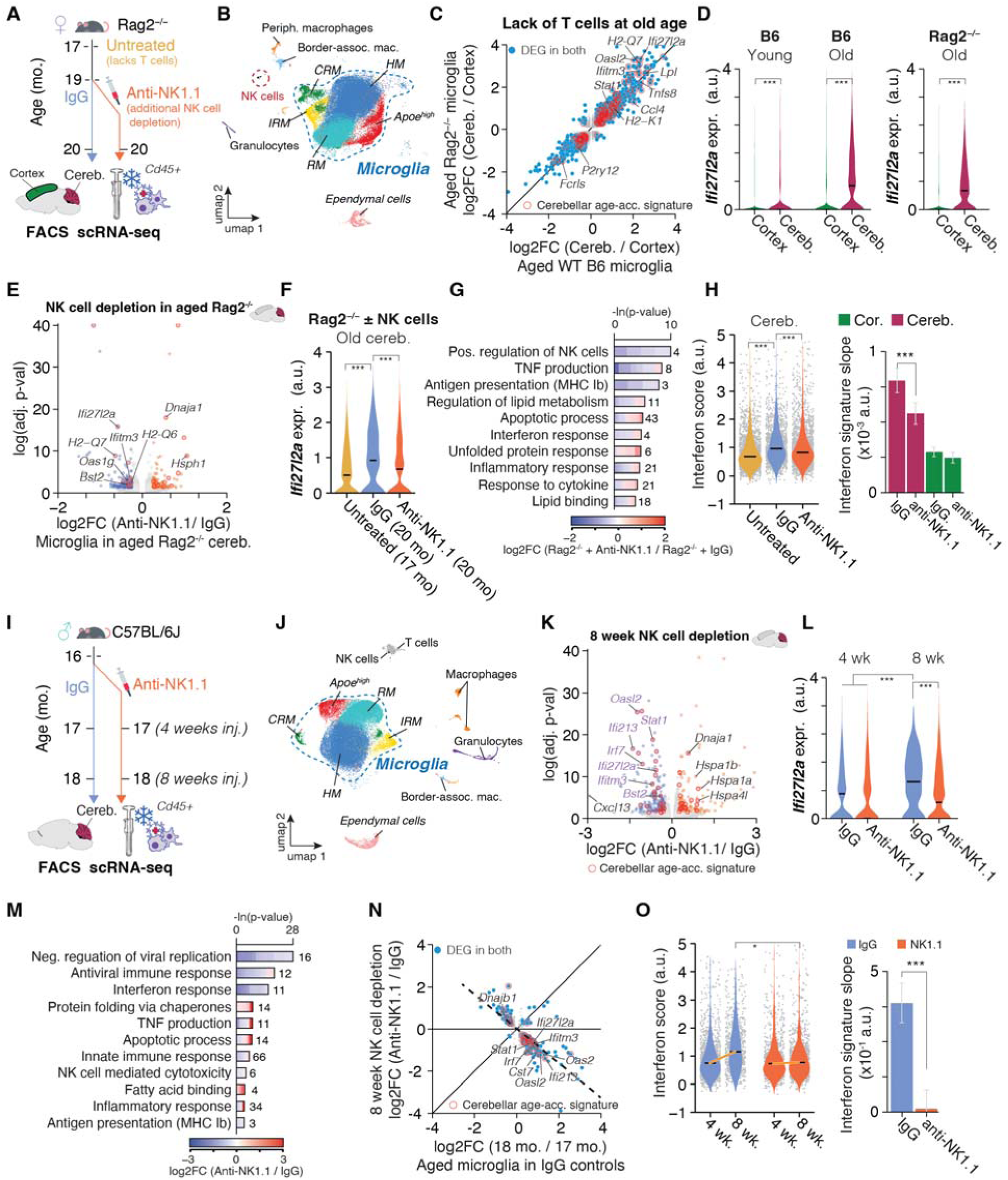
NK cell depletion prevents interferon signaling in aged microglia. **(A)** Experimental design: aged female Rag2^-/-^ mice lacking T cells were either analyzed at 17 months of age (‘untreated’) or received IgG or anti-NK1.1 antibody from 19 to 20 months (IgG, anti-NK1.1). CD45+ immune cells from cerebellum and cortex were analyzed by scRNA-seq. **(B)** UMAP of CD45+ immune cells from Rag2^-/-^ cerebellum and cortex. **(C)** Comparing cerebellum/cortex log2FC in aged Rag2^-/-^ versus aged WT microglia (data from Figure 1B); cerebellar-acceleration signature (CAAS) genes in red. **(D)** *Ifi27l2a* expression in microglia from young (3 mo) and old (18 mo) WT C57BL/6J and aged Rag2^-/-^ mice. Median is indicated. *MAST*, Benjamini-Hochberg correction. ***padj < 0.001. **(E)** DEGs in 20 months-old Rag2^-/-^ cerebellar microglia after anti-NK1.1 treatment: log2FC (anti-NK1.1/IgG) versus –ln(padj); CAAS genes highlighted in red. *MAST*, Benjamini-Hochberg correction. **(F)** *Ifi27l2a* expression in cerebellar microglia from untreated, IgG- and anti-NK1.1–treated Rag2^-/-^ mice. *MAST*, Benjamini-Hochberg correction. ***padj < 0.001. **(G)** Representative GO-term enrichment for DEGs in (E) in aged cerebellum. **(H)** Left, violin plots of interferon response signature score in cerebellar microglia from untreated, IgG- and anti-NK1.1–treated Rag2^-/-^ mice; median indicated. Right, slopes of signature scores by linear regression; mean ± 95% CI; ***p < 0.001 by two-sided *Tukey’s HSD test*. Untreated samples were used as reference for both IgG and anti-NK1.1 treated mice. **(I)** Experimental design: wild-type male C57BL/6J mice received at 16 months of age IgG or anti-NK1.1 antibody for 4 or 8 weeks before cerebellar microglia isolation and scRNA-seq. **(J)** UMAP of CD45+ cells from cerebellum of IgG- and anti-NK1.1–treated C57BL/6J mice colored by clusters as in (B). **(K)** DEGs in cerebellar microglia after 8-week NK cell depletion: log2FC (anti-NK1.1/IgG) versus –ln(padj); CAAS genes in red. *MAST*, Benjamini-Hochberg correction. **(L)** *Ifi27l2a* expression in cerebellar microglia after 4- and 8-week IgG or anti-NK1.1 treatment. Median is indicated. *MAST*, Benjamini-Hochberg correction. ***padj < 0.001. **(M)** Representative GO-term enrichment for DEGs in (K) in aged cerebellum. **(N)** Scatter plot comparing log2FC in cerebellar microglia of 8-week anti-NK1.1–treated C57BL/6J mice versus aged (18/17 mo) IgG controls; CAAS genes highlighted. **(O)** Left, violin plots of interferon signature score in cerebellar microglia after 4 and 8 weeks of IgG versus anti-NK1.1 treatment; median indicated; *p < 0.05, ***p < 0.001 by two-sided Wilcoxon test. Right, slopes of signature scores by linear regression; mean ± 95% CI; ***p < 0.001 by two-sided *Tukey’s HSD test*. Complete DGE and GO tables are listed in Table S3.

To corroborate our observations with independent data, we performed a meta-analysis scRNA-seq data of subcortical white matter- and cortical grey matter-derived microglia from 24-month-old Rag1^-/-^ mice, which also lack T and B cells^23^, and age-matched controls (Figure S14A-E). We note that this dataset did not contain cerebellar-derived cells. White matter-derived microglia exhibited a heightened interferon response, increased expression of DAM-like genes, and a markedly elevated CAAS (Figure S14F-G; Table S3). Compared to WT mice, white matter-derived microglia from Rag1^-/-^ animals showed a mild dampening of the CAAS, but maintained markedly higher interferon response gene expression relative to grey matter microglia (Figure S14H). These findings extend our observations that the absence of T and B cells alone is insufficient to reduce the age-related increase in interferon signaling in microglia, particularly in the cerebellum.

### Depletion of NK cells prevents age-related interferon signaling in microglia

Despite NK cells’ resilience to broad chemotherapy-based ablation, intraperitoneal injections of anti-NK1.1 antibody can effectively and specifically deplete circulating and tissue-resident NK cells in mice, including those in the brain^61, 62^. To determine whether NK cells are necessary for driving the age-related increased interferon signaling response in microglia, we administered biweekly injections of anti-NK1.1 antibody or IgG control^63^ for four weeks to 19 month old Rag2^-/-^ mice, from the same cohort as described above. Compared to IgG controls, anti-NK1.1 treatment significantly decreased NK cell numbers in the cerebellum (Figures S13A and S15A), and caused a significant downregulation of interferon response and antigen presentation genes (*Ifi27l2a*, *Ifitm3*, *Bst2*, *H2–Q7)* in microglia (Figures 6E-H and S15B,C; Table S3). Consistent with Rag2^-/-^γc^-/-^ mice, anti-NK1.1 treatment dampened specifically interferon response genes but caused no significant reduction of DAM-like genes or the CAAS as a whole (Figure S15D-E).

We next tested whether depletion of NK cells alone would be sufficient to affect the induction of age-related interferon signaling in cerebellar microglia. WT C57BL/6J mice were treated with biweekly injections of anti-NK1.1 antibody or IgG control starting at 16 months of age, shortly before most brain regions begin showing accelerated age-related transcriptional changes^7, 64^ (Figures 6I,J and S15F-H). In IgG-treated mice, scRNA-seq of cerebellar immune cells at 17 and 18 months revealed an age-related increase in interferon response, whereas anti-NK1.1 treatment significantly prevented this change (Figure 6K,L; Table S3). At 18 months, NK-depleted mice showed significant downregulation of interferon signaling, innate immune response and antigen presentation pathways in cerebellar microglia, without affecting DAM-like gene expression patterns (Figure 6M-O). These findings identify NK cells as necessary drivers of the age-related interferon response in cerebellar microglia.

## Discussion

The importance of the neuro-immune homeostasis network to support lifelong brain health is becoming increasingly clear, and microglia, the resident immune cells of the CNS, play a central role in its maintenance^4, 47^. Here, we present a scalable and robust myeloid cell replacement strategy that, for the first time, was successfully applied in aged animals, enabling controlled introduction and genetic manipulation of intrinsically young cells into the aged brain (Figure 7A). This platform, compatible with Cas9-mediated editing and pooled perturbation approaches^65^, overcomes limitations of viral transduction in primary microglia and offers a powerful system to dissect cell-extrinsic drivers of microglial biology in the context of aging and possibly disease backgrounds.

**Figure 7.**
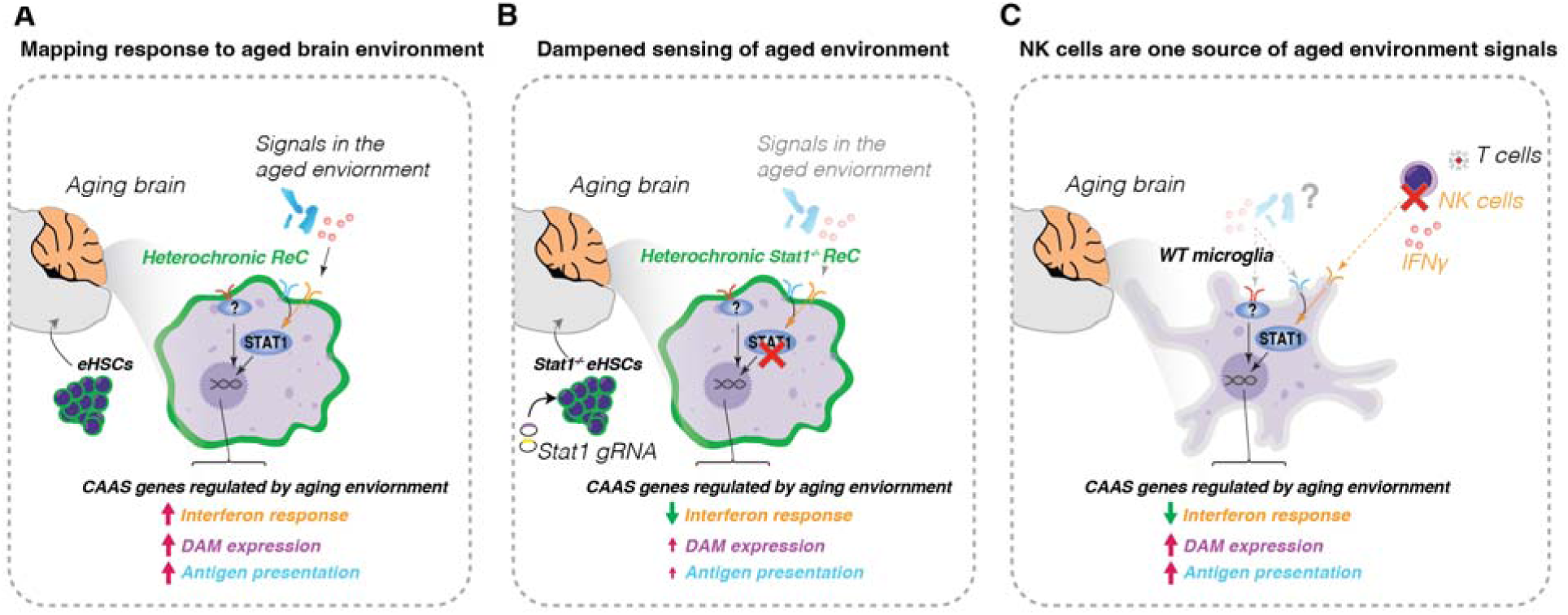
Model for the environmental control of microglia aging. **(A)** Intrinsically young myeloid cells (ReCs) transplanted into an aged brain acutely adopt age-related transcriptional programs, an effect most pronounced in the cerebellum. These programs are characterized by a heightened interferon response, the activation of DAM genes, and increased antigen presentation. **(B)** Genetic deletion of *Stat1* in these young donor cells blocks their ability to fully respond to the aged environment. This cell-autonomous intervention protects ReCs from acquiring age-related expression signatures, identifying STAT1 signaling as an important mediator of environmentally-driven signatures of aging. **(C)** Our findings identify NK cells as a necessary upstream driver of the age-related interferon response in microglia. NK cells likely act via interferon-gamma (IFN-γ) to activate the STAT1 pathway, linking the local cellular environment to the molecular aging of microglia. It is likely that multiple, potentially region-specific, extrinsic signals beyond NK cells act on microglia in the aging brain.

The ability of ReCs to adopt region-specific expression profiles and morphological features indicates that the mechanisms for sensing external cues that shape microglial identity and function are not exclusive to microglial origin, but are preserved in developmentally distinct myeloid cells. Given the critical roles of microglia in brain immune surveillance^4, 47, 66^, such redundancy may serve as a protective mechanism, allowing non-resident myeloid cells to assume essential functions when endogenous microglia are absent or impaired^67^. Though ReCs and resident microglia have distinct ontogeny, ReCs nonetheless rapidly acquire region-specific transcriptional signatures that are seen in microglia that have been present since embryonic development^68^. In line with previous reports of microglia adopting region- and age-specific expression patterns post-PLX treatment^69, 70^, native microglia sampled from the brains of reconstituted mice also displayed these region-dependent profiles. This further supports the notion that age-related and region-dependent changes observed in ReCs are likely not driven by intrinsic features specific to eHSCs or peripheral programming of eHSCs. Peripheral cell trafficking into the CNS can occur through the blood-brain barrier^71, 72^ via the endothelium, the choroid plexus (ChP) with its leaky blood-stroma interface^73, 74^, and the velum interpositum, a route implicated in myeloid cell entry during development, after irradiation, and in demyelinating disease^75^. Whether distinct entry points into the brain contribute to the phenotypic diversity of circulation-derived myeloid cells warrants further study.

Utilizing the flexible gene editing capabilities of our platform, we successfully knocked out *Stat1* in ReCs, which revealed a central role for interferon signaling in driving age-related transcriptional changes in the cerebellum in ReCs and likely microglia (Figure 7B). Interestingly, the deletion of *Stat1* exerted only a limited effect on the ability of ReCs to adopt cerebellar-specific molecular and morphological features at young baseline, suggesting other mechanisms are mediating the heightened immune-vigilant expression phenotype of cerebellar microglia at young age^19, 31^. Recent studies have highlighted CSF-1 and the cytokine IL-34 as important modulators of microglial tiling density, morphology and function, such as phagocytic capacity, in the cerebellum and forebrain^32, 34^. Notably, blocking IL-34 in the cortex produces less ramified microglia, resembling those typically found in the cerebellum^32, 34^. Cell-intrinsic mechanisms likely also contribute to age-related changes and may converge with region-specific influences^4^. Repeated microglial depletion with PLX causes aging-like molecular and functional changes in repopulated microglia, suggesting that extensive turnover may drive microglial exhaustion and impair homeostasis^76, 77^. Additionally, release of mitochondrial DNA into the cytoplasm of aged microglia can activate the cGAS-STING pathway, in turn triggering neurodegeneration in the hippocampus and leading to memory impairment^20^. The extent to which cell-intrinsic mechanisms contribute to the regional heterogeneity of age-related microglial changes remains unclear.

In addition to recent data that suggest a role for T cells in driving the age-related interferon response in microglia^12, 46^, we identified NK cells as previously unrecognized and necessary drivers in the aged cerebellar microglia (Figure 7C). *In vitro* IFN-γ treatment of microglia induced a transcriptional response resembling that of aged cerebellar microglia, supporting IFN-γ as an important mediator of this phenotype. In young healthy mice, NK cells comprise only ∼1% of total CNS immune cells and are largely restricted to boundary regions such as the meninges and ChP^78, 79^. We consistently observed NK cells in the ChP of the 4^th^ ventricle but found only rare instances within the cerebellar parenchyma. Transient NK cell localization in the cerebrum has been reported during remyelination^62^, raising the possibility that short-lived NK cell infiltration could also occur during aging but is difficult to capture. NK cell accumulation has been shown in the aged hippocampus of both humans and mice, leading to impaired neurogenesis and cognition^63^, whereas depletion in Alzheimer’s disease models lead to improved cognitive function and reduced cortical neuroinflammation^80^. Interestingly, NK cell-mediated IFN-γ signaling in response to cytomegalovirus in newborns can disrupt cerebellar development^81^, but beyond this, little is known about NK cell function in the cerebellum, and we are not aware of previous work describing their role in cerebellar aging. We show that depleting NK cells eliminated the age-related interferon increase in cerebellar microglia, establishing NK cells as an essential upstream driver of this pathway. Future studies are needed to determine if this effect is mediated by direct NK cell secretion of IFN-γ or indirect activation of local IFN-γ-producing cells^82^. How other IFN-γ-producing cells, like tancycytes, might drive microglial aging phenotypes in other brain regions like the hypothalamus, also warrants further investigation.

It is likely that multiple, potentially region-specific, extrinsic signals beyond interferons act on microglia in the aging brain. The region- and age-dependent expression patterns identified in our results may provide insights into identifying these signals and the mechanisms by which microglia sense them. For instance, cerebellar ReCs in the arbor vitae exhibited the strongest DAM-associated changes and those in the molecular layer the weakest, consistent with our data demonstrating that oligodendrocyte-adjacent ReCs had a significantly elevated CAAS score and other reports of white-matter associated activation^83, 84^. While *Stat1* knockout broadly suppressed age-related increases in interferon response, DAM activation, and antigen presentation in ReCs, only the interferon response was reduced in microglia from Rag2^-/-^γc^-/-^ and NK cell-depleted animals. Thus, alongside NK cells, white matter-derived factors like myelin debris may be converging in parallel on STAT1 to promote immune vigilance and shape region-specific microglial aging^85^.

Our findings demonstrate that the aging brain enters a feed-forward loop that promotes age-related changes even when old cells are replaced with intrinsically young cells, challenging the concept of ‘rejuvenation’ via cell replacement^86^. Disrupting this loop through targeted ablation of key cell types or pathway-specific interventions, such as JAK-STAT inhibition, can reset these interactions, and may represent an interesting target for intervention in humans. The *in vivo* reconstitution approach described here provides a scalable approach to uncover additional modulators of microglial aging and potential therapeutic targets for age-related inflammation and neurodegeneration.

### Limitations of the study

This study primarily addresses age-related effects on microglia and ReCs, leaving the effects of NK cell depletion on other cell types, such as oligodendrocytes, to be explored. We focused on immune cells within the brain parenchyma and ChP, and cannot infer how myeloid cells in other niches, such as the leptomeninges^79^, respond to their local environments. Our approach to defining CAAS genes was intentionally conservative, requiring differential expression in both WT microglia and ReCs, and across two regions. This strategy likely excluded true positives. As such, *Gpnmb* and *Spp1* were excluded from the CAAS. However, we cannot rule out the potential residual effects, which has been reported to cause mild white matter damage^28^, as a contributor to the elevated expression of DAM-like genes in ReCs. Analysis of Spatial-seq data involved computational pooling by age and cell type without replicate sensitivity, raising the possibility of pseudo-replication. However, we provide replicate-resolved plots for key results. The low abundance of NK cells in the ChP prevented direct assessment of age-related transcriptional or numerical changes in this compartment. Despite these limitations, our research lays the groundwork for future investigations into the age-related and region-specific factors that shape microglia heterogeneity. Finally, this study utilized both male and female mice across different experimental arms, including a female-to-male transplantation model in our primary experiments. While key findings were replicated in sex-matched paradigms, we cannot fully exclude potential contributions of sex-specific variables to our observations.

## Acknowledgments

We thank all members of Calico’s Tissue Homeostasis division and members of the Stevens and Wernig labs for feedback and support, and appreciate the Lab Operations Team for assistance in laboratory management. We greatly appreciate the support by Calico’s Laboratory Animal Resources Team throughout this study. We particularly thank Calvin Jan, Carmela Sidrauski, Michael Closser, Sandip Chatterjee and Phillip Seitzer for internal review of the manuscript and code base. We further thank Michael Dolan and Raphael Rakosi-Schmidt for advice on NK cell staining techniques. The research leading to these results received funding from the Simons Foundation (B.S.), the Cure Alzheimer’s Fund (B.S.), the Howard Hughes Medical Institute (B.S.) and Calico Life Sciences (B.S., O.H.).

## Author contributions

C.G., G.P. and O.H. conceptualized the study and prepared the manuscript. G.P. and H.S. conducted isolation and expansion of HSCs and culture of neonatal microglia, with help from M.A. and K.L.; B.J.W. established and conducted gene editing of eHSCs; C.G., G.P., H.S., W.C. and B.M-M. conducted myeloid cell replacement experiments; T.D.L. analyzed cultured eHSCs and conducted splenocyte analysis; C.G., H.S. and O.H. conducted tissue collections; H.S. and O.H. conducted all bulk- and scRNA-seq experiments with help from N.F. and D.J. Spatial-seq experiments were conducted by W.K., H.T. and C.G.; NK cell depletion experiments in Rag2^-/-^ and WT mice were planned and carried out by C.G., W.C., B.M-M., A.W. and O.H.; Rag2^-/-^γc^-/-^-derived brains were provided by C.D. M.H., and B.S.; M.M.M., D.H., A.G. and A.C. aided in experimental strategy and method design. Experiments involving transplantations of bone marrow–derived myeloid cells were carried out by M.M.M. and Y.S., with guidance and conceptualization by M.W.; Y.Y. prepared bulk-seq libraries of bone marrow-derived myeloid cells; C.G. conducted confocal imaging and tracing analysis; O.H. conducted all bioinformatic and statistical analysis; C.G., G.P. and O.H. edited the manuscript with input from all authors. All authors read and approved the final manuscript.

## Supplementary Figure titles and legends

**Figure S1.**
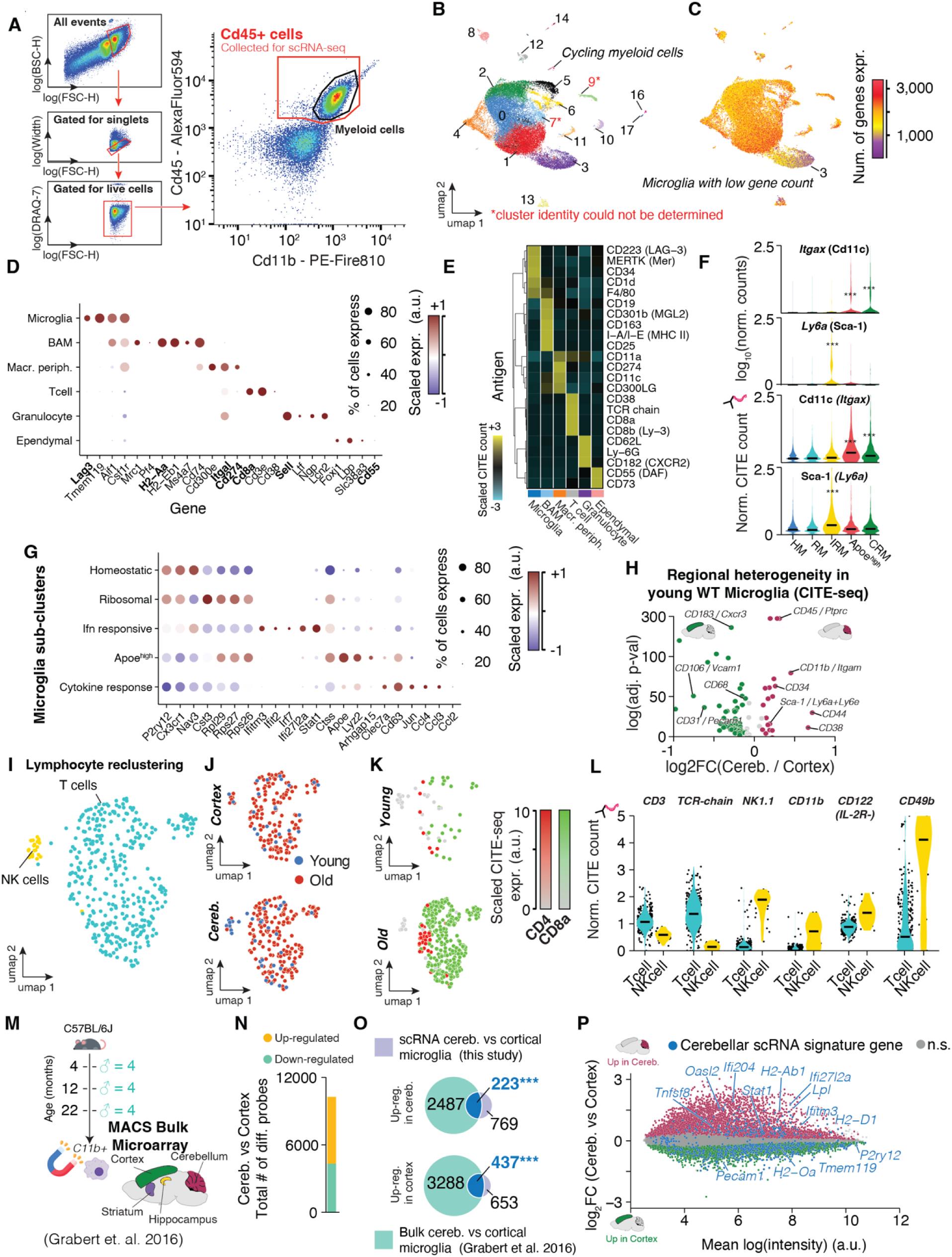
Profiling and annotation of brain immune populations by CITE- and scRNA-seq. **(A)** Representative FACS gating strategy for isolating live CD45+ cells (stained for both CD45 and CD11b) from cortex and cerebellum, including gating for singlets and viability, followed by collection for scRNA-seq. **(B)** UMAP of all CD45+ cells annotated by cluster identity; cluster 3 contained cycling myeloid cells, and cluster 13 contained microglia with low gene counts. **(C)** Number of detected genes per cell overlaid on UMAP. **(D)** Dot plot showing scaled RNA expression (color) and percentage of expressing cells (dot size) for selected cell type–specific marker genes across annotated populations. **(E)** Heatmap of selected, scaled CITE-seq counts for indicated surface proteins across annotated immune populations. **(F)** Violin plots of normalized RNA counts for *Itgax* and *Ly6a* in indicated microglial clusters at young age, with corresponding scaled CITE-seq counts for CD11c and Sca-1, respectively. Center line = median. Statistical significance assessed by two-sided *Wilcoxon rank-sum*. **(G)** Dot plot showing scaled RNA expression of subtype-defining genes across microglial subclusters: homeostatic microglia (HM), interferon response microglia (IRM), ribosomal processing microglia (RM), cytokine response microglia (CRM), Apoe^high^ microglia. Complete results listed in Table S1. **(H)** Volcano plot of differential surface protein expression in WT microglia (cerebellum vs cortex) as quantified via CITE-seq. Protein expression shifts were assessed via *MAST* and considered significant if Benjamini-Hochberg-corrected padj < 0.05 and |log2FC| > 0.1. Complete results listed in Table S1. **(I)** UMAP of reclustered T and NK cell subsets from all samples. Cells from cluster 8 from (B) were re-clustered and re-integrated. **(J–K)** UMAPs of cortex **(J)** and cerebellum **(K)** lymphocytes colored by age group (young vs old), overlaid with scaled CITE-seq counts for CD8a and CD4 to indicate CD8^+^ and CD4^+^ T cells. **(L)** Violin plots of normalized CITE-seq counts for T cell and NK cell surface markers (CD3, TCR-chain, NK1.1, CD11b, CD122, CD49b) in indicated lymphocyte subtypes. Center line = median. **(M)** Schematic of meta-analysed MACS bulk microarray dataset from Grabert et al.^19^, showing regions profiled in C57BL/6J mice (n = 4 per age group). **(N)** Counts of upregulated (yellow) and downregulated (turquoise) probes in cerebellum versus cortex from bulk microarray datasets; Statistical significance of differential expression was assessed using limma’s *moderated t test* with FDR correction. **(O)** Venn diagrams show overlap of DEGs from bulk microarray and scRNA-seq method; ***padj < 0.001 by *Fisher’s exact test*. **(P)** MA plot comparing mean RNA expression versus log2FC(cerebellum/cortex) in bulk microarray data. Non-significant genes are in grey. *padj < 0.05. Statistical significance was assessed using limma’s *moderated t test* with FDR correction. Matching region-specific DEGs detected by scRNA-seq are indicated.

**Figure S2.**
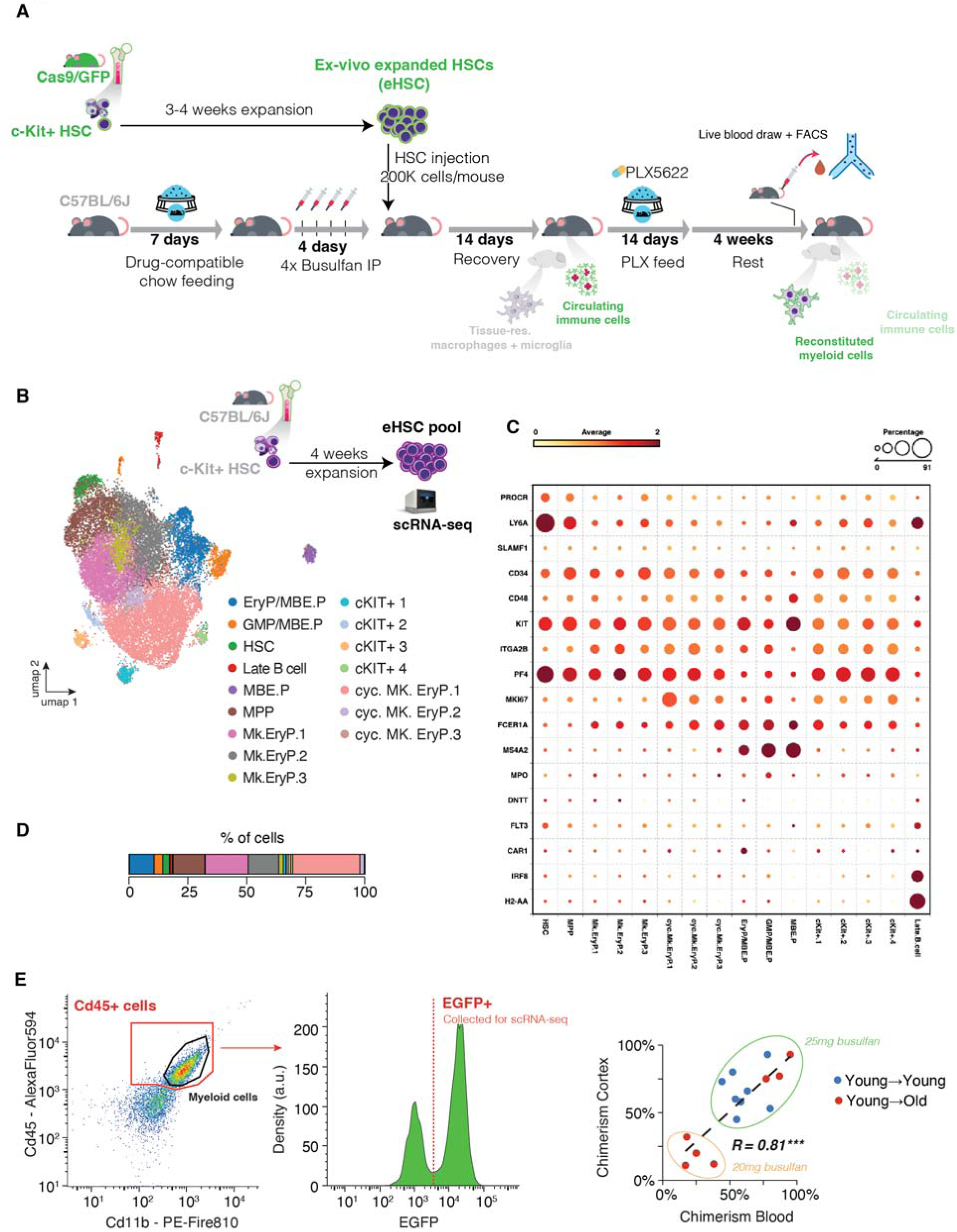
Experimental workflow and transcriptional characterization of eHSC populations. **(A)** Schematic of heterochronic myeloid replacement workflow. Cas9/GFP+ c-Kit+ HSCs of female animals were expanded in vitro for 3-4 weeks, followed by retro-orbital injection (200,000 cells/mouse) into busulfan-conditioned, male WT C57BL/6J recipients after drug-compatible chow feeding and 4 daily intraperitoneal busulfan injections (25mg/kg body weight for young recipients; aged mice received injections of 25 or 20 mg/kg body weight). Mice recovered for 14 days before receiving PLX5622 chow for 14 days to deplete tissue-resident macrophages and microglia. After 4 weeks of rest, blood was collected for flow cytometric analysis of circulating immune cells via tailbleed. At the end of the experiment, mice were perfused and brains were processed to assess reconstituted myeloid populations. **(B)** UMAP embedding of scRNA-seq data from in vitro–expanded c-Kit+ eHSC pools (n=3 pooled wells), annotated by cluster identity: erythroid progenitors/megakaryocyte–erythroid progenitors (EryP/MBE.P), granulocyte–monocyte progenitors (GMP/MBE.P), hematopoietic stem cells (HSC), late B cells, multipotent progenitors (MPP), multiple c-Kit+ cell clusters (1–4), cycling erythroid and megakaryocyte progenitors (cyc. MK.EryP), and megakaryocyte progenitor subclusters (Mk.EryP.1–3). **(C)** Dot plot of scaled expression (color) and percent expression (dot size) for canonical lineage and progenitor marker genes across clusters. **(D)** Relative abundance of each cluster as percentage of all cells in the eHSC pool. **(E)** Representative FACS gating for CD45+ cells from cortex and cerebellum, including gating for EGFP+ events collected for scRNA-seq (middle). Right, correlation of cortical versus blood chimerism in young→young (blue) and young→old (red) recipients receiving busulfan at dosages of 20 mg or 25 mg/kg bodyweight; p < 0.001 by Pearson correlation.

**Figure S3.**
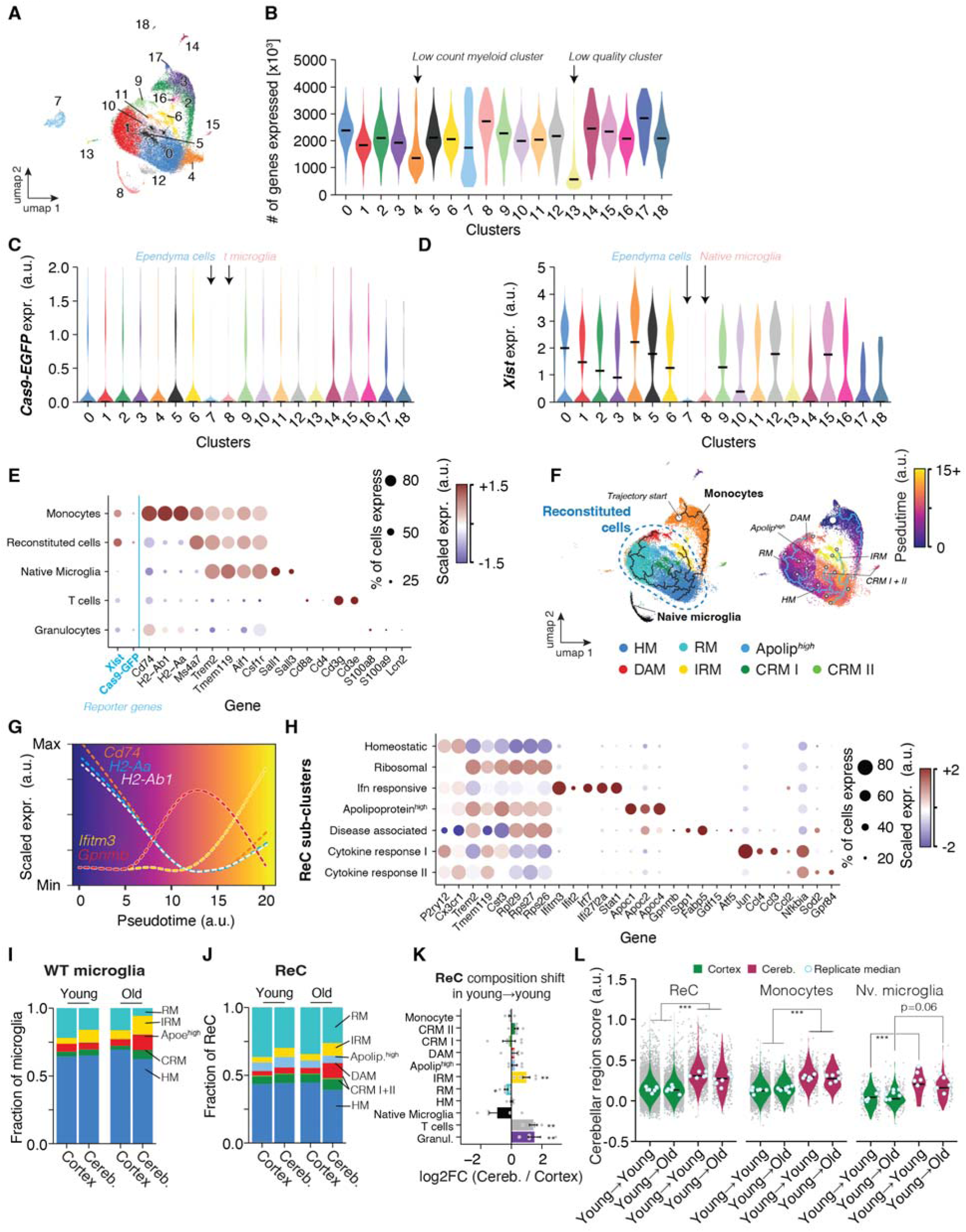
Profiling and annotation of reconstituted myeloid cells by scRNA-seq. **(A)** UMAP embedding of all cells collected via Cd45+/EGFP+ gating, annotated by cluster identity. **(B)** Violin plots showing the number of genes detected per cell across clusters. Center line = median. Clusters with low gene (cluster 4) or low quality (cluster 16) are indicated - these were censored going forward. **(C–D)** Violin plots of normalized expression for Cas9–EGFP **(C)** and female marker gene *Xist* **(D)** across clusters to indicate clusters with low donor chimerism. **(E)** Dot plot showing scaled RNA expression (color) and percent expressing cells (dot size) for selected lineage markers across annotated immune cell types. **(F)** UMAP of reconstituted cells and native microglia annotated by cell state: homeostatic myeloid cells (HM), interferon response myeloid cells (IRM), ribosomal processing myeloid cells (RM), cytokine response myeloid cells (CRM), Apoe^high^ cells, Disease-associated myeloid cells (DAM); pseudotime coloring indicates inferred trajectory from monocytes to ReCs. **(G)** Pseudotime plot showing scaled expression of representative genes, including *Cd74*, *H2-Aa*, *H2-Ab1*, *Ifitm3*, and *Gpnmb*. **(H)** Dot plot showing scaled RNA expression (color) and percent expression (dot size) for representative marker genes across ReC subclusters. **(I)** WT microglial and **(J)** ReC cluster composition. Complete results listed in Table S1. **(K)** log2FC in ReC cluster proportions at young age across regions. Mean ± SEM. Two-sided *Wilcoxon rank-sum tests*. *padj < 0.05; **padj < 0.01). **(L)** Cerebellar-region score in ReC, monocytes and native microglia. Center line = median. Significance were tested across regions and age groups within a given cell type via two-tailed *t test* on per-replicate median of score. Pvalues we adjusted for multiple testing. The highest (least significant) padj value per cell type is indicated. *p < 0.05, **p < 0.01,***p < 0.01; n = 5-7/group.

**Figure S4.**
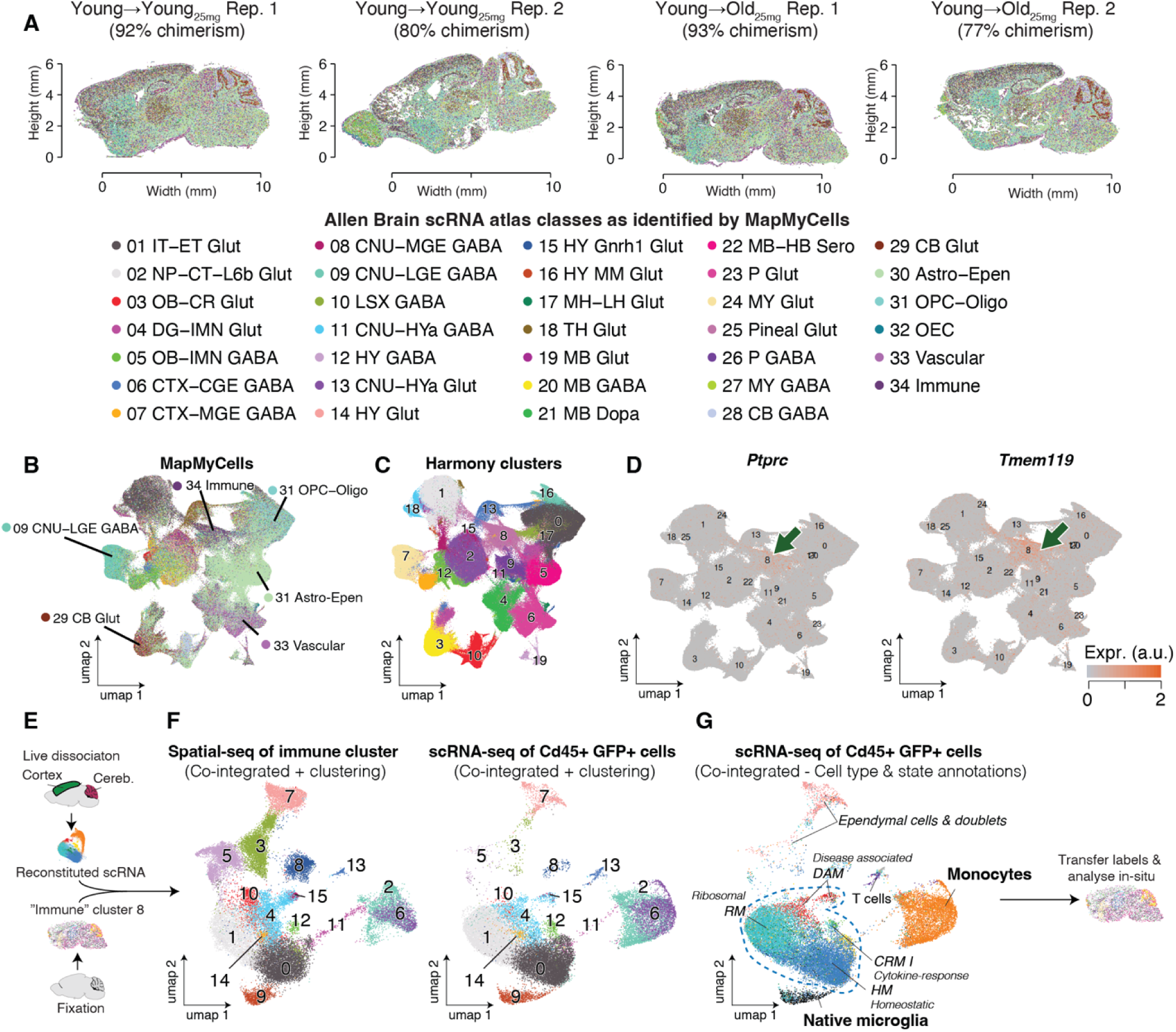
Co-integration of spatial-seq (CosMx) data with scRNA-seq to annotate ReC clusters in situ. **(A)** CosMx spatial maps from young→young and young→old chimeras (n=2 replicates per group; chimerism as quantified by FACS of cortical CD45+ of the same animals is indicated), overlaid with cell class labels from Allen Brain Atlas reference mapping (10x Whole mouse brain taxonomy CCN20240722) using MapMyCells^88^. 1 slice per age group was processed on the same CosMx slide to average out batch effects. **(B)** UMAP of MapMyCells-annotated spatial-seq data showing distribution of major cell types, with immune cluster highlighted. **(C)** Harmony-integrated clustering of spatial-seq data colored by cluster identity. **(D)** Expression maps for *Ptprc/*CD45 and *Tmem119* in Harmony clusters; green arrows indicate immune clusters enriched for ReCs. **(E)** Schematic of co-integration workflow: immune cluster from spatial-seq was co-integrated with scRNA-seq data from Figure 1K, followed by transfer of labels for in situ annotation. **(F)** UMAP of spatial-seq immune cluster after co-integration and clustering with scRNA-seq of GFP+CD45+ cells. **(G)** UMAP of co-integrated GFP+CD45+ cells annotated by cell type/state. IRMs were not identified due to the absence of most interferon response genes in the CosMx panel. Falsely identified cells in MapMyCells cluster 8, such as ependymal cells and doublets were removed. Labels were transferred back to spatial-seq data for downstream in situ analysis.

**Figure S5.**
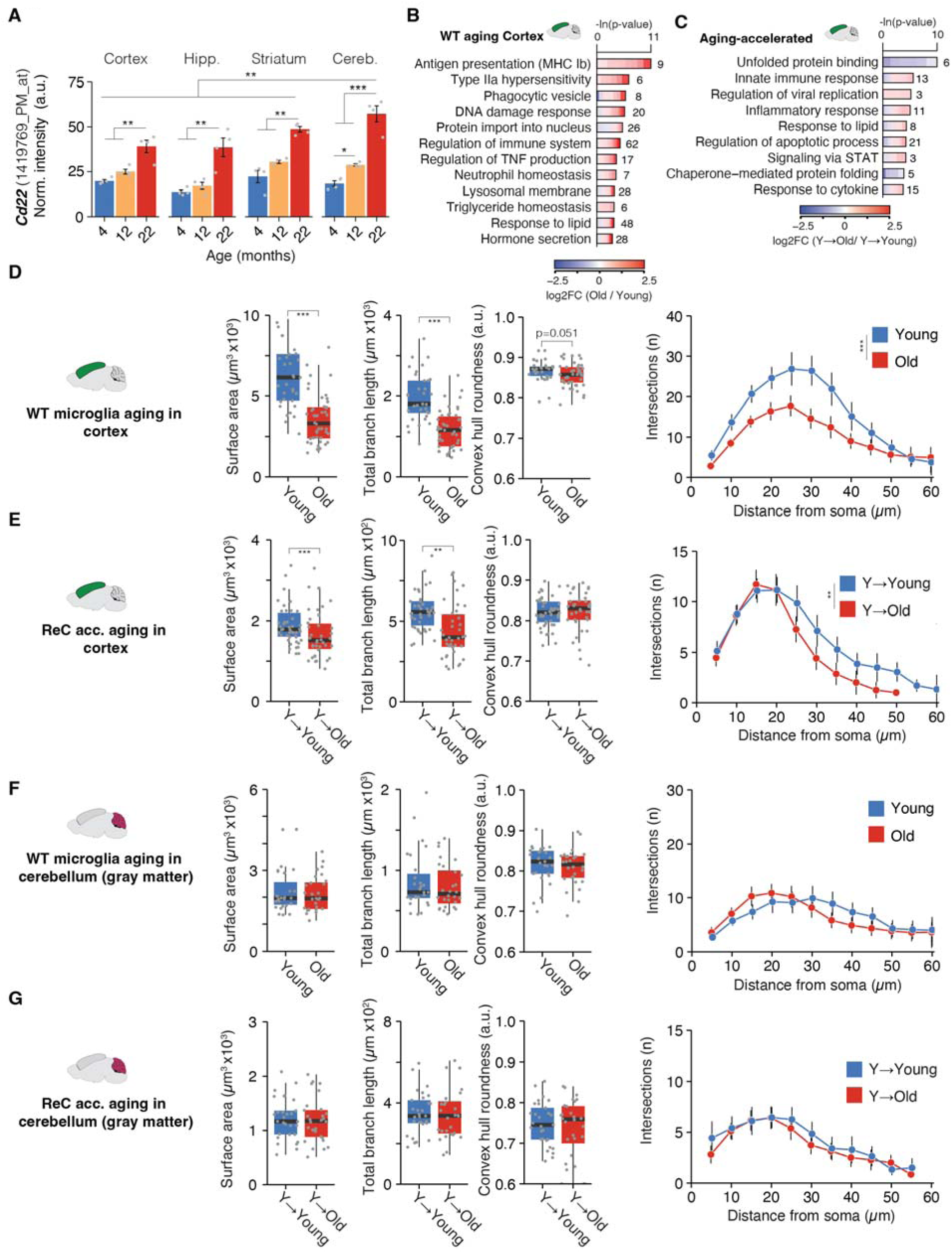
Regional and morphological correlates of age-associated ReC changes in cortex and cerebellum. (A) Quantification of *Cd22* (1419769_PM_at) from microglia bulk RNA microarray data^19^ across cortex, hippocampus, striatum, and cerebellum at 4, 12, and 22 months of age (n = 4/group). Mean ± SD. *p < 0.05, **p < 0.01, ***p < 0.001 by limma’s *moderated t test* with FDR correction. **(B,C)** Representative GO-term enrichment for age-related DEGs in cortical microglia (B), and aging-accelerated DEGs in cortical ReC (C) from young→old chimeras. Complete DGE and GO tables are listed in Table S2. **(D-G)** Age-related shifts in surface area, total branch length, convex hull roundness and Sholl profile (intersections versus distance from soma) in cortical microglia (D), cortical ReC (E), cerebellar microglia (F) and cerebellar ReC (G). (n = 3 mice per group, 12 to 20 cells per animal and region were quantified). Sholl profile shifts were tested via Two-sided *Wilcoxon rank-sum tests* on area under the curve AUC; other metrics were tested via two-tailed *t test*.; **p < 0.01, ***p < 0.001. Sholl curves show mean ± SEM intersections versus distance from soma.

**Figure S6.**
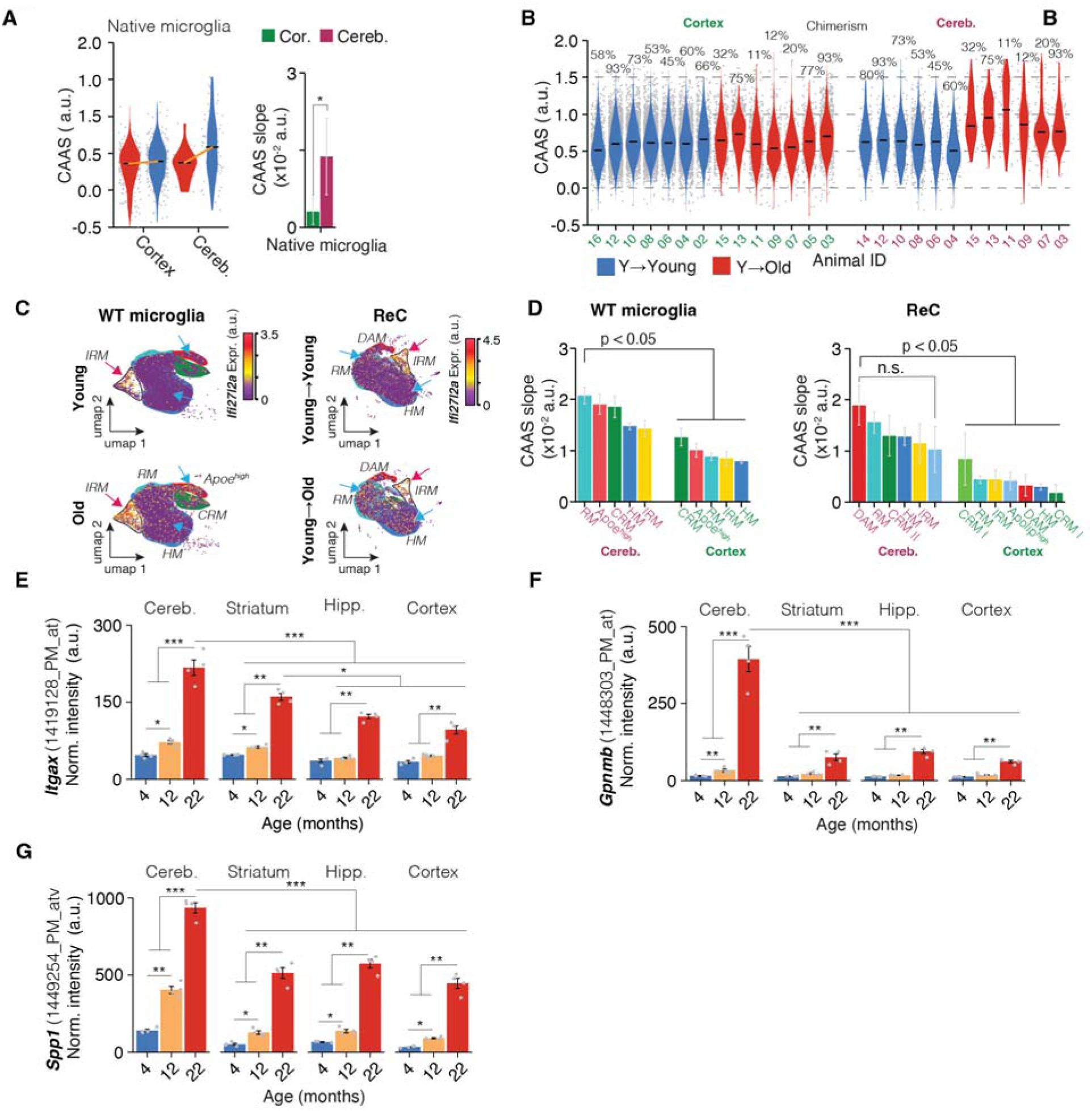
Regional variation in cerebellar-acceleration aging score (CAAS). **(A)** Left, distributions of cerebellar-acceleration aging score (CAAS) in native microglia of reconstituted mice from cortex and cerebellum. Medians are indicated. Right, CAAS slope of linear regressions for each group. Mean ± 95% confidence intervals. Two-sided *Tukey’s HSD test*. The highest (least significant) p value is indicated. *p < 0.05. **(B)** Per animal-resolved CAAS scores in ReCs, with corresponding cortical chimerism percentages indicated above each violin plot. Center line = median. **(C)** UMAPs of WT microglia (left) and ReCs (right) from young and aged mice, colored by normalized expression of *Ifi27l2a*; major states are annotated (HM, RM, IRM, Apoe^high^, DAM, CRM). Blue arrows indicate the age-related, elevated expression of interferon response gene *Ifi27l2a* outside the IRM **(D)** State-resolved slopes of CAAS versus chimerism in cerebellum and cortex for WT microglia (left) and ReC (right). Mean ± 95% confidence intervals. The highest (least significant) p value is indicated. Two-sided *Tukey’s HSD test*. ***p < 0.001, ***p < 0.001, n.s. = not significant. **(E–G)** Bulk RNA microarray quantification^19^ of *Itgax* (E), *Gpnmb* (F), and *Spp1* (G) expression across cerebellum, striatum, hippocampus, and cortex at 4, 12, and 22 months of age (n = 4/group). Mean ± SD. *p < 0.05, **p < 0.01, ***p < 0.001 by limma’s *moderated t test* with FDR correction.

**Figure S7.**
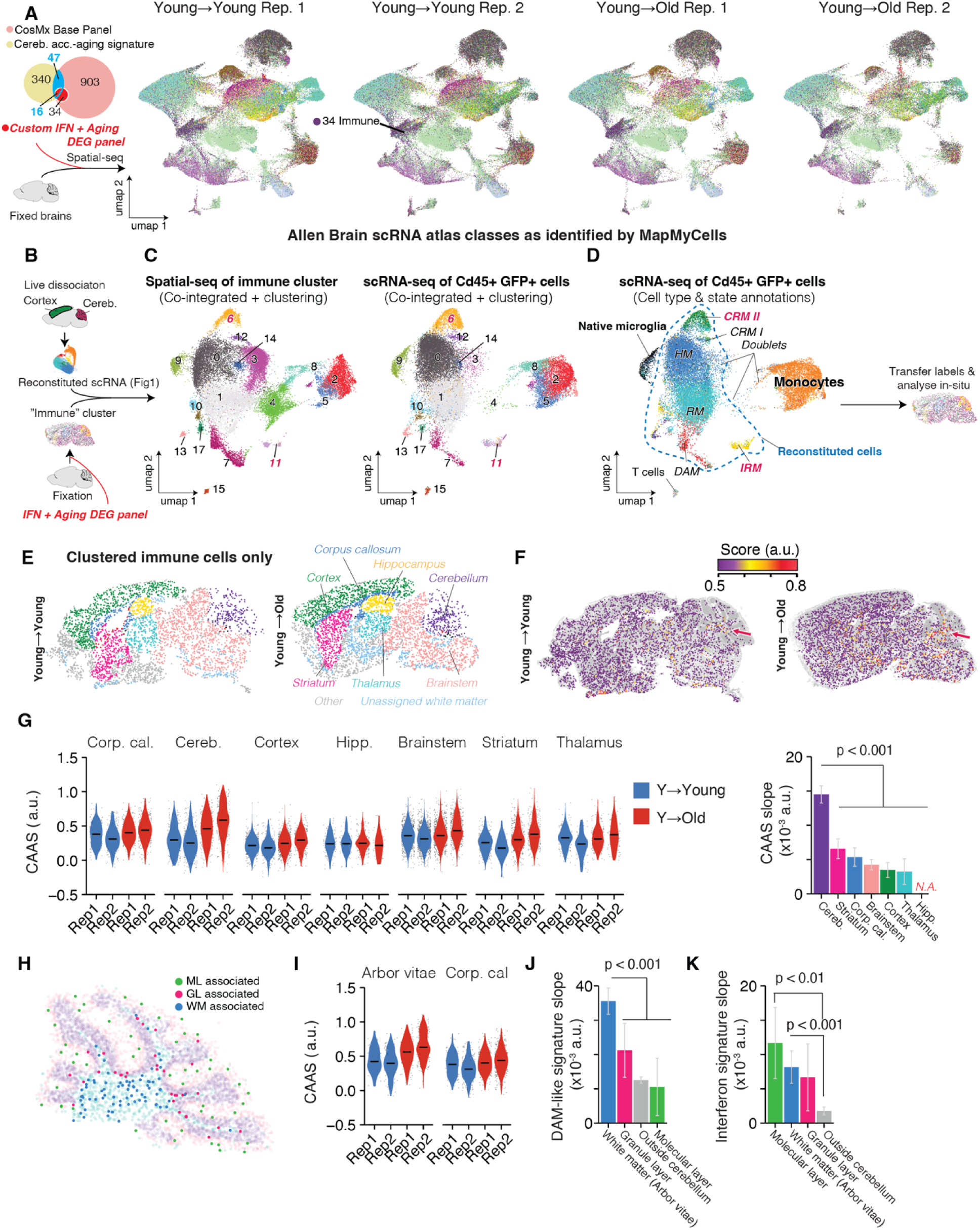
Spatial-seq mapping and in situ analysis of immune cluster states using custom IFN + aging DEG panel. **(A)** Representative CosMx spatial maps generated with custom interferon and CAAS panel from young→young and young→old chimeras (two replicates each), with cells annotated by Allen Brain Atlas reference mapping using MapMyCells. The same brains as in S2 (but differing slices) were analyzed. Immune cluster was utilized for integration with scRNA-seq. Pie chart (left) shows gene panel composition: CAAS, custom panel genes, and overlaps. 1 slice per age group was processed on the same CosMx slide to average out batch effects. **(B)** Schematic of integration workflow: immune cluster from spatial-seq co-integrated with scRNA-seq of reconstituted GFP+CD45+ cells (see Figure S2), clustered, and annotated. **(C)** UMAP of immune cluster cells from spatial-seq after co-integration and clustering with GFP+CD45+ scRNA-seq data. **(D)** UMAP of co-integrated GFP+CD45+ cells annotated by cell type/state. Note that IRMs are now identified as a distinctive cluster **(E)** Regional annotation of ReC in young→young and young→old brains. **(F)** Spatial CAAS score maps for ReCs from young→young and young→old. Cerebellar region is shown in Figure 3. **(G)** CAAS distributions across indicated regions for each replicate, with slopes shown at right. Mean ± 95% confidence intervals. The highest (least significant) p value is indicated. Two-sided *Tukey’s HSD test*. N.A., not applicable where CAAS exhibited no association with age. **(H)** Spatial distribution of ReCs in cerebellum and their classification into white matter (WM), granular layer (GL), and molecular layer (ML). **(I)** Replicate-resolved CAAS in ReCs from arbor vitae versus corpus callosum. **(J)** DAM-like signature slopes for WM-, GL-, and ML-associated immune cells. **(K)** Interferon signature slopes for the same cell groups as in (J). Mean ± 95% confidence intervals. The highest (least significant) p value is indicated. Two-sided *Tukey’s HSD test*.

**Figure S8.**
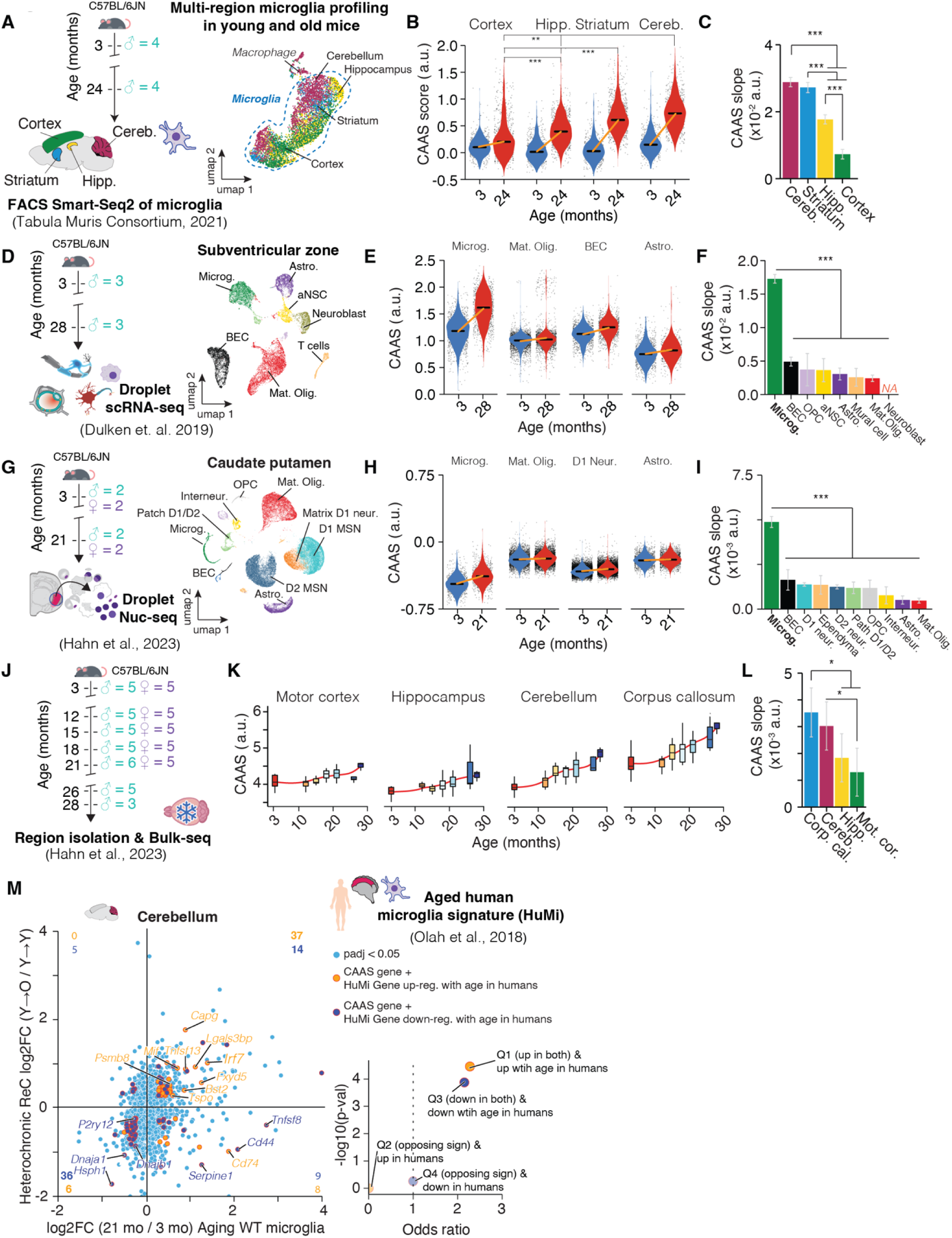
Meta-analysis of CAAS across brain regions, cell types and human data. (A) Schematic of FACS Smart-Seq2 profiling of microglia from cortex, hippocampus, striatum, and cerebellum of C57BL/6J/N mice^49^ (n = 6,373 cells; n = 4 males/age group). **(B)** CAAS distributions across regions and ages (3 mo vs 24 mo). Center line = median; ***p < 0.001 by *two-sided Wilcoxon test* on per-replicate median scores. **(C)** CAAS slope (mean ± 95% CI) across regions from (B); Mean ± 95% confidence intervals. The highest (least significant) p value is indicated. Two-sided *Tukey’s HSD test*. ***padj < 0.001, **padj < 0.01, *padj < 0.05. **(D)** Schematic of droplet-based scRNA-seq from subventricular zone (SVZ) of C57BL/6J mice^46^; (n = 15,684 cells; n = 3 male mice/age group). **(E)** CAAS in SVZ cells. **(F)** CAAS slopes from (E); statistical analysis as in (C). **(G)** Schematic of droplet/nuclear RNA-seq profiling of caudate putamen from C57BL/6J mice^7^ (n = 45,277 cells; n = 2 males, n = 2 females/age group). **(H)** CAAS distributions for caudate putamen cells. **(I)** CAAS slopes from (H); statistical analysis as in (C). **(J)** Schematic of bulk RNA-seq dataset^7^ from selected regions: motor cortex, hippocampus, cerebellum, and corpus callosum of C57BL/6J mice (n = 5/group). **(K)** CAAS versus age for each region. **(L)** CAAS slopes from (K) for corpus cerebellum, hippocampus, motor cortex, and corpus callosum; *padj < 0.05. **(M)** Left, cerebellar scatter comparing per-gene log2FC(Old/Young) in WT microglia (x-axis) versus heterochronic ReCs (y-axis), restricted to genes with padj < 0.05 in both analyses. Genes significant in both and |log2FC| > 0.25 are highlighted. Human microglia (HuMi) genes that change with age^50^ are overlaid and colored by direction in humans (up or down with age). Right, quadrant summary showing enrichment (odds ratio, one-sided *Fisher’s exact test*) of HuMi genes up-regulated or down-regulated with age in humans across each concordance quadrant: Q1 (up in WT & up in ReC), Q2 (up in WT & down in ReC), Q3 (down in WT & down in ReC), Q4 (down in WT & up in ReC). Points are positioned by odds ratio (x-axis) and –log10(p-value) (y-axis). Full analysis results listed in Table S2.

**Figure S9.**
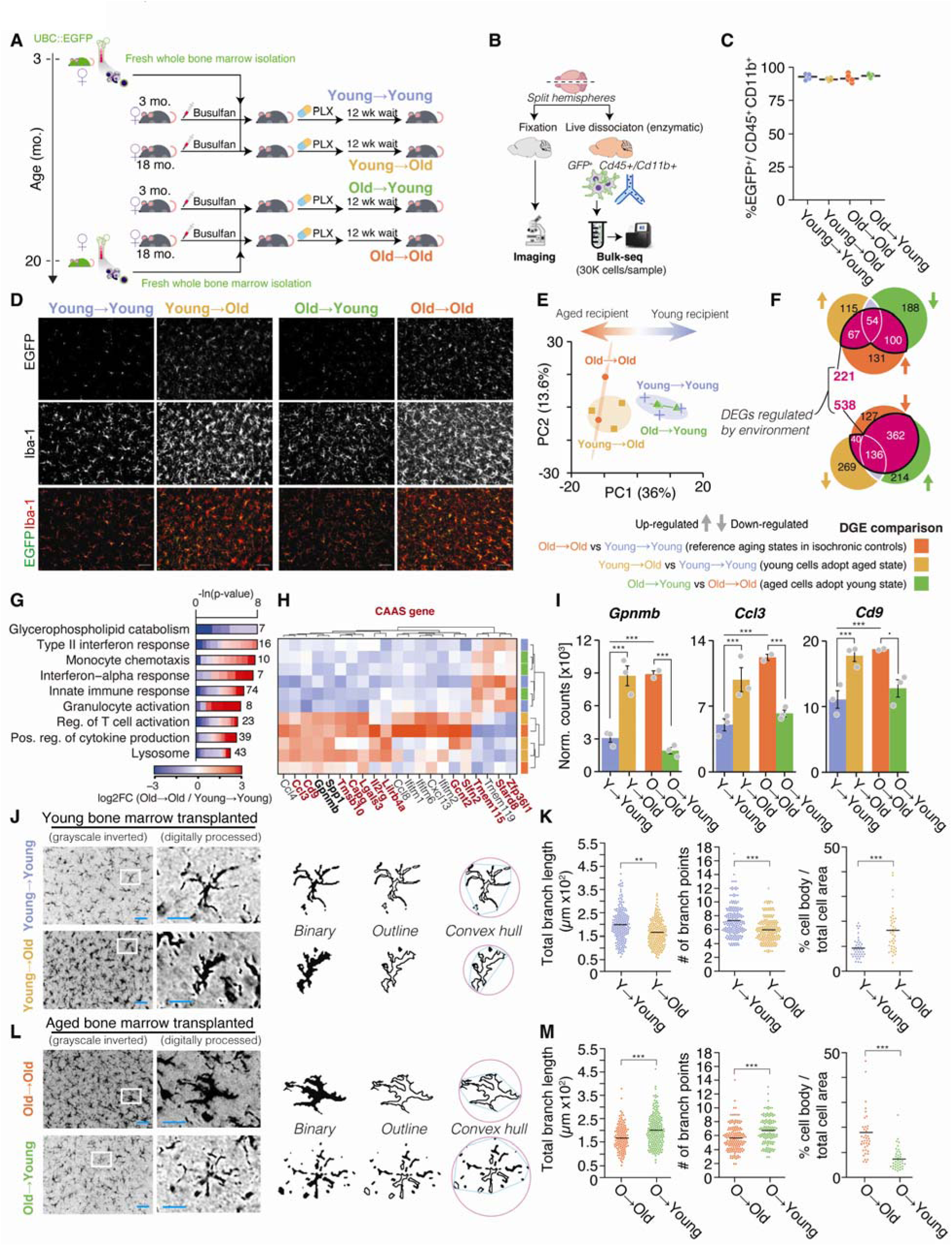
Transcriptomic and morphological effects of the young and aged brain environment on bone marrow–derived myeloid cells. **(A)** Schematic of heterochronic whole bone marrow (BM) transplantation (BMT) paradigm. Fresh whole bone marrow from UBC::EGFP donors (young or aged) was transplanted into busulfan-conditioned young (3 mo) or aged (18 mo) recipients. Mice were then treated for 14 days with PLX. Following 12 weeks of recovery, brains were harvested for imaging and bulk RNA-seq. **(B)** Experimental workflow: hemispheres were split for imaging or enzymatic dissociation and FACS isolation of GFP+CD45+CD11b+ cells for bulk RNA-seq. **(C)** Percent EGFP+ cells among brain CD45+CD11b+ myeloid cells quantified by flow cytometry. Mean ± SD; two-tailed *t test*. **(D)** Representative confocal images of EGFP, Iba-1, and merged EGFP/Iba-1 from young→young, young→old, old→young, and old→old groups. Scale bars: 50 μm. **(E)** Principal component analysis (PCA) of bulk RNA-seq data, colored by donor and recipient age. **(F)** Venn diagrams showing differentially expressed genes (DEGs; padj < 0.1) by environmental condition (aged vs young recipient) and overlap with other comparison groups. Arrows indicate if DEGs in respective comparison were up- or down-regulated. Upper Venn diagram: Overlap of DEGs up-regulated in (old→old vs young→young) cells, up-regulated in (young→old vs young→young) cells and down-regulated in (old→young vs old→old) cells. Lower panel vice versa. **(G)** Representative GO-term enrichment for DEGs enriched in old→old versus young→young reconstituted myeloid cells. **(H)** Heatmap of DEG’s across transplant groups, with several CAAS genes highlighted. **(I)** Bulk RNA-seq normalized counts for exemplary genes *Gpnmb*, *Ccl3*, and *Cd9* in indicated groups. Mean ± SEM. Two-sided *Wald test*. ***p < 0.001, °p < 0.1. **(J–M)** Representative image of Iba-1+ cells in the brains of young BM-transplanted (J) and aged BM-transplanted (L) young and aged mice. Images were digitally processed and converted for quantification. Quantification of Iba-1+ cell morphology in the cortex of young (K) and aged mice (M). Left columns: Total length of processes. Center columns: number of branch points. Right columns: Ratio of cell body to total cell area. Horizontal bars represent median values, n=3/group, 3 brain sections/animal were quantified. Two-tailed *t test*. ***p < 0.001, **p < 0.01. Complete DGE and GO tables are listed in Table S2.

**Figure S10.**
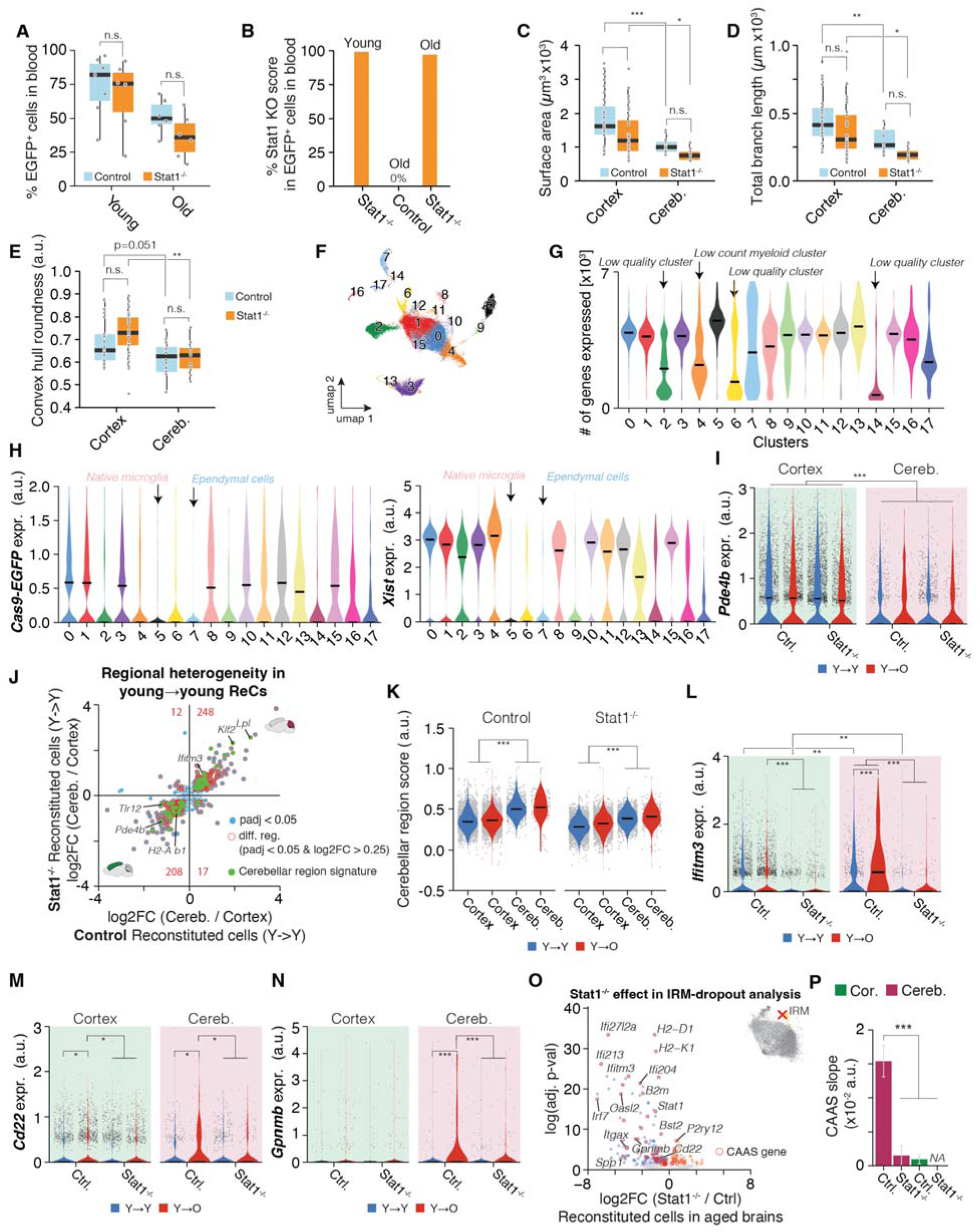
Effects of Stat1 deletion on ReC morphology, transcriptome, and region identity. **(A)** % EGFP+ cells in blood of young and aged recipients, resolved by genotype. **(B)** % KO scores for bulk-collected EGFP+ cells (10^4^ cells/sample) as collected from one animal per condition post-busulfan. Sanger sequencing data from PCR amplicons containing all three sgRNA target loci within the Stat1 gene (Editco-designed multimers) were compared to WT using Editco’s ICE analysis tool. **(C-E)** Surface area (C), total branch length (D), and convex hull roundness (E) quantifications of Iba-1+ cell morphology in the cortex of aged ReCs in cortex and cerebellum from control (Ctrl) and Stat1^-/-^ chimeras. quantified from Iba-1+ reconstructions. *p < 0.05, **p < 0.01, ***p < 0.001, n.s. = not significant; (n = 3 mice per group, 10 to 15 cells per animal and region were quantified). Two-tailed *t test*.; **p < 0.01, ***p < 0.001. **(F)** UMAP embedding of all GFP+CD45+ cells, annotated by cluster identity. **(G)** Number of genes detected per cell in each cluster. Low quality clusters and a low count myeloid cell cluster were identified and censored. **(H)** Cas9–EGFP and *Xist* expression across clusters, with native microglia and ependymal cell clusters indicated. **(I)** Expression of region-dependent gene *Pde4b* in control and Stat1^-/-^ ReCs from cortex and cerebellum. Median is indicated. *MAST*, Benjamini-Hochberg correction. ***p < 0.001. **(J)** Scatter plot comparing log2FC(cerebellum/cortex) between Stat1^-/-^ and control ReCs (both young→young). Genes with padj < 0.05 and |log2FC| > 0.25 are highlighted; cerebellar region signature genes (compare Figures 1M, S3L) are indicated in green. **(K)** Cerebellar-region scores for ReCs in cortex and cerebellum, resolved by genotype and donor–recipient age combination. ***p < 0.001 by two-sided *Wilcoxon test* **(L)** *Ifitm3* expression in control and Stat1^-/-^ ReCs from cortex and cerebellum. **(M–N)** Expression of *Cd22* **(M)** and *Gpnmb* **(N)** in ReCs, resolved by genotype and donor–recipient age combination. *MAST*, Benjamini-Hochberg correction. *p < 0.05, **p < 0.01, ***p < 0.001, ****p < 0.0001. **(O)** Volcano plot showing Stat1^-/-^ effect on gene expression in ReCs in aged brains after excluding all cells belonging to the IRM; CAAS genes are labeled. **(P)** CAAS slopes for ReCs from cortex and cerebellum in Ctrl and Stat1^-/-^chimeras after excluding all cells belonging to the IRM. Mean ± 95% confidence intervals. Two-sided *Tukey’s HSD test*. The highest (least significant) p value is indicated. ***p < 0.001. N.A., not applicable where CAAS exhibited no association with age.

**Figure S11.**
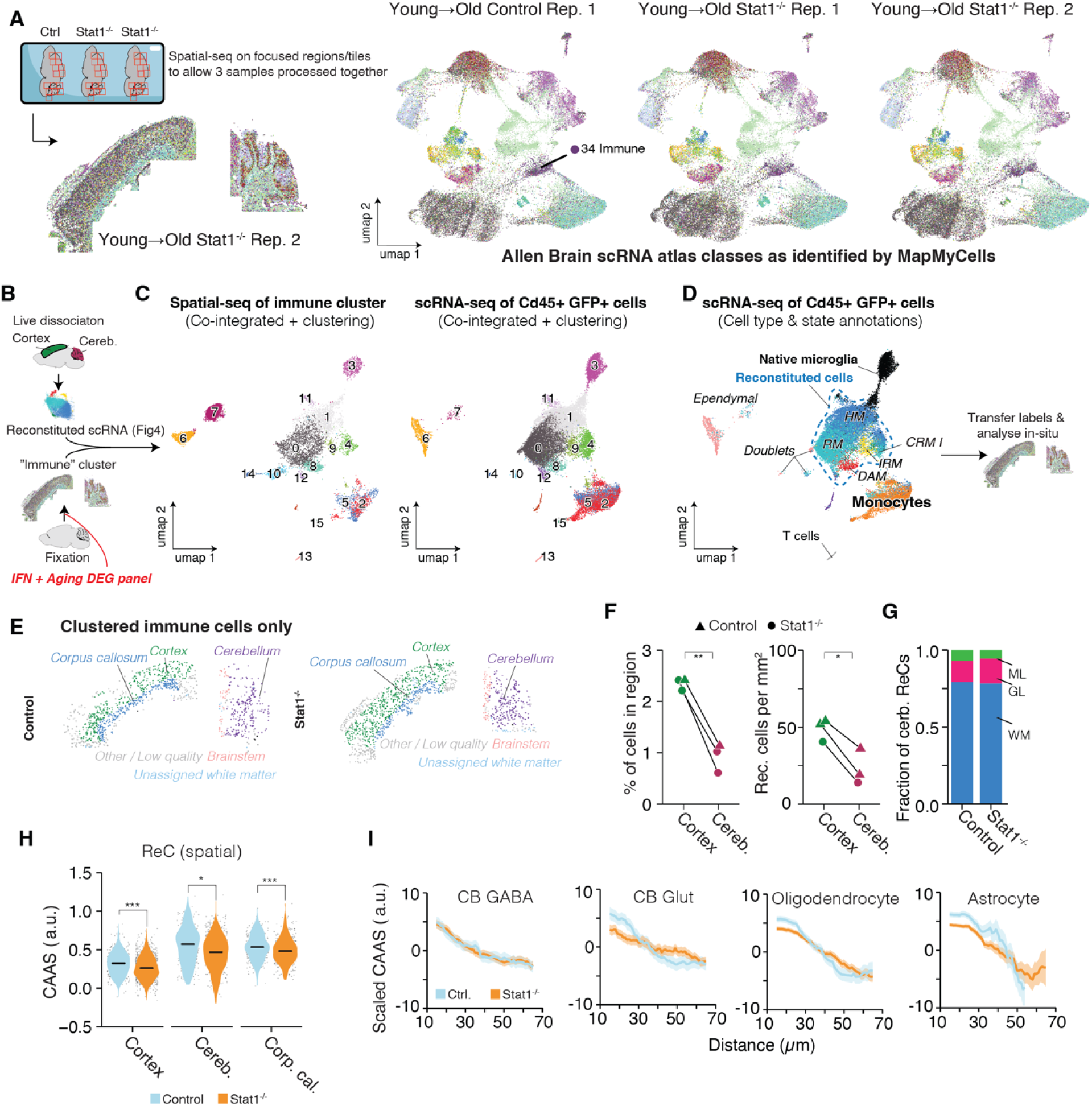
Spatial-seq mapping and in situ analysis of immune cluster states in Stat1-deficient and control chimeras. **(A)** CosMx was run using the custom IFN + aging DEG panel. Representative CosMx spatial maps from young→old control (Ctrl.) and young→old Stat1^-/-^ chimeras. Brain slices were processed together (3 slices in total) on the same CosMx slide but only imaged on cortex and cerebellum to enable total imaging times comparable to whole slice quantifications (for two slices) as performed in Figures S4 and S7. On the right: UMAP representation of cells annotated by Allen Brain Atlas reference mapping using MapMyCells. Immune cluster was utilized for integration with scRNA-seq. **(B)** Schematic of integration workflow: immune cluster from spatial-seq co-integrated with scRNA-seq of GFP+CD45+ cells (see Figure 4), clustered, and annotated. **(C)** UMAP of immune cluster cells from spatial-seq after co-integration and clustering with GFP+CD45+ scRNA-seq data. **(D)** UMAP of co-integrated GFP+CD45+ cells annotated by cell type/state. Labels were transferred back to spatial-seq data for in situ analysis. **(E)** Regional annotation of clustered ReCs in cerebellum, cortex and corpus callosum from control and Stat1^-/-^ chimeras. **(F)** Left, fraction of ReCs in cortex and cerebellum of young→old Ctrl (▲) and Stat1^-/-^ (●) chimeras. Right, density of ReCs per mm². **p < 0.01. Paired two-tailed *t test*. **p < 0.01, *p < 0.05. **(G)** Relative fraction of cerebellar Ctrl and Stat1^-/-^ ReCs in ML, GL and WM. **(H)** Violin plots of in-situ quantified CAAS in Ctrl and Stat1^-/-^ ReCs. *Two-sided Wilcoxon rank-sum test*. ***p < 0.001. **(I)** Distance-dependent CAAS profiles for ReCs relative to cerebellar (CB) GABAergic neurons, CB glutamatergic neurons (CB Glut), oligodendrocytes (Oligo.) and astrocytes. A constant that equals to the mean score across all distances is subtracted from each curve. Data are presented as mean (solid line) ± SEM (shade). Complete DGE and GO tables are listed in Table S2.

**Figure S12.**
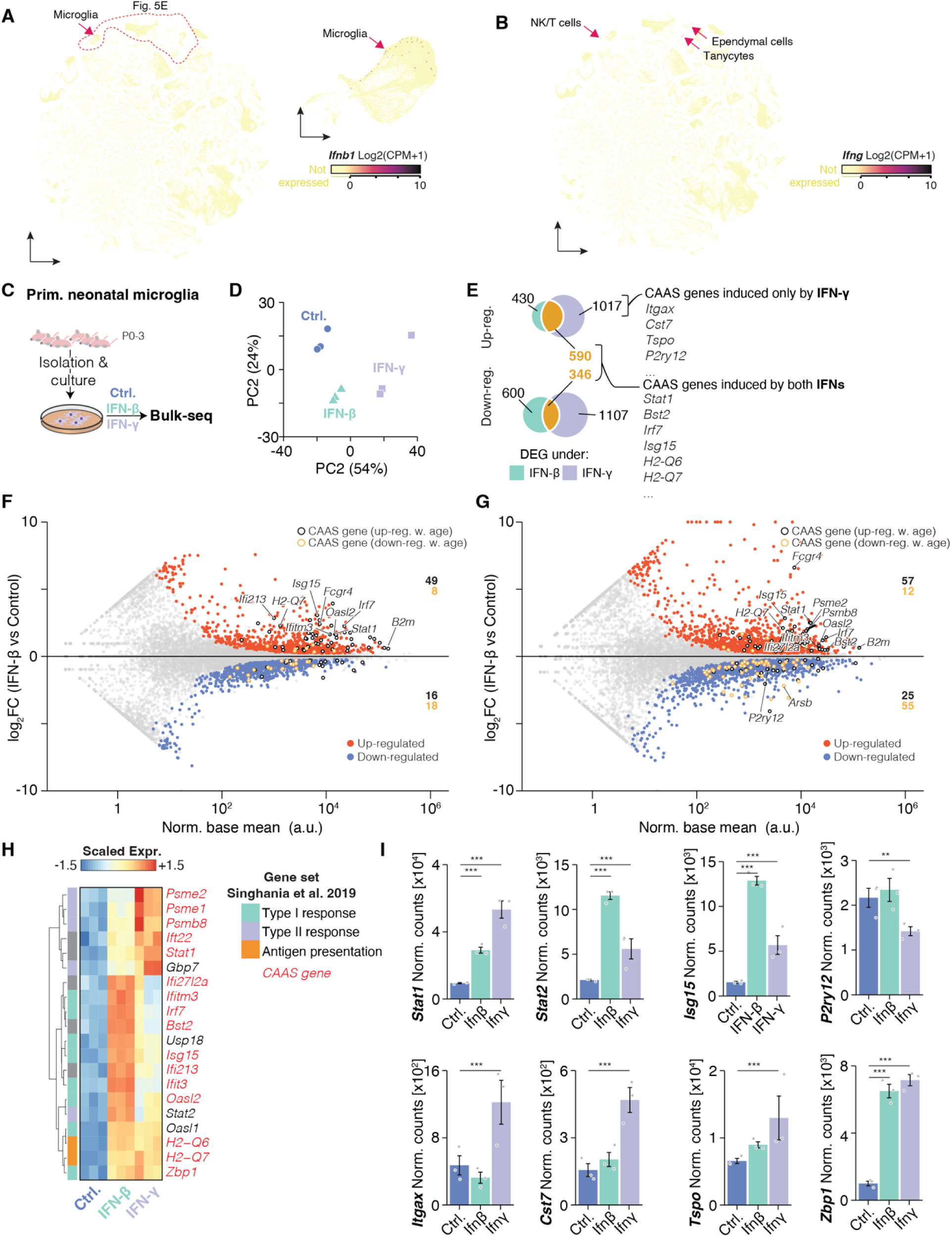
Transcriptional responses of primary microglia to IFN-β and IFN-γ stimulation. (A–B) Visualization of *Ifnb1* **(A)** and *Ifng* **(B)** expression across cell types in the Allen Brain Cell Atlas^88^ visualization tool. *Ifnb1* expression is enriched in microglia; *Ifng* expression is detected in NK cells, T cells, ependymal cells, and tanycytes. Closed up visualization of relevant cell types from (B) are displayed in Figure 5E. Quantifications of *Ifnb1* and *Ifng* expressing cells are displayed in Figure 5F. **(C)** Experimental design: primary neonatal microglia were isolated from P0–3 neonates and cultured before being treated with DMSO (Ctrl.), IFN-β, or IFN-γ for 16 hours, followed by bulk RNA-seq. **(D)** Principal component analysis (PCA) of bulk RNA-seq profiles. **(E)** Venn diagrams of differentially expressed genes (DEGs; *padj < 0.1 in *Wald* test) induced under IFN-β (green) and IFN-γ (purple), indicating several exemplary CAAS genes specific to IFN-γ (e.g., *Itgax*, *Cst7*, *Tspo*, *P2ry12*) and those induced by both IFNs (e.g., *Stat1*, *Bst2*, *Irf7*, *Isg15*, *H2-Q6*, *H2-Q7*). **(F–G)** MA plot comparing mean RNA expression (Norm. base mean) versus log2FC(treated/control) in bulk RNA-seq data for IFN-β and IFN-γ treated cells. Up- and down-regulated genes are shown in red and blue, respectively; non-significant genes are in grey; CAAS genes are labeled. The number of CAAS genes (resolved by up- or down-regulation with age) in the upper and lower halves of the plots are indicated. padj < 0.1 was considered as significant. Statistical significance was assessed using DeSEQ2;s two-sided *Wald test*. Complete DGE tables are listed in Table S3. **(H)** Heatmap of scaled expression for selected DEGs that were annotated as Type I or Type II interferon response genes according to published annotations^56^, as well as antigen presentation genes are indicated. Selected CAAS genes are highlighted in red. **(I)** Normalized counts for selected genes shown in (H) (*Stat1*, *Stat2*, *Isg15*, *P2ry12*, *Itgax*, *Cst7*, *Tspo*, Zbp1) across stimulation conditions. Mean ± SEM. Two-sided *Wald test*. ***p < 0.001, **p < 0.01.

**Figure S13.**
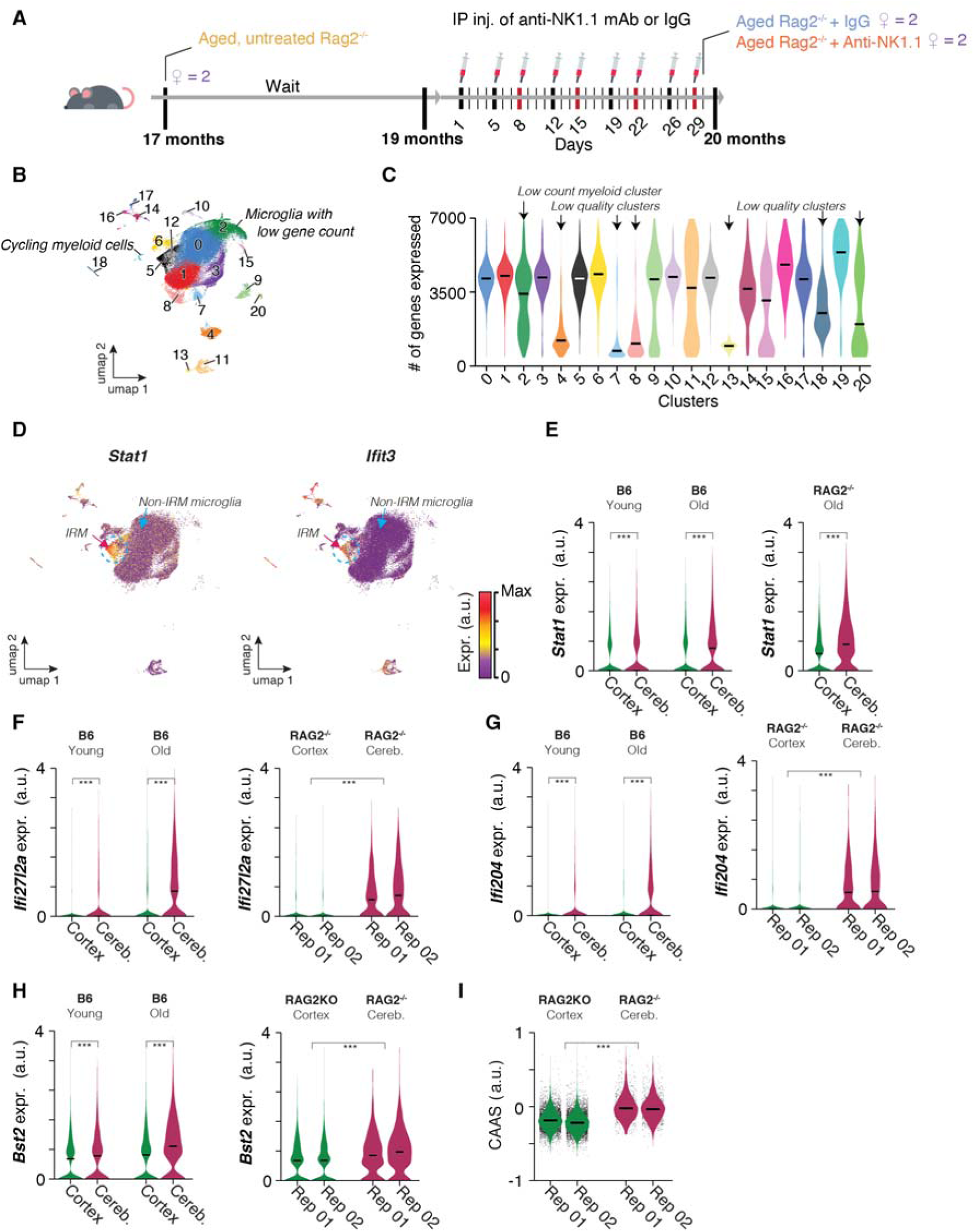
Effects of genetic ablation of T and B cells on cerebellar microglia in aged mice. **(A)** Schematic of Rag2^-/-^ experiments: female Rag2^-/-^ mice were aged in-house and microglia were isolated at 17 months of age (‘untreated’ group). The remaining mice of the same cohort received intraperitoneal injections of anti-NK1.1 monoclonal antibody or IgG control from 19 to 20 months of age (injections of 100 µg per mouse per day in 5 days-3 days-5 days-3 days intervals; red bars indicate weeks). **(B)** UMAP embedding of CD45+ cells from aged Rag2^-/-^ cerebellum and cortex, annotated by cluster identity; microglia with low gene counts and cycling myeloid cells are indicated. **(C)** Number of genes detected per cell in each cluster. Low quality clusters and a low count myeloid cell cluster were identified and censored. **(D)** UMAPs showing expression of IRM marker genes *Stat1* and *Ifit3* across microglial clusters, highlighting interferon-response microglia (IRM). IRMs are present and detectable in Rag2^-/-^ animals. **(E–H)** Expression of selected CAAS genes which are part of the interferon response: *Stat1* **(E)**, *Ifi27l2a* **(F)**, *Ifi204* **(G)**, and *Bst2* **(H)** in cortex and cerebellum from young and aged WT B6 and aged, untreated Rag2^-/-^. Rag2^-/-^ data is resolved by biological replicates. Center line = median. *MAST*, Benjamini-Hochberg correction. *p < 0.05, **p < 0.01, ***p < 0.001. **(I)** CAAS distributions in cortex and cerebellum for aged Rag2^-/-^ mice. ***p < 0.001 by two-sided *Wilcoxon test*

**Figure S14.**
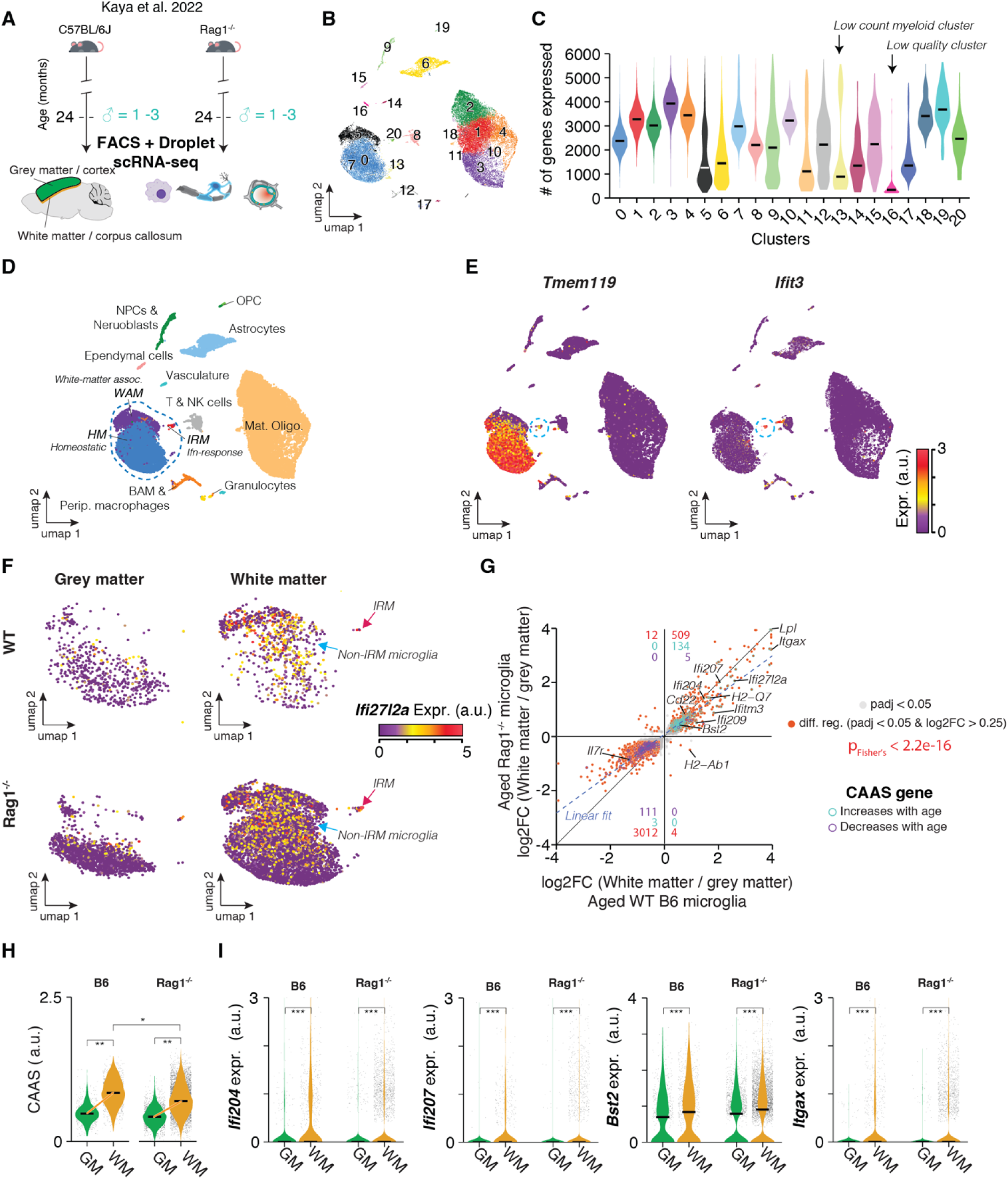
Meta-analysis of effects of genetic ablation of T and B cells on microglia in aged Rag1^-/-^ mice. **(A)** Schematic of published FACS-sorted droplet-based scRNA-seq profiling of cells from C57BL/6J and Rag1^-/-^ mice of cortical grey and subcortical white matter^23^ (GM, WM; n = 8,960 microglia; n = 1-3 males /genotype and tissue). **(B)** UMAP embedding of all cells colored by cluster identity. **(C)** Number of genes detected per cell in each cluster. Low quality clusters and a low count myeloid cell cluster were identified and censored. **(D)** UMAP embedding annotated by major cell types and states, including homeostatic microglia (HM), interferon-response microglia (IRM), white-matter–associated microglia (WAM), and other immune and non-immune cell types. **(E)** UMAP feature plots for *Tmem119* and *Ifit3* expression. **(F)** UMAPs showing white matter vs. grey matter distribution of microglial subtypes in WT and Rag1^-/-^ mice, colored by *Ifi27l2a* expression. Blue arrow indicates strong expression of *Ifi27l2a* in non-IRM microglia, similar to Figures S6C. **(G)** Scatter plot comparing per-gene log2FC(white matter/grey matter) between aged Rag1^-/-^ and aged WT microglia; CAAS genes are highlighted. Numbers in each quadrant indicate DEGs (padj<0.05, |log2FC| > 0.25). Number of DEGs that overlap with up- (turquoise) or down-regulated (purple) CAAS genes are indicated in each quadrant. Significance of overlap of DEGs in WT and Rag1^-/-^ was quantified by *Fisher’s exact test*. Complete DGE tables are listed in Table S3. **(H)** CAAS in microglia from grey vs. white matter for each genotype. Center line = median. Significance were tested via two-tailed *t test* on per-replicate median of score. *p < 0.05, **p < 0.01. **(I)** Expression of selected CAAS genes: *Ifi204*, *Ifi207*, *Bst2* and *Itgax*. Center line = median. *MAST*, Benjamini-Hochberg correction. *p < 0.05, **p < 0.01, ***p < 0.001.

**Figure S15.**
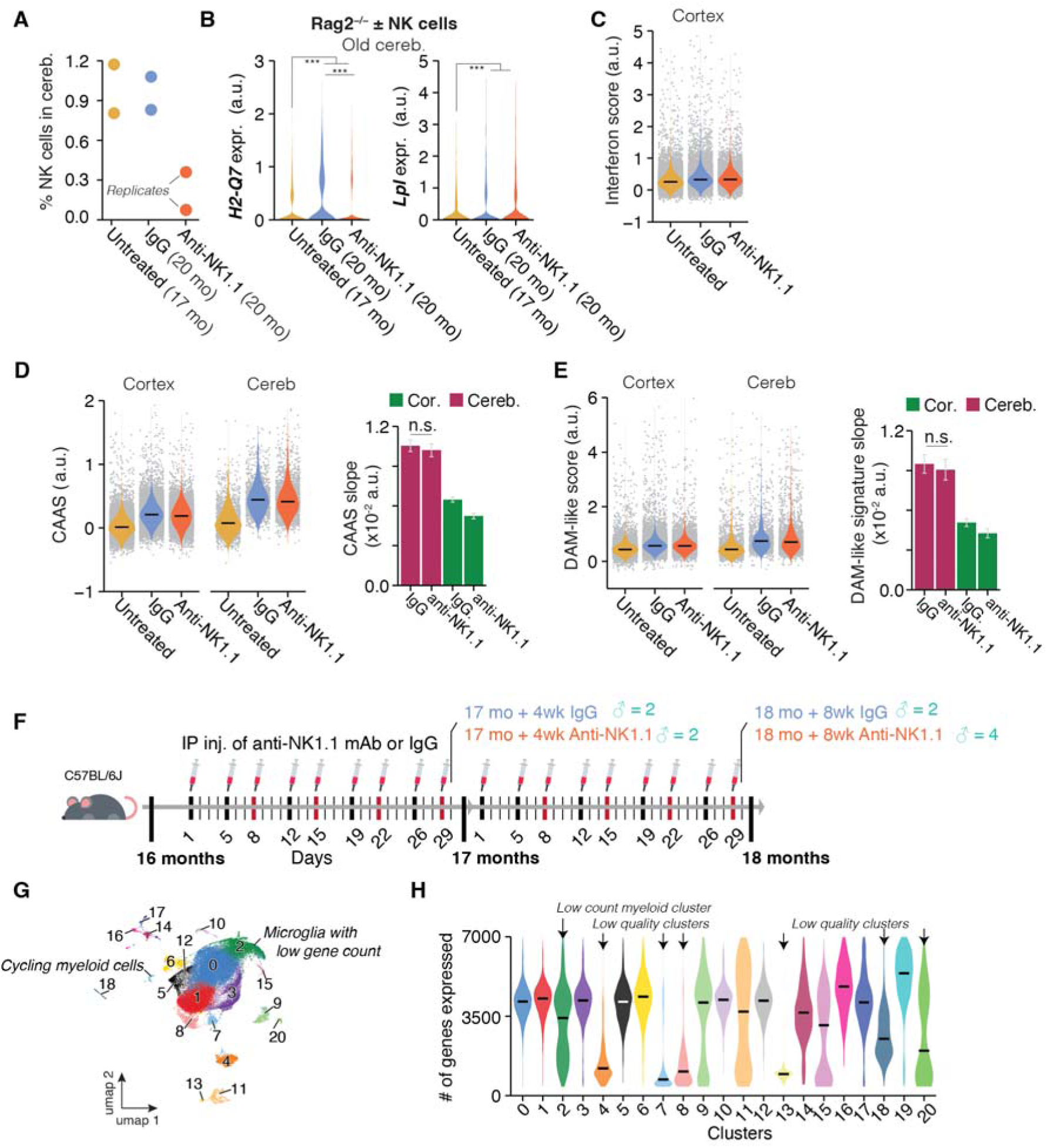
NK cell depletion in aged cerebellum and effects on interferon and DAM-like signatures. **(A)** Percentage of NK cells in cerebellum from individual replicates of untreated (17 months old), IgG-treated (20 months old), and anti-NK1.1–treated (20 months old) aged Rag2^-/-^ mice. **(B)** Expression of *H2-Q7* and *Lpl* in cerebellar microglia from the same groups. Center line = median. *MAST*, Benjamini-Hochberg correction. *p < 0.05, **p < 0.01, ***p < 0.001. **(C)** Interferon signature scores in cortical microglia following treatments. Statistical analysis of signature slope is represented in Figure 6H. **(D)** CAAS distributions in cortical and cerebellar microglia (left) and slope quantification (right). **(E)** DAM-like signature scores in cortical and cerebellar microglia (left) and slope quantification (right). For (D) and (E), Center line = median; n.s., not significant by two-sided *Tukey’s HSD test*. Untreated samples were used as reference for both IgG and anti-NK1.1 treated mice. **(F)** Experimental design: aged C57BL/6J males received intraperitoneal injections of anti-NK1.1 monoclonal antibody or IgG control for 4 or 8 weeks starting at 16 months of age (injections of 100 µg per mouse per day in 5 days-3 days-5 days-3 days intervals; red bars indicate weeks). **(G)** UMAP embedding of all cells colored by cluster identity; microglia with low gene counts indicated. **(H)** Number of genes detected per cell across clusters; arrows indicate low-count or low-quality clusters.

## Lead Contact

Requests for resources and reagents should be directed to the lead contact, Oliver Hahn.

## Materials availability

This study did not generate new unique reagents.

## Data and Code Availability

- The sequencing data have been deposited at Gene Expression Omnibus repository and are publicly available as of the date of publication. Publicly available datasets were obtained from the following repositories: BioProject PRJNA600501 and Gene Expression Omnibus GSE306641, GSE306331, GSE306335, GSE306344, GSE306351, GSE306354. CosMx data was deposited and is accessible via the GitHub repository indicated below.
- All original code has been deposited at
- https://github.com/calico/Heterochronic_myeloidCellTransplantation and is publicly available as of the date of publication.
- Any additional information required to reanalyze the data reported in this paper is available from the lead contact upon request.

### EXPERIMENTAL MODEL AND STUDY PARTICIPANT DETAILS

#### Animal husbandry and organ collection

Unless stated otherwise, young and aged C57BL/6J (000664, Jackson Laboratory) were shipped from Jackson Laboratory. Female Rosa26-Cas9 knock-in mice (026179, Jackson Laboratory) for generation of eHSC pools were shipped from Jackson Laboratory at 2 months of age. Mice of the same cage undergoing reconstitution with edited eHSCs received either control or Stat1^-/-^ eHSCs to average out cage- or litter-related batch effects. All mice were housed in cages of 3-5 mice at the Calico Life Sciences LLC Laboratory Animal Resources (Calico LAR) facility under a 12 h/12 h light/dark cycle at 67–73 °F and provided with food and water ad libitum. Mice were housed for at least 2 weeks before commencing experiments. Mice older than 18 months were housed at the Calico LAR beginning at 18 months. For environmental enrichment, mice had constant access to nesting material and housing.

Rag2^-/-^ (008449, Jackson Laboratory) mice were bred and aged in-house at Calcio LAR in a dedicated room for immunocompromised subjects at sterile barrier level. Administration of anti-NK1.1 mAb and IgG controls injections also occurred in the same location.

Unless stated otherwise, mice were anesthetized via isoflurane and ∼700ul blood was drawn via cardiac puncture before transcardial perfusion with 20 ml cold PBS. The brain was immediately removed and submerged in 4 °C cold PBS for 15 seconds before separation of hemispheres and downstream processing. Takedowns of any experimental cohort were conducted between 9:00am-11:00am to minimize shifts in circadian rhythms. Age and treatment groups were rotated through over the duration of any takedown to average out the impact of takedown time and day. All animal care and procedures complied with the Animal Welfare Act and were in accordance with institutional guidelines and approved by the Institutional Animal Care and Use Committee of Calico Life Sciences LLC.

Husbandry of, and experiments with, animals involving bone marrow transplantation (Figure S9) were conducted via animal procedures approved by the administrative panel on laboratory animal care at Stanford University. All mice were age and sex matched (except for breeding pairs) and group-housed (up to 5 mice per cage) under a 12 h/12 h light/dark cycle at 67–73 °F and provided with food and water ad libitum. Female WT C57BL/6JN mice at the age of 3 or 18 months were obtained from the National Institute on Aging rodent colony. Female 3 and 20 months old C57BL/6-Tg(UBC-GFP)30Scha/J strain (004353, Jackson Laboratory) mice, referred to as UBC::EGFP mice, were prepared and aged in-house. For brain processing, anesthetized mice (100 mg/kg ketamine and 10 mg/kg xylazine, IP) were perfused through the left ventricle with ice-cold PBS without Ca2^+^ and Mg2^+^. Brains were removed and cut into two pieces along the midline.

C;129S4-*Rag2*^tm^^1.1Flv^ *Csf1*^tm^^1(CSF^^1)Flv^ *Il2rg^t^*^m^^1.1Flv/J^ mice (017708, Jackson Laboratory; referred to as Rag2^-/-^γc^-/-^ animals for brevity) were obtained and a breeding colony was established at the Animal Core Facility of the Broad Institute of MIT and Harvard. All animals used for sequencing experiments were bred, born, and aged within this facility. Mice were group-housed under a 12 h/12 h light/dark cycle at 67–73 °F and provided with food and water ad libitum. All procedures were reviewed and approved by the Broad Institute Institutional Animal Care and Use Committee and were conducted in accordance with relevant guidelines and regulations. Male animals were anesthetized using 2.5% v/v Avertin and transcardially perfused with cold HBSS until the liver was fully decolorized. Whole brains were extracted and snap-frozen by submerging in liquid nitrogen-cooled isopentane. Brains were stored at -80 °C until further processing

### METHOD DETAILS

#### Isolation and expansion of HSC progenitor cells

HSCs were isolated and cultured based on a previously published protocol^35, 89^. Mouse tibia and femur bones from 3-4 adult female mice were placed into ice-cold HBSS (Gibco, catalog #14175103), and muscle and connective tissue was carefully removed. Bones were then crushed in an agate mortar and pestle (Fisherbrand, catalog # FB970K) to liberate bone marrow cells, and HBSS with 0.5 mg/ml DNase (Worthington Biochemical Corporation, catalog #NC9199796) was added to the resulting cell suspension. Cell suspension was passed through a 40 µm cell strainer, and any remaining clumps of cells were gently pressed through the filter. Cells were then spun down at 440xg for 5 minutes and incubated with 10 ml of ACK lysis buffer (Quality Biological, catalog #50983219) for 15 minutes at room temperature. 20 ml of HBSS were added, and the cells were spun down at 440xg for 5 minutes. c-KIT-positive cells were isolated with magnetic beads (Miltenyi Biotec, catalog #130-091-224) and LS columns (Miltenyi Biotec, catalog # 130-042-401) following manufacturer’s instructions. The resulting HSCs were resuspended in medium composed of F12 basal medium (Gibco, catalog # 11765062), 1% ITSX (Gibco, catalog # 51500056), 10 mM HEPES (Corning, catalog # MT25060CI), 1% P/S (Gibco, catalog # 15140122), 1% Glutamax (Gibco, catalog # 35050061), 0.1% PVA (Sigma-Aldrich, catalog # P8136-250G), 100 ng/ml mouse TPO (StemCell Technology, catalog # 502269647), 10 ng/ml mouse SCF (StemCell Technology, Catalog # 78064) at 5 million cells per well of a cell culture-grade six well plate. The cells were maintained in a hypoxic incubator (37^0^C, 5% CO_2_, 5% O_2_) and split at 1:3 ratio every week.

#### CRISPR editing of ex vivo eHSC progenitor cells

Cells were nucleofected with either the murine Stat1 Gene Knockout Kit or NTC1 control sgRNA (EditCo) as follows: 8 x 10^6^ cells were spun down (250rcf / 5min. / Room temp) and the pellet was resuspended in 200µl of nucleofection buffer P3 (Lonza Catalog #: V4XP-3024) containing 200pmol sgRNA and an end concentration of 4µM Alt-R™ Cas9 Electroporation Enhancer (IDT catalog #10007805). Cell suspensions were split among 2 Nucleocuvette^®^ Vessels in 2 x 100µl aliquots and nucleofected on the 4D-Nucleofector® X Unit (Lonza Catalog #: AAF-1003X) using program DS-130. After nucleofection, the cells in each cuvette were resuspended with 900µl of complete HSC media and split among 2 wells of a 6-well plate. 2.5 mls of media was added to each well and the plates were placed in a hypoxic incubator (37°C / 3%O_2_ / 5% CO_2_) until ready for reconstitution. We confirmed knockout efficiencies in eHSCs in addition to collecting 10,000 EGFP+ blood cells post-reconstitution from young and aged animals that received Stat1^-/-^ cells (Figure S10B). To determine KO efficiency, Sanger sequencing data from PCR amplicons containing all three sgRNA target loci within the Stat1 gene (Editco-designed multimers) were compared to EGFP+ Control cells using Editco’s ICE analysis tool.

#### In vivo reconstitution of brain myeloid cells (eHSC-based)

Unless otherwise noted, we used male, 3-4 months (young) or 18-19 months (aged) old C57BL/6J mice as recipients. Recipient mice were placed on drug-compatible chow (D11112201, ResearchDiets) for one week prior to the start of experiments. Recipient mice were conditioned with busulfan (Sigma, B1170000; intraperitoneal injections, once daily for 4 days). For the reconstitution of young and aged mice with unedited eHSCs derived from Rosa26-Cas9 donors (Figures 1-3), young mice received a busulfan dose of 25 mg/kg body weight while aged animals received either a 25 mg/kg or 20 mg/kg body weight dosage to assess the impact of chimerism on age-related effects in ReCs. For the reconstitution of young and aged mice with eHSCs edited via control or Stat1-targeting gRNAs (Figure 4), mice of both age groups received a busulfan dose of 25 mg/kg body weight. Due to temporary supply chain issues limiting availability of busulfan from Sigma, mice were instead injected with pre-formulated busulfan (McKesson, 1207842). This may explain the slightly lower chimerisms achieved in aged individuals (Figure S10A). Regardless of dosage regimen or busulfan source, aged mice with total body weights over 40g received no more than a maximum dose of 1mg/day.

Mice were housed on heating pads and provided supportive hydrogel or moistened chow during chemotherapy. At final busulfan dose, eHSCs were washed and resuspended in sterile saline and administered by a single retro-orbital intravenous injection (100–120 µl containing 2×10⁴–5×10⁶ viable cells) under brief isoflurane anesthesia (2–2.5% in oxygen); one drop of 0.5% proparacaine ophthalmic solution was applied at the time of injection. Supportive measures were provided for an additional seven days post-busulfan.

To enable donor-derived myeloid cells to populate the CNS, animals were switched 14 days after eHSC infusion to a chow containing the CSF-1R inhibitor PLX5622 (1,200 ppm formulated by ResearchDiets in OpenSource Diet base, D11112201) for 14 days, after which they were returned to base diet for 4-5 weeks prior to tissue collection. Peripheral engraftment was monitored at interim (post-busulfan) and pre-terminal (two days prior takedown) time points by tail-nick blood collection. For mice receiving control or Stat1^-/-^ cells, cage-mate mice of the same recipient age were randomly assigned to their respective cell populations and group-housed (up to 5 per cage).

#### In vivo reconstitution of brain myeloid cells (bone marrow-based)

We used male, 3-4 months (young) or 18-19 months (aged) old C57BL/6JN mice as recipients. Recipient mice were conditioned with busulfan (Sigma, B1170000; intraperitoneal injections, once daily for 5 days), at 25 mg/kg body weight per day. Aged mice with total body weights over 40g received no more than a maximum dose of 1mg/day. On day 6, bone marrow cells (BMCs) were isolated from the tibias and femurs of UBC-GFP mice. Isolated BMCs were resuspended in 1x red blood cell lysis buffer (e-bioscience, 00-4300-54) and incubated for 20 minutes at room temperature. BMCs were passed through a 70 µm cell strainer, washed by applying 10 ml of ice-cold PBS, and then resuspended in fresh PBS. BMCs (5x10^6^) in 200 µl PBS were injected into the retro-orbital sinus of the pre-conditioned recipients. 28 days post-conditioning, recipient mice were treated with PLX5622 (1,200 ppm formulated into AIN-76A standard chow by ResearchDiets) or a control diet for 10 days. 12 weeks post-transplantation, mice were euthanised for analysis of myeloid cells and immunofluorescence analysis.

#### Blood chimerism analysis and flow cytometry

Blood chimerisms of reconstituted mice were analyzed approximately one week after eHSC injection and confirmed one week before takedown. Mice were fixed within cylindrical restrainers. Tail nick blood collections using sterile scalpels resulted in ∼20ul blood which was collected in tubes coated with of 250 mM EDTA (Thermo Fisher Scientific, 15575020) and kept on ice until processing. Blood samples were incubated with 2 ml of ACK lysis buffer (Quality Biological, catalog #50983219) for 10 minutes at room temperature under mild shaking. Lysis was quenched by adding 10 ml PBS. Cells were pelleted and resuspended in 2 ml fresh PBS and filtered through a 35-μm cell strainer into a 5 ml round bottom FACS tube (Corning, 352235). Cells were analyzed on a Wolf G2 Flow Cytometer (Nanocellect). Cells were gated on size and chimerism was identified via GFP expression. We note that the documented^36^ relatively weak expression of EGFP from the NLS-Cas9-NLS-P2A-EGFP transgene means that we likely did not count (and sort) several true positives while also collecting erroneously non-chimeric cells. FlowJo (version 10.10.0) was used for data analysis.

#### Anti-NK1.1 mAb treatment

To deplete NK cells in vivo, we followed previously published protocols to deplete NK cells in aged brains^63^. In brief, we treated animals with InVivoPlus anti-mouse NK1.1 mAB (BP0036, Bio X Cell) or InVivoPlus mouse IgG2a isotype control (BP0085, Bio X Cell). mABs for injection were prepared in InVivoPure pH 7.0 Dilution Buffer (IP0070, Bio X Cell). 19 months-old female Rag2^-/-^ or 16 months-old male C57BL/6J mice received anti-NK1.1 mAb or IgG control at a dosage of 100 µg per mouse per day in 5 days-3 days-5 days-3 days intervals until the end of experiments. Depletion of NK1.1+ cells was confirmed by flow cytometry and was consistently >90%.

#### Splenocyte analysis and flow cytometry

Spleens were surgically removed from mice, cut into smaller pieces, and then mechanically dissociated using syringe plungers and cell strainers. The resulting cell suspensions were filtered through 70µm cell strainers, centrifuged, and resuspended in ACK lysis buffer for 5 minutes at room temperature. After lysis, cells were diluted ten-fold with PBS and passed through 70µm cell strainers to eliminate debris. Cell counts and viability were determined using a Vicell XR Cell Viability Analyzer (Beckman Coulter). Cells were then resuspended in EasySep buffer (Stemcell Technologies, Cat.# 20144) for antibody staining and subsequent flow cytometry analysis, which was conducted using an LSRFortessa Flow Cytometer. Dead cells were identified with a LIVE/DEAD™ Fixable Near-IR Dead Cell Stain Kit. Host and engrafted cells were distinguished based on GFP expression. The antibodies used for flow cytometry included: CD45 (BUV396, clone 30-F11, BD, Cat.# 564279, 1:400), B220 (BUV737, clone RA3-6B2, BD, Cat.# 612838, 1:200), CD3e (PerCP Cy5.5, clone 17A2, Biolegend, Cat.# 100218, 1:200), NK1.1 (BV711, clone PK136, Biolegend, Cat.# 108745, 1:200), Ly6G (BV605, clone RB6-8C5, Biolegend, Cat.#, 1:200), and CD11b (BV786, clone M1/70, ThermoFisher, Cat.# 417-0112-82, 1:200). FlowJo (version 10.10.0) was used for data analysis.

#### Isolation and scRNA-seq of CD45+ and CD45+/EGFP+ cells from cortex and cerebellum

Unless otherwise indicated, immune cells and reconstituted immune cells were isolated by an ice-cold dounce homogenize method as previously described^90^. Mice were perfused with 20 ml 1x PBS and split into hemispheres. From one hemisphere, cortex, and cerebellum were dissected. Brain tissues were chopped and homogenized using a dounce homogenizer in 2 ml of cold medium A (HBSS, 15 mM HEPES, 0.5% glucose and 1:500 DNase I), filtered through a 70-µm cell strainer, rinsed with 6 ml medium A and centrifuged at 400 × g for 5 min. Myelin removal beads and columns from MACS separation system (Miltenyi, beads #130-096-433, columns #130-042-901) were used to remove myelin by incubating 50 µl beads with cells in 450 µl MACS buffer (1x PBS, 2 mM EDTA and 0.5% BSA). After myelin removal, cells were resuspended in the FACS buffer (1x PBS, 2 mM EDTA, 1% FBS and 25 mM HEPES). The samples were blocked with FC (CD16/CD32, BD, 553141, 1/100) for 5 min on ice and then stained with anti-CD45-Alexa Fluor 594 (Clone 30-F11, BD biosciences) and anti-CD11b-PE-Fire 810 (clone M1/70, BioLegend) antibodies for 30 min on ice, followed by adding 3 ml FACS buffer and centrifuged at 400 × g for 5 min. Dead cells were labeled by staining with DRAQ7 Dye (1:1,000, Thermo Fisher, D15106) in the final FACS buffer.

We note that the documented^36^ relatively weak expression of EGFP from the NLS-Cas9-NLS-P2A-EGFP transgene likely resulted in an underestimation of true positives and the inclusion of erroneously non-chimeric cells.

For CITE-seq (Figure 1B), TotalSeq™-B Mouse Universal Cocktail, V1.0 (BioLegend, 199902) was resuspended and prepared as described in the manufacturer’s protocol. Cell elutions post-myelin cleanup were spun down and resuspended in 25 µl FcR block (provided with kit) for 5 min on ice and then stained with 25 µl reconstituted cocktail for 30 min on ice. Cells were washed once with 3 ml FACS buffer and subsequently stained with anti-CD45-Alexa Fluor 594 (Clone 30-F11, BD biosciences) and anti-CD11b-PE-Fire 810 (clone M1/70, BioLegend) antibodies for 30 min on ice, before continuing the cell isolation and sorting workflow as described above.

In order to maximize information yield and minimize scRNA-seq reagent costs, we pooled cortical or cerebellar material from biological replicates post-myelin cleanup for the following experiments: 1) scRNA-seq + CITE-seq of young aged microglia (pooled n=3 per tissue and age group; Figure 1B) 2) scRNA-seq of control or STAT1^-/-^ reconstituted cells (pooled n=2 per genotype, age and tissue group. Material of mice with similar chimerisms in blood were pooled; Figure 4A). Pooled samples were stained and processed as described above.

CD45+ (non-reconstituted mice; Figures 1B, 6B and 6J) or CD45+/EGFP+ (reconstituted mice; Figures 1K and 4D) cells were analyzed and collected on a WOLF G2 Flow Cytometer (Nanocellect) into 1.5 ml DNA lo-bind tubes (Eppendorf, 022431021) containing 1 ml buffer mix with PBS, UltraPure BSA (Thermo Fisher, AM2618), and RNase inhibitor (Takara, 2313B). Collected cells were centrifuged at 400 x g for 5 minutes at 4°C with break 2. Supernatant was removed leaving ∼25 µl suspended cells.

Reagents of the Chromium Single Cell 3’ GEM & Gel Bead Kit v3.1 (10X Genomics, 1000121; Figures 1B,K) or Chromium GEM-X Single Cell 3’ v4 Gene Expression GEM & Gel Bead Kit v4 (10X Genomics, 1000693; Figures 4D, 6B,J) were thawed and prepared according to the manufacturer’s protocol. Cells and master mix solution were loaded on a standard Chromium X (10X Genomics, 1000331; Figures 1B,K) or Chromium GEM-X Single Cell 3’ Chip Kit v4 (10X Genomics, 1000690; Figures 4D, 6B,J) according to manufacturer protocols. We applied 11-13 PCR cycles to generate cDNA. Library construction was conducted using Chromium Single Cell 3’ Library Construction Kit v3 (10X Genomics, 1000689; Figures 1B,K) or Library Construction Kit C (10X Genomics, 1000121; Figures 4D, 6B,J). All reaction and quality control steps were carried out according to the manufacturer’s protocol and with recommended reagents, consumables, and instruments. We chose 11 PCR cycles for library generation. Quality control of cDNA and libraries was conducted using a Bioanalyzer (Agilent). Illumina sequencing of the resulting libraries was performed on an Illumina NovaSeq S4 (Illumina). Base calling, demultiplexing, and generation of FastQ files were conducted using custom in house pipelines.

#### scRNA-seq of eHSC progenitor cells

eHSCs of 2-3 months old C57BL/6J mice were cultured and expanded as described above. After 4 weeks of culture, cells from 3 independent wells were collected, washed twice with 4°C PBS and centrifuged at 400 × g for 5 min. The cell pellet was resuspended in FACS buffer (1x PBS, 2 mM EDTA, 1% FBS and 25 mM HEPES). Dead cells were excluded by staining with DRAQ7 Dye (1:1,000, Thermo Fisher, D15106) in the final FACS buffer. 40,000 live cells were collected on a WOLF G2 Flow Cytometer (Nanocellect) into 1.5 ml DNA lo-bind tubes (Eppendorf, 022431021) containing 1 ml buffer mix with PBS, UltraPure BSA (Thermo Fisher, AM2618), and RNase inhibitor (Takara, 2313B). Collected cells were centrifuged at 400 x g for 5 minutes at 4°C with break 2. Supernatant was removed leaving ∼25 µl suspended cells.

Collected cells were processed for scRNA-seq as described above using Chromium Single Cell 3’ GEM & Gel Bead Kit v3.1 (10X Genomics, 1000121) and Chromium Single Cell 3’ Library Construction Kit v3 (10X Genomics, 1000689), with 11 PCR cycles to generate cDNA and 11 PCR cycles for library generation. Quality control of cDNA and libraries was conducted using a Bioanalyzer (Agilent). Illumina sequencing of the resulting libraries was performed on an Illumina NovaSeq S4 (Illumina). Base calling, demultiplexing, and generation of FastQ files were conducted using custom in house pipelines.

#### Slide Preparation and Loading for CosMx

The slides were prepared following the user manual, CosMx SMI Manual Slide Preparation for RNA Assays (Fresh Frozen; MAN-10184-04), with minor modifications. Briefly, the frozen tissue blocks were sectioned with a cryostat (Leica CM3050 S) at 7 µm thickness and placed on Superfrost Plus slides (Fisherbrand; Cat No. 22-037-246). As the tissues were fixed prior to embedding, the slides were processed without additional 10% NBF fixation and then placed in a 60°C incubator for a 30-minute bake. The tissue sections were then washed, rehydrated, and underwent target retrieval in a pressure cooker. The slides were allowed to dry for at least 30 minutes but no longer than 1 hour. The tissue sections were permeabilized with a combination of proteinase K and protease A treatment following the dilution ratio listed in the manual. The fiducials were then prepared and applied to the slides. After application of the fiducials, the tissue sections were fixed with 10% NBF, blocked, and incubated overnight at 37°C in an ACD hybridization oven (PN 321710/321720) with denatured CosMx Mouse Neuroscience RNA panel with or without add-on probes. The following day, within 18 hours, the slides were washed, blocked, and stained with CosMx Mouse Neuroscience Cell Segmentation Kit (RNA), including nuclear (DAPI) and cell segmentation markers (rRNA, histone, and GFAP). The flow cells were then attached to the slides, flowed through with 2x SSC immediately after, and stored in the fridge before loading or immediately loaded into the instrument. Prior to loading the instrument, the CosMx RNA imaging tray 1000-plex was allowed to come to room temperature for at least 1 hour, and RNase Inhibitor was added immediately before loading of the imaging tray. The slides were loaded into the instrument, inverted with a pre-bleaching profile of Configuration C and a cell segmentation profile of Configuration B. With all reagents loaded following the loading manual (MAN-10161-10), the slides were scanned, and the fields of view were selected to cover either the entire section or regions of interest (Cortex and Cerebellum) for RNA profiling.

Once the runs were finished, the data were uploaded into AtoMx, and initial analysis, including quality control, normalization, PCA, and UMAP, was performed to generate a Seurat^91^ object for downstream analysis.

#### Bulk myeloid cell isolation and RNA sequencing from bone marrow transplanted animals

Myeloid cells from bone marrow transplanted mice (Figure S9) were isolated from adult mouse brains as previously described^27^. One hemisphere was immediately transferred to 4% paraformaldehyde for fixation. The other hemisphere was minced with a surgical knife and then transferred to ice-cold HBSS without Ca2^+^ and Mg2^+^. Minced hemi-brains were centrifuged at 300 x g for 2 minutes at room temperature, then enzymatically digested at 37 °C for 35 minutes using the Neural Tissue Dissociation Kit (Miltenyi Biotec, 130-092-628) according to the manufacturer’s protocol. The cell suspension was passed through a 70 µm cell strainer and washed by applying 10 ml of HBSS with Ca2^+^ and Mg2^+^ to the strainer. The cell suspension was centrifuged at 300 x g for 10 minutes at room temperature, re-suspended in 30% Percoll (GE Healthcare Lifesciences, 17089102), and then centrifuged at 700 x g for 10 minutes at 4 °C. The supernatant containing myelin was removed, and the pellet was washed with 10 ml of HBSS with Ca2^+^ and Mg2^+^. The resulting pellet was resuspended in FACS buffer and subjected to antibody staining for flow cytometry. Myeloid cells were identified using anti-CD45 (Clone 30-F11, BD biosciences) and anti-CD11b (clone M1/70, BioLegend) antibodies. Chimeric cells were further identified via the EGFP signal. Approximately 30,000 CD45+/CD11b+/EGFP+ cells per hemi-brain were identified by detecting expression of CD45, CD11b, and GFP using a FACS Aria II and then sorted into 700 µl of Trizol LS (Thermo Fisher, 10296010). Total RNA was extracted and then purified using a Qiagen RNeasy MinElute Cleanup Kit (Qiagen, 74204) according to the manufacturer’s protocols. Quality and quantity of RNA were assessed on a Bioanalyzer (Agilent), and samples with an RNA integrity number (RIN) > 8 were used for subsequent library preparation. The NEBNext Single Cell/Low Input RNA Library Prep Kit for Illumina (New England Biolabs, E6420S) was used to generate cDNA and libraries according to the manufacturer’s protocol. The libraries were indexed using the NEBNext Multiplex Oligos for Illumina (New England Biolabs, E6420S) and cleaned up using SPRIselect reagent (Beckman Coulter, B23317). The indexed libraries were pooled and sequenced on the Illumina platform. Raw sequencing reads were demultiplexed with bcl2fastq.

#### Immunofluorescence microscopy

##### For WT microglia and eHSC transplanted animals

Brains were isolated as described above, and transferred immediately to 10% formalin, and fixed overnight at room temperature, and stored at 4 °C until further processing. Brains were washed 3x with PBS for 10 minutes at RT and placed in 30% sucrose in PBS solution. Brains were cryo-embedded in a 1:1 solution of Tissue-Tek OCT (Sakura Finetek, 4583) and 30% sucrose. Sagittal sections (20-40 μm) were obtained using a cryostat (Leica CM3050 S) and mounted directly on Superfrost plus microscopy slides (Fisherbrand; 22-037-246) and stored at -80 °C until further processing. Slides were washed 3x with PBS for 10 minutes at RT then permeabilized and blocked with blocking buffer (.025% Triton X-100 (ThermoFisher; 16046-AE) and 10% normal donkey serum (Fisher Scientific; OB003001) in PBS. For NK cell staining, slides were first rinsed in PBS, then incubated at 60 °C for 1h30. Slides were transferred to prewarmed 1x Citrate Buffer (pH 6.0 10x antigen retriever; Sigma Aldrich Inc C9999-100ML) and placed in a water bath for 20 minutes (>99 °C). Slides were then left to cool for 20 minutes at RT, and washed in PBS x2. Slides were permeabilized and blocked with blocking buffer (0.3% Triton X-100, 4% Normal Goat Serum (Thermo Fisher; PCN5000) in PBS) for 1hr at RT. Slides were incubated with primary antibodies overnight at 4 °C in blocking solution. Primary antibodies: Rabbit anti-Iba1 (1:500; Fujifilm Wako; 019-19741), Guinea pig anti-Tmem119 (1:500; Synaptic Systems GmbH; 400 004), Rabbit anti-NK1.1 (1:400; Cell Signaling; 39197), Chicken anti-GFP (1:500, Abcam; ab13970-100ul). Next sections were washed 3x with PBS at RT for 10 min and incubated with respective secondaries (1:800-1000) after spinning down for 5 minutes at 4 °C. Secondary antibodies: AF488 Donkey Anti-Chicken IgG (ThermoFisher; A11039), AF488 Goat Anti-Chicken IgY (ThermoFisher; A-21449), AF488 Goat anti-Guinea Pig IgG (ThermoFisher; A-11073), AF568 Donkey Anti-Rabbit IgG (ThermoFisher; A10042), AF647 Goat Anti-Guinea Pig IgG (ThermoFisher; A-21450), AF647^+^ Goat anti-Rabbit IgG (ThermoFisher; A32733TR), AF568 Goat Anti-Chicken IgY (ThermoFisher; A11041). Sections were incubated in DAPI (1:1000; ThermoFisher; D3571) for 15 minutes, washed 3x in PBS 10 mins at RT then #1.5H coverglass slips (ThorLabs; CG15KH) were mounted with ProLong Glass Anti-fade (ThermoFisher; P36984).

##### Imaging

High-resolution confocal images were collected using a Leica Stellaris 8 upright point scanning confocal, with a white light laser tunable from 440-790 nm and HyD detectors (X, S and R). Images were acquired with a 10x/0.40 NA air objective; with a 20x/0.75 NA oil objective; 40x/ 1.25 NA oil objective at 1.8x zoom. Z-stacks were obtained at Nyquist sampling. Images were collected with dimensions of 1024 × 1024 pixels, scan speed 400 Hz, pixel dwell time 0.4 µs and line averaging of 2. High rendering of 3D images were obtained with Arivis Vision4D, Zeiss (Ultra HD, 4K: 3840x2160(16:9). For 2D images, a TIFF file was exported from Arivis and opened in Fiji/ImageJ and exported as a PNG.

##### Immunofluorescence microscopy of brain tissue from bone marrow transplanted animals

Brain tissue from bone marrow transplanted animals (Figure S9) was analyzed using Iba-1 (FUJIFILM Wako Pure Chemical, 019-19741) and GFP (Abcam, ab13970). Brains were isolated as described above, transferred immediately to 4% paraformaldehyde, and then fixed overnight at 4°C, followed by cryo-protection with 30% sucrose (in PBS) solution. Brains were then cryo-embedded with Tissue-Tek OCT compound (Sakura Finetek, 4583), and sagittal sections (40 μm) were obtained using a cryostat (CM 3050S, Leica). Sections were washed 4 times with PBS and processed for antibody staining. Brain sections were permeabilized and blocked with blocking buffer (0.3% Triton X-100 (Sigma, X-100) and 5% cosmic calf serum in PBS) for 1 hour. Sections were incubated with primary antibodies in blocking buffer either for 1 hour at room temperature or overnight at 4°C. Brain sections were washed 3 times with PBS and then incubated with AlexaFluor dye-conjugated secondary antibodies in blocking buffer for 1 hour at room temperature. Sections were washed 3 times with PBS and then incubated with DAPI in PBS for 2 minutes at room temperature. Sections were washed once with PBS and mounted on glass slides with a drop of ProLong Gold antifade reagent (Thermo Fisher, P36930). Images were obtained using a Zeiss LSM710 confocal microscope or Zeiss AxioImager motorized widefield fluorescence microscope.

#### Primary neonatal mouse microglia culture, interferon treatment and bulk RNA sequencing

Primary mouse microglia were purified from postnatal mouse brain tissue using MACS kit with CD11b magnetic beads (Miltenyi Biotec, 130-097-142) following manufacturer’s instructions. Briefly, brains from P4 mouse pups were placed in ice-cold HBSS, olfactory bulbs and meninges were removed under a dissection microscope, brain tissue was minced to ∼1mm^3^ pieces and enzymatically digested in 5 ml of 0.25% trypsin reconstituted from 2.5% trypsin (Gibco, 15090046) in HBSS for 30 mins at 37^0^C, with 0.25 ml of 10 mg/ml of Dnase (Worthington Biochemical Corporation, catalog #NC9199796) added in the last 5 minutes of dissociation. After enzymatic digestion, 0.5 ml of heat-inactivated FBS (to 10% of final concentration) was added to neutralize trypsin, and tissue was mechanically triturated using a 1 ml pipette, filtered through a 40 μm cell strainer (Corning 352340), pelleted at 400xg for 5 minutes and washed twice with HBSS. Dissociated cells were resuspended in MACS buffer (HBSS with 0.5% BSA) with addition of 0.5 mg/ml DNAse and incubated with CD11b antibody (Miltenyi Biotec, 130-097-142) for 15 minutes at 40C. After the incubation, cells were washed with 10 ml of HBSS, pelleted at 400xg for 5 minutes and loaded on LS columns (Miltenyi Biotec, 130-042-401) in 0.5 ml of MACS buffer on the magnetic stand. Cells were washed 3 times with 3 ml of HBSS, then the column was removed from the magnetic field, and microglia cells were eluted in 5 ml of HBSS. Cells were pelleted at 400xg, re-suspended in 1 ml of culture media, counted and used for plating.

For microglia cell culture, microglia were plated at 1x10^5^ cells/well in a 24 well plate in media composed of DMEM/F12 (Gibco, catalog # 21331020), 2% B27 (Gibco, catalog # 17504001), 0.5% N2 (Gibco, catalog # 17502001), 2% ITSX (Gibco, catalog # 51500056), 1% P/S (Gibco, catalog # 15140122), 1% Glutamax (Gibco, catalog # 35050061), 1% NEAA (Gibco, catalog # 11140050), 5µg/ml insulin (Sigma Aldrich, catalog # I9278), 5µg/ml cholesterol (MP Biomedicals, catalog # MP219934230) and freshly supplemented with 200 ng/mL IL-34 (Biolegend, catalog # 577608), 0.2 ng/mL TGFβ2 (Gibco, catalog # 10035B2UG), and 50 ng/mL M-CSF/CSF1 (StemCell, catalog # 78059). After 2 days in culture, the cells were treated with 2ng/ml of mouse recombinant interferon beta (R&D Systems, catalog # 8234MB010) or 2ng/ml of interferon gamma (Gibco, catalog # PMC4034) for 24 hours. After that, cells were washed 3x with PBS and flash-frozen for RNA extraction. Total RNA was extracted using the RNeasy Plus Micro Kit (Qiagen, cat. no. 74034) according to the manufacturer’s protocol. Libraries for RNA sequencing were prepared from total RNA with poly(A) selection using the NEBNext Poly(A) mRNA Magnetic Isolation Module (New England Biolabs, cat. no. E7490L) followed by library construction with the NEBNext Ultra II Directional RNA Library Prep Kit (New England Biolabs, cat. no. E7760L). Library quality and concentration were assessed using a fragment analyzer or TapeStation. Libraries were then pooled at equimolar concentrations and sequenced on either a NovaSeq 6000 or NovaSeq X system, targeting 25 million reads per sample. Raw sequencing reads were demultiplexed with bcl2fastq.

#### Brain region dissection and bulk RNA sequencing of Rag2^-/-^γc^-/-^ mice

Brain regions were dissected from frozen mouse brains through a modification of a previously developed protocol^7^. Frozen brains were sliced into 1mm thick coronal slices at -20°C using a metal brain matrix and .22mm razor blades (Ted Pella, 15045; VWR, 55411-050) and were then placed on dry ice and covered to prevent condensation. One slice at a time was placed on a metal block cooled on wet ice and 1.5mm and 2mm diameter regions of interest were dissected quickly via disposable biopsy punches (Alimed, 98PUN6-2, 98PUN6-3) from the left and right hemispheres guided by visual landmarks and the Allen Mouse Brain Atlas. The same biopsy punch was used for identical regions between left and right hemispheres, but replaced between regions and mice. We collected and processed punches from the motor cortex and cerebellum. Regions were stored at -80°C until further processing. RNA from the right hemisphere brain regions were isolated as described before^7^ using the RNeasy 96 kit (Qiagen, 74181) and a TissueLyser II (Qiagen, 85300), according to RNeasy 96 Handbook protocol “Purification of Total RNA from Animal Tissues using Spin Technology” without the optional on-plate DNase digestion. Quality control of RNA was conducted using a Bioanalyzer (Agilent). RNA-seq libraries were prepared using the NEBNext Ultra II Directional RNA Library Prep Kit with Poly-A enrichment. The indexed libraries were pooled and sequenced on the Illumina platform. Raw sequencing reads were demultiplexed with bcl2fastq.

### QUANTIFICATION AND STATISTICAL ANALYSIS

#### Single-cell RNA-seq preprocessing and integration

Cell Ranger (v.8.0.0) analysis pipelines were utilized to align reads to a custom mm10 reference genome (containing the NLS-Cas9-NLS-P2A-EGFP transgene^36^) and count barcodes/UMIs. To account for unspliced transcripts, reads mapping to pre-mRNA were counted. Only genes expressed in at least three cells were retained. 10X Genomics count matrices (Cell Ranger outputs) were loaded into Seurat (R 4.4.0)^91^. Cells with >200 and <6,000 detected genes and <10% mitochondrial UMIs were retained. Data were normalized with SCTransform, principal components were computed, and datasets were split into ‘layers’ by sample ID and integrated using Harmony^92^ as embedded in Seurat. A shared-nearest-neighbor graph and clustering were performed on the Harmony embedding (dims 1–30; resolution 0.4), followed by UMAP for visualization. Clusters were defined on the Harmony embedding and marker genes were identified per cluster using Seurat’s FindAllMarkers (positive markers; min.pct = 0.15; log2FC > 0.15) on the normalized assay. Clusters of microglia or ReCs were annotated manually based on canonical markers^30^ to the following categories: homeostatic (HM), interferon-response (IRM), ribosomal-processing (RM), cytokine response (CRM), Apoe^high^, disease-associated (DAM), border-associated macrophages, T/NK cells, granulocytes, and ependymal cells. T/NK cells were sub-clustered and annotated separately. Annotations were then mapped back to the main object. Low-quality clusters and likely doublets were optionally removed based on mixed lineage markers and QC metrics. We note that we consistently observed a cluster of myeloid cells (microglia or Rec) with unclear state but notably lower gene count (Figures 1C and 3C). These likely represent cells at the lowest end of the coverage distribution, as described previously^93^. Given that we cannot, however, distinguish between low-quality and low-coverage cells, we censored these clusters throughout our datasets. In general, if we could not annotate a cluster, we ignored the respective cells in downstream analyses. Exact code and parameters are provided in the accompanying GitHub repository.

#### Pseudotime analysis

Pseudotime inference was performed in R 4.3.1 using Monocle 3^94^ with SeuratWrappers. The Harmony-integrated Seurat object described above was converted to a Monocle cell_data_set (as.cell_data_set). To ensure the same embedding was used throughout, UMAP coordinates from the Seurat Harmony embedding were directly transferred to the Monocle object (reducedDims(cds)$UMAP <- umap.harmony). Cells were clustered on this UMAP space with cluster_cells (defaults), and a principal graph was learned using learn_graph(use_partition = FALSE, close_loop = FALSE). Cells were then ordered along the learned graph with order_cells to obtain pseudotime values. When appropriate, the trajectory root was chosen interactively from the UMAP (e.g., using Seurat’s CellSelector) to reflect the expected biological starting state; otherwise Monocle’s default root selection was used. Pseudotime visualizations were generated with plot_cells (colored by pseudotime or by cell type on the same graph). To identify genes whose expression varied along the trajectory, we applied Monocle’s Moran’s I test via graph_test(cds, neighbor_graph = “principal_graph”) and considered genes with q < 0.05 as significantly associated with pseudotime. Selected genes were displayed with plot_genes_in_pseudotime to visualize expression dynamics along the inferred trajectory.

#### Differential expression in scRNA-seq data

For single-cell analyses, raw UMI matrices were processed in Seurat (R 4.4.0). Cells were filtered and integrated as described above; all differential expression (DE) was performed on the RNA assay. DE between conditions used MAST^95^ with a hurdle model via FindMarkers (arguments: test.use=”MAST”, min.pct=0.05, logfc.threshold=0.05). P values were adjusted by Benjamini–Hochberg. Unless stated otherwise, genes were considered significant when adjusted P < 0.05 and |log2FC| ≥ 0.25 and were expressed in ≥5% of cells in at least one group.

Aging-related differential expression was run within tissue and within background (WT microglia or ReC) as follows: for WT microglia (WTB6) and reconstituted cells (ReC), cells were subset to either cortex or cerebellum, Idents were set to age (Old vs Young; Young→Old vs Young→Young), and DE was computed Old vs Young per tissue and background. To compare aging responses across backgrounds, we merged results by gene and plotted log2FC(Old/Young) from WT (x-axis) vs ReC (y-axis) for cortex and cerebellum separately (Fig. 3D,E). Genes significant in both datasets (adj. P ≤ 0.05 and |log2FC| ≥ 0.25) were highlighted; overlap directionality (Old-up/Old-down in both) defined the shared sets per region. Where indicated, overlap enrichment was assessed with Fisher’s exact test on concordant/discordant quadrants.

For CITE-seq data, we normalized in Seurat via the CLR method (margin = 2). Differential protein expression analysis was run using FindMarkers (arguments: test.use=”MAST”, min.pct=0.05, logfc.threshold=0.05). P values were adjusted by Benjamini–Hochberg. There is no known consensus on what change in CITE-seq data is considered biologically relevant. Thus, unless stated otherwise, genes were considered significant when adjusted P < 0.05 and |log2FC| ≥ 0.1 and were expressed in ≥5% of cells in at least one group.

#### DEG Gene Ontology functional enrichment

Unless stated otherwise, we performed functional enrichment analysis for DEGs using the Bioconductor package topGO^96^. To provide a coherent set of background genes for all analyses, we utilized a recent bulk RNA-seq dataset of primary, ex vivo of microglia^22^ at middle age to define a set of microglia expressed genes (defined as passing the independent filtering criterion of DEseq2^97^). This resulting 15,073 gene set was used as background for all functional enrichment analyses involving expression (bulk and single-cell RNA-seq) data, and the complete list is provided in Table S1. Top-ranked, representative Gene Ontology (GO) terms were selected and visualized using the CellPlot package. The full-length GO terms were shortened to fit into the figure format.

#### Signature construction, scoring, and slope analysis

Signature construction, scoring and analysis was conducted as described previously^7^. In brief, direction-consistent regional (Figure 1M) or age-related DEGs (Figure 3D,E); adj. P ≤ 0.05; |log2FC| ≥ 0.25) were converted into gene-set signatures. For instance, for age-accelerated DEG signatures: genes Old-up were assigned +1 and Old-down −1 (duplicates removed). Region-specific “shared” signatures were built by intersecting WT and ReC significant genes within cortex and within cerebellum; a pan-regional signature was defined by the intersection of cortex-shared and cerebellum-shared sets. Signatures were scored per cell with VISION^40^: for stability, raw counts were copied to a plain assay and used to initialize a *Vision* object (no projection methods), followed by *analyze*; signature scores (SigScores) were mapped back to Seurat metadata. We further established two signatures to represent the age-related increase in interferon signaling and DAM genes. The genes used for the Interferon signature were: *Bst2*, *Ifi204*, *Ifi207*, *Ifi209*, *Ifi27l2a*, *Ifi30*, *Ifi47*, *Ifit3*, *Ifitm3*, *Irf7*, *Irf9*, *Oasl2*, *Stat1*, *Ly6e*. The genes used for the DAM signature were: *Cd22*, *Gpnmb*, *Spp1*, *Ccl3*, *Cst7*, *Ctsb*, *Lpl*, *Itgax*.

To quantify signature velocities (e.g. CAAS change with age), we modeled per-cell scores with a linear model that allowed for group-specific slopes. For WT microglia we fit score ∼ age + tissue + age:tissue; for reconstituted cells the same design was used. Age was treated as a numeric covariate in months. Slopes and 95% confidence intervals were obtained with *lstrends*^98^ (lsmeans/emmeans) on the fitted model, and pairwise slope differences were tested (Tukey’s range test) using the built-in contrasts. For figure panels (e.g., Fig. 3H), we visualized per-cell scores by age with violin/point overlays and added the fitted linear trend. As single-cell models treat cells as replicates, we additionally summarize scores by per-sample medians and perform two-tailed tests on those medians where appropriate (t-test or Wilcoxon, as indicated).

#### Publicly available bulk and scRNA-seq data analysis

We re-analyzed two previously published single-cell RNA-seq datasets: (1) Droplet-based scRNA-seq of freshly dissected SVZ at young and old age (n = 3 males per age; aged 3 and 28 months; all C57BL/6JN strain)^46^; (2) Smart-seq2-based scRNA-seq of freshly isolated cells from the myeloid and non-myeloid fraction of the striatum, cerebellum, hippocampus and cortex at young and old age (n = 4 males per age; aged 3 and 24 months; all C57BL/6JN strain)^49^. (3) Droplet-based single-nucleus RNA-seq of freshly dissected caudate putamen at young and old age (n = 2 males and n = 2 females per age; aged 3 and 21 months; all C57BL/6JN strain)^7^. (4) Region-resolved bulk RNA-seq of regions representing the spectrum of differing magnitudes of age-related transcriptomic changes in the brain (motor cortex, hippocampus, cerebellum, corpus callosum), from young to old age (males: n = 3-5, 3–28 months; females: n = 5, 3–21 months C57BL/6JN; all C57BL/6JN strain)^7^; (5) Droplet-based scRNA-seq of freshly dissociated grey and white matter (no cerebellar white matter) microglia and oligodendrocytes from 24 months old C57BL/6J or Rag1^-/-^ mice (n = 1-3 males per condition)^23^.

Single-cell and -nuc RNA-seq datasets from (1), (2), (3) and (4) have been previously embedded, annotated and analyzed^7^. Count matrices were subsequently normalized and scaled to allow for quantification and analysis of expression scores. For dataset (5), we obtained raw count matrices and sample annotation. Integration, annotation and analysis of microglia was performed using Harmony, with the same parameters described above in analysis of microglia and ReC scRNA-seq analysis.

We further reanalyzed a public, microarray dataset of bulk-collected primary microglia from 4, 12 and 22 months old mice, isolated from cortex, cerebellum, striatum and hippocampus (n=4 males per age group; all C57BL/6J strain; platform GPL11180)^19^. Raw data was analyzed in R using GEOquery and limma^99^. Expression matrices were retrieved as Series Matrix files, probe-level intensities were log2–transformed when required by the data distribution and normalized across arrays with normalizeBetweenArrays. Samples were grouped by brain region (Cerebellum, Cortex, Hippocampus, Striatum) and age (4, 12, 22 months) to form twelve groups (e.g., “Cerebellum_4months”, “Cortex_4months”, …). Probes with missing values were removed prior to modeling. We fit a linear model with one column per group (∼ 0 + group) and tested all directional pairwise contrasts between groups; statistics were moderated with empirical Bayes. Differential expression was called with Benjamini–Hochberg FDR < 0.05 using decideTests (no additional fold-change threshold).

#### Overlap of CAAS with Human microglia (HuMi) signature

We evaluated concordance between CAAS genes and a previously published set of age-related DEGs from human microglia (HuMi; from^50^; identified by comparing microglia data from humans of mean age = 53 years, and mean age = 94.07 years). Human symbols were converted to mouse-style capitalization (sentence case) and intersected with the CAAS set and plotted over a scatterplot comparing Log2FC of WT aging microglia (Old/Young) and heterochronic ReCs (Young→Old vs Young→Young). Genes falling into each concordance quadrant were counted. Enrichment of HuMi genes was assessed via quadrant-resolved overlap: age-up and age-down subsets from humans (HuMi_up, HuMi_down) were tested separately for enrichment among DEGs in each mouse quadrant: Q1 (up in WT & ReC), Q2 (up WT & down ReC), Q3 (down WT & ReC), Q4 (down WT & up ReC). In all tests, enrichment was computed with one-sided Fisher’s exact test using the expressed-gene intersection as background

#### Bulk-seq quantification and differential expression analysis

Bulk-seq data of ex vivo bone marrow-derived chimeric brain myeloid cells (Figure S9E-I), interferon-treated primary microglia in culture (Figure S12) and tissue punches of cortex and cerebellum in Rag2^-/-^γc^-/-^ mice were conducted as previously described^7^. In brief, raw sequence reads were trimmed to remove adaptor contamination and poor-quality reads using Trim Galore! (v.0.4.4, parameters: --paired --length 20 --phred33 --q 30). Trimmed sequences were aligned using STAR (v.2.5.3a, default parameters). Multi-mapped reads were filtered. Read quality control and counting were performed using SeqMonk v.1.48.0. Data visualization and analysis were performed using R scripts previously published^7, 100^. and the following Bioconductor packages: Deseq2^97^, and org.Mm.eg.db. Finally, we excluded pseudogenes and predicted genes from the count matrix to focus predominantly on well-annotated, protein-coding genes. In total, all of the following analyses were performed on the same set of 21,076 genes. In order to limit false negatives when intersecting DEG sets from different comparisons (i.e. intersect of DGE sets induced by IFN-β and IFN-γ) we considered DEGs significant if they passed the adjusted p < 0.1, as recommended by the original Deseq2 publication^97^.

#### Cell–cell communication analysis

Cell–cell signaling was inferred with CellChat^54^ using the mouse ligand–receptor database. We analyzed the WT scRNA-seq/CITE-seq dataset after QC and Harmony integration, using the cell_class annotation to define cell groups (microglia, border-associated macrophages, peripheral macrophages, granulocytes, T cells, NK cells, etc.). To assess aging effects, we split the dataset by age and created one CellChat object per group (Young and Old). For each object we ran the standard workflow of: subsetData → identifyOverExpressedGenes → identifyOverExpressedInteractions → computeCommunProb → computeCommunProbPathway → aggregateNet. Network visualizations and summaries were produced with CellChat’s plotting utilities. Incoming communication towards microglia was quantified from the aggregated networks. Figure 5A shows the number of incoming interactions by source cell type; Figure 5B summarizes pathway-level information flow (Young vs Old). For a direct group comparison, the two CellChat objects were merged and contrasted using compareInteractions (interaction counts and weights) and rankNet (stacked bar comparison of pathway contributions) with microglia as targets. We only examined interactions that were detectable in both young and aged mice. Example pathways (e.g., IFN-II axis) were displayed with netVisual_bubble and netVisual_aggregate.

To gauge the contribution of specific lymphocyte populations, we recomputed CellChat networks on (i) the full WT dataset, (ii) a dataset without T cells, and (iii) a dataset without NK cells (all grouped by cell_class and processed identically). We compared the set of active signaling pathways (those retained by CellChat after probability computation and pathway aggregation) between the full network and each dropout network.

#### CosMx integration, neighborhood clustering, and region annotation

CosMx single-cell transcriptomic sections were loaded into *Seurat*. When needed, section orientations were harmonized by sign-flipping slide coordinates. Sections were merged and the CosMx *QC flag* was used to remove low-quality cells. To enable spatial visualization we assigned the slide coordinates as 2-D embeddings.

For integration, the RNA assay was split by section, normalized with *SCTransform*. Samples were integrated with Harmony as embedded in Seurat (split by sample), followed by construction of an SNN graph, *Leiden* clustering (resolution = 0.4), and *UMAP* generation on the Harmony space. Marker genes were computed with *FindAllMarkers* to guide subsequent cell-class checks.

To capture mesoscale spatial context, we computed per-cell neighborhood composition vectors. Briefly, for each cell we queried neighbors within 100 µm using dbscan’s^101^ *frNN* function, tallied neighbor counts by coarse cluster, and row-wise z-scored these frequency vectors. The resulting matrix was reduced by principal component analysis and partitioned into k-means clusters to yield regional neighborhoods. These neighborhood clusters were then manually curated with interactive selections to assemble anatomically meaningful regions—e.g., cortex, cerebellum (including reassignment of cerebellar white matter), corpus callosum, hippocampus, striatum, thalamus, midbrain/pons/medulla, choroid plexus, ventricles, remaining white matter, and “other.” Final curated region labels were mapped back to the integrated object via stable cell indices for all sections and used for downstream visualization and analyses.

Count matrices were exported and annotated via the Allen Brain Atlas’ MapMyCells^88^ tool to classify major neuronal subtypes.

Given that CosMx data is not best suited for identification of subtle transcriptomic clusters, we utilized the deep-coverage and high-confidence labeling of microglia/ReC clusters to inform ReC states in CosMx data. To refine ReC states, we co-embedded CosMx immune cells (selected cells annotated as immune by MapMyCells (class probability ≥ 0.6) and/or expressing canonical immune markers *Ptprc* and *Tmem119*) with the single-cell ReC dataset (Figure 1K), using SCT normalization and Harmony integration across “data type” (CosMx vs. scRNA). We then performed manual label transfer from the ReC reference to CosMx cells. Immune states and cluster IDs were written back to the full spatial object and visualized in tissue coordinates.

#### Cerebellar layer assignment for reconstituted cells (CosMx)

To classify ReCs within the cerebellum into granule layer (GL), molecular layer (ML), or white matter (WM), we used Allen Brain Map “MapMyCells” annotations to define landmark neurons: Purkinje cells (class = “28 CB GABA”, subclass = “313 CBX Purkinje Gaba”) and granule neurons (class = “29 CB Glut”, subclass = “314 CB Granule Glut”). For each section independently, we subset cells within the cerebellum. We computed for reach ReC nearest-neighbor distances to the Purkinje and granule sets in the (x,y), and applied a 30 µm threshold: if neither landmark was within 30 µm, the ReC was labeled WM; if only Purkinje was within 30 µm, ML; if only granule was within 30 µm, GL; if both were within 30 µm, the closer landmark determined the label. When a landmark population was absent in a section, distances defaulted to ∞, preventing spurious assignments. Resulting labels were written back to the Seurat metadata as cerebellar_layer and visualized on spatial embeddings and raw XY plots for quality control.

**Spatial-seq CAAS calculation and cell proximity effects on CAAS**

CAAS scores were calculated per-cell as described for the scRNA-seq data. We quantified spatial coupling between ReCs and nearby cell types in CosMx data using a nearest-neighbor framework, following in principle previously described analyses^8, 12^. For neighbor categories we used MapMyCells subclass labels and, for immune cells, overrode subclass with the ReC annotation to ensure they appeared as distinct targets. The CAAS was normalized to [0,1] within each age×replicate stratum to reduce batch effects. For each ReC, we computed the minimum Euclidean distance to the nearest cell of every target type using KD-tree queries (RANN/nn2, k=1), yielding one distance per target type per cell. Target types were limited to sufficiently frequent subclasses (≥200 cells) plus immune categories as annotated before (especially for T cells). To visualize spatial relationships, we binned signature values as a function of distance with a sliding-window approach (window 30 µm, step 1 µm, up to 80 µm). Within each bin we calculated the mean and SEM of the normalized signature and then mean-centered each cell-type curve within an age stratum, so deviations reflect spatial modulation rather than global offsets. The pipeline outputs a per-cell distance table and binned curves (per age and cell type), and produces facet plots (mean±SEM) of signature vs. distance. This design tests whether the CAAS increases or decreases near specific partner cell types, while controlling for sample/age effects and enabling region-specific analyses.

#### Analysis of confocal microscopy data and microglia processes analysis

For colocalization analysis of EGFP+/Tmem119+/Iba1+ cells, Imaris (Oxford Instruments) was used to generate surface reconstructions for each marker, assign class identities, and quantify colocalization as the percentage of overlapping cells. For analysis of microglial processes, image processing and reconstruction were performed using Arivis Vision4D (Zeiss). A custom pipeline for process detection was developed, incorporating a deep learning model trained to reconstruct microglia from immunofluorescent images. Soma were detected using the Cellpose85 module, and processes were traced with the Neuron Tracer tool. Manual corrections were applied to ensure tracing accuracy. Resulting reconstructions were exported as SWC files and imported into Fiji/ImageJ, where the SNT plugin86 and SNT viewer were used for visualization and analysis. Sholl87 analysis was then performed, and measurements including branch length, convex hull roundness, and surface area were calculated and exported for further analysis. For additional visual representation, Imaris was used to generate surface renderings of microglia and NK cells, and images were exported as high-resolution TIFF files.

## ADDITIONAL RESOURCES

We created an web application app to provide interactive access to the scRNA-seq data of aged WT microglia and heterochronic ReCs: https://tinyurl.com/Microglial-Aging-Explorer.

## Supplementary table legends

**Table S1. Complete gene lists for microglia and ReC cluster marker analysis, regional differential gene expression results and functional enrichment analysis; related to Figures 1, S1-3.**

Each tab contains complete DEG lists and annotations for cluster marker genes, cortical-versus-cerebellar DEGs and GO results lists for functional enrichment analysis.

**Table S2. Complete age-related DEG lists for microglia, ReCs and Stat1^-/-^ ReCs, CAAS gene list, functional enrichment analysis and custom CosMx panel; related to Figures 3,4 and S5-9.**

Each tab contains complete DEG lists and GO results lists for functional enrichment analyses related to effects of aging in WT microglia, ReCs, bone-marrow derived reconstitution results and Stat1^-/-^ ReCs. Further contains the custom CosMx panel.

**Table S3. Complete age-related DEG lists for microglia in Rag2^-/^-γc^-/-^, Rag2^-/-^, Rag1^-/-^ and NK-depleted mice, and functional enrichment analysis; related to Figures 5,6 and S12.**

Each tab contains complete DEG lists and GO results lists for functional enrichment analyses related to effects of genetic and antibody-based lymphocyte depletions on aged microglia.

